# Multi-omics profiling reveals MAGEL2-driven defects in human corticogenesis shared across Prader-Willi and Schaaf-Yang syndromes

**DOI:** 10.64898/2026.05.01.722223

**Authors:** Jannis Buecking, Baran Enes Güler, Michael Eibl, Azmal Syed Ali, Tobias Walczuch, Tobias Beschauner, Susanne Theiss, Melanie Spanjaard, Katrin Hinderhofer, Freya Herrmann-Sim, Celine E. de Esch, Derek J.C. Tai, Michael E. Talkowski, Jeroen Krijgsveld, Christian P. Schaaf, Magdalena Laugsch

## Abstract

The human cortex acquires its advanced cognitive capacity through tightly regulated developmental programs, disruption of which underlies neurodevelopmental disorders such as Schaaf-Yang syndrome (SYS) and Prader-Willi syndrome (PWS). While SYS results from pathogenic variants in the imprinted gene *MAGEL2*, PWS arises from chromosomal deletions, imprinting defects or uniparental disomy encompassing the *MAGEL2* locus. However, the contribution of *MAGEL2* to disease pathogenesis and human corticogenesis is not fully understood. Here, we performed integrated transcriptomic, proteomic, and ubiquitinomic profiling of cortical neurons derived from CRISPR/Cas9-engineered isogenic human pluripotent stem cells (hiPSC) modeling SYS and PWS. Beyond PWS-specific signatures including dysregulated ribosomal processes, we identified *MAGEL2*-dependent defects shared across both disorders. These include reduced progenitor proliferation, accelerated neuronal maturation, impaired migration and adhesion, as well as abnormal synaptic development, collectively linking PWS and SYS at the level of cortical development. Notably, these phenotypes partially overlap with those observed in other neurodevelopmental disorders, suggesting that *MAGEL2* governs core pathways broadly vulnerable in disease. Together, our findings establish *MAGEL2* as a key regulator of human cortical development, provide a unifying mechanistic framework for SYS and PWS, accessible via a web-based platform.

## INTRODUCTION

The expanded size and cellular diversity of the human cortex support higher cognitive functions and depend on tightly regulated coordinated programs of neuronal differentiation and migration.^1–4^ Disruptions of these processes during development can result in neurodevelopmental disorders (NDDs), frequently manifesting as intellectual disability (ID) and autism spectrum disorder (ASD).^5–7^ These features are prominent in Schaaf-Yang Syndrome (SYS), caused by pathogenic variants in the maternally imprinted gene *MAGEL2*^8^, and in Prader-Willi Syndrome (PWS), caused by chromosomal deletions, imprinting defects or uniparental disomy of chromosome 15q11.2-q13, encompassing several imprinted protein-coding genes and non-coding RNAs (ncRNAs), including *MAGEL2.*^9,10^ Despite this shared genetic locus, the contribution of *MAGEL2* to disease pathogenesis in both disorders and its role in cortical development remain poorly understood.

Although both conditions share overlapping features, SYS patients exhibit more severe ID, higher rates of ASD, and increased neonatal mortality compared to PWS.^11,12^ Studies of small deletions within the PWS locus, including *MAGEL2*, have suggested that *MAGEL2* deletion alone is insufficient to cause PWS^13–15^, implicating a role of other genes or ncRNAs.^16–19^ However, we recently reported a patient with a deletion that includes *MAGEL2* but spares *SNORD*/*SNRPN*, while still exhibiting the full PWS phenotype^20^, raising the question of how MAGEL2 mechanistically contributes to PWS and SYS.

*MAGEL2* expression is mostly restricted to the nervous system and endocrine tissues.^21,22^ However, progress in understanding its function has been limited by the lack of reliable antibodies, making MAGEL2 a challenging protein to study. Results from non-neuronal cell lines overexpressing *MAGEL2* have implicated it in several key cellular processes, including endosomal trafficking^23–29^, RNA metabolism^30,31^, circadian rhythm^32,33^, and protein ubiquitination^23,34^. Other studies using hiPSC-derived neurons showed that *MAGEL2* plays an important role in neurite outgrowth, synapse development, and neuropeptide trafficking.^28,35–38^ However, the molecular mechanisms by which MAGEL2 governs these cellular processes, particularly in the context of human cortical neuronal differentiation, and how deletions and point mutations contribute to the pathophysiology of PWS and SYS, remain poorly understood.

Here, we performed an integrative multi-omics analysis of the transcriptome, proteome, and ubiquitinome in CRISPR/Cas9-engineered hiPSC-derived cortical neurons across six genotypes modeling SYS and PWS in two genetic backgrounds, including their isogenic controls. These data, validated by functional studies confirmed distinct PWS-specific dysregulated ribosomal processes, and uncovered shared defects between SYS and PWS, including decreased progenitor proliferation, accelerated neuronal maturation, impaired migration/adhesion, and abnormal synaptic development, mechanistically linking MAGEL2 to both disorders.

Overall, these findings position MAGEL2 as a key regulator of cortical neuron differentiation, with dysregulated pathways partially converging in hallmarks of other neurodevelopmental disorders (NDDs), thereby offering novel insights into the pathophysiology of SYS and PWS accessible through a web-based tool.

## RESULTS

### CRISPR/Cas9-engineered isogenic hiPSC lines with *MAGEL2* mutations, *MAGEL2* deletions, and PWS deletions for modeling SYS and PWS in cortical neurons

hiPSC lines carrying the two most common PWS deletions in the 15q11.2-13 region spanning 6 Mb (*PWS Type I*) and 5.3 Mb (*PWS Type II*), hereafter referred to as PWSTI and PWSTII, (Figure S1A), and their isogenic wild-type (WT) control hiPSC lines were previously generated via CRISPR/Cas9 in two distinct genetic backgrounds (male GM08330 (8330) and female MGH2069 (MGH)) and characterized in NGN2-induced neurons.^28^ As *MAGEL2* is only one of several genes deleted in PWS, we sought to specifically assess its contribution to the phenotype by generating heterozygous and homozygous full-gene *MAGEL2* deletions (HetDel and HomoDel) using CRISPR/Cas9 in the same two WT hiPSC lines. To distinguish between effects of *MAGEL2* deletions from those of point-mutations, we additionally generated the *c.1996dupC* (DupC) mutation (most common SYS mutation^11^), *c.1996delC* (DelC) mutation (perinatally lethal^12,39,40^) in both WT hiPSC lines (Figure 1A, S1A). This approach yielded a comprehensive panel of seven isogenic hiPSC lines per genetic background, comprising 14 hiPSC lines in total. These were rigorously validated by comprehensive molecular and epigenetic testing to confirm correct CRISPR/Cas9 targeting, imprinting and the paternal origin of the targeted allele (Figure S1B-F). Only clones that met all quality criteria were differentiated into cortical neurons for 30 days (d30) using a dual-SMAD inhibition protocol^41^ (Figure 1A). Hereafter, DelC, DupC, HetDel, HomoDel, PWSTI, and PWSTII, when compared with their respective isogenic WT controls, are collectively referred to as variants.

**Figure 1:**
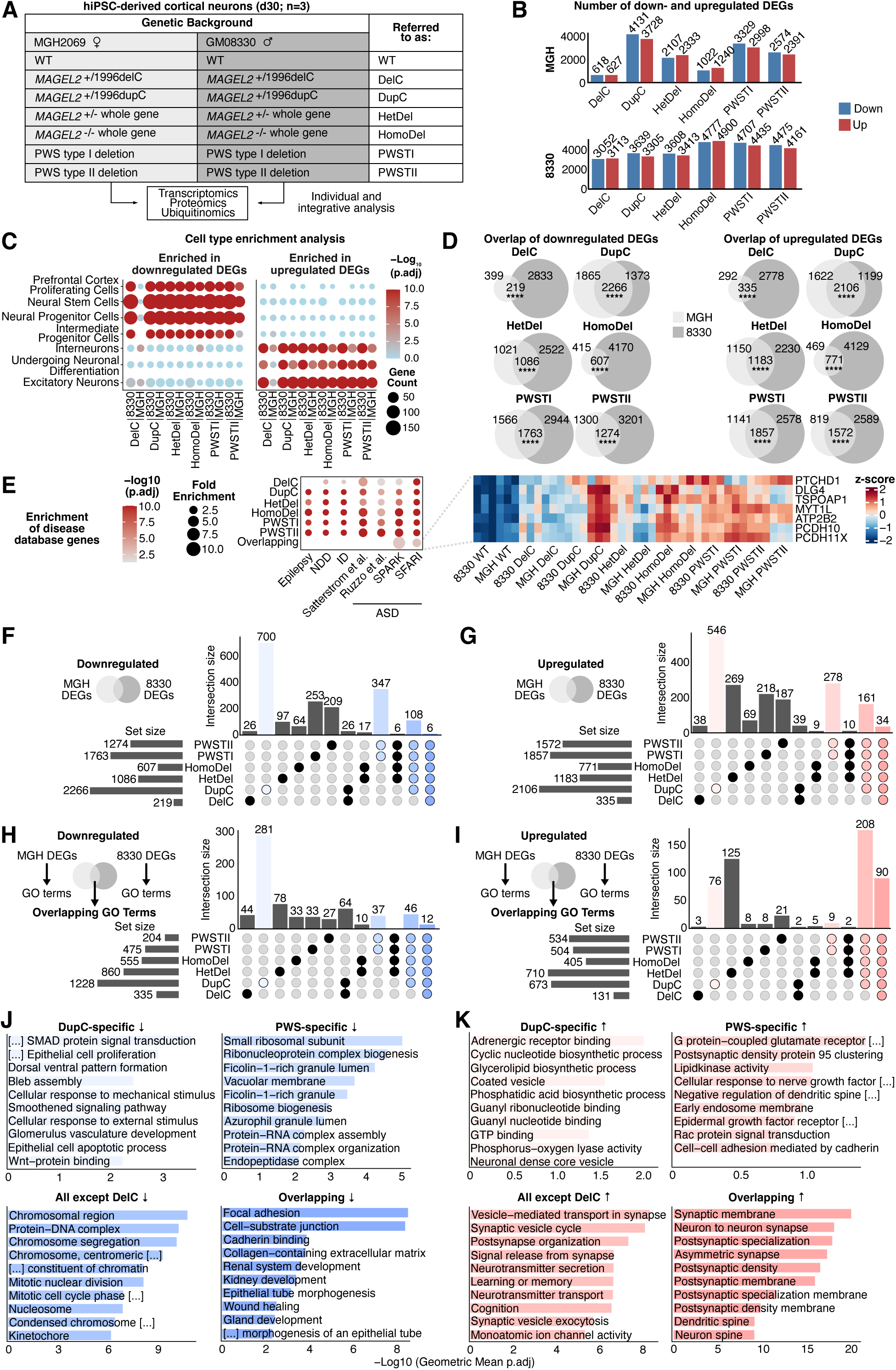
*MAGEL2* variants lead to broad transcriptional dysregulation in cortical neurons. (A) Overview of CRISPR/Cas9-engineered hiPSC lines in the 8330 and MGH genetic backgrounds. (B) Number of down- and upregulated DEGs in (FDR < 0.05) in hiPSC-derived neurons. (C) Cell type enrichment analysis using the MSigDB C8 database for down- and upregulated DEGs per variant and genetic background with significance determined by a hypergeometric test. (D) Overlap of down- and upregulated DEGs between both genetic backgrounds with significance of overlap determined with a hypergeometric test. (E) Enrichment of ASD^212,214,215,220^, ID^213^, epilepsy^213^, and NDD^213^ genes in background-overlapping DEGs, with significance determined with a hypergeometric test. Heatmap with z-score-normalized expression of significantly dysregulated SFARI genes shared across all genotypes. (F-G) UpSet plots of background-overlapping downregulated (F) and upregulated (G) DEGs. (H-I) UpSet plots of enriched GO terms for downregulated (H) and upregulated (I) DEGs. (J-K) Enriched GO terms of UpSet plots intersections (*p < 0.05, **p < 0.01, ***p < 0.001, ****p < 0.0001).

### *MAGEL2* variants lead to strong transcriptional dysregulation, including ASD-risk genes

While PWS deletions are known to cause widespread transcriptional deregulation of neuronal gene networks and developmental processes due to loss of small nucleolar RNAs (snoRNAs) such as *SNORD*/*SNRPN* transcripts^42,43^, we assumed that *MAGEL2* point mutations or whole-gene deletions would have a more restricted impact on gene expression. To test this hypothesis, we performed RNA-seq in cortical neurons derived from all 14 hiPSC lines modeling SYS and PWS (Figure S1G). Principal component analysis (PCA) of the RNA-seq profiles showed distinct clustering of all variants, highlighting variant-specific transcriptional dysregulation (PC2, 11.9%). The separation by genetic backgrounds (PC1, 20.3%), however, underscores the importance of using multiple genetic backgrounds for hiPSC-based studies (Figure S1H).

Contrary to our initial expectation, both SYS and PWS neurons displayed substantial transcriptional dysregulation. The number of differentially expressed genes (DEGs; FDR < 0.05) varied across variants and genetic backgrounds ranging from 618 downregulated genes in MGH DelC neurons to 4,900 upregulated genes in 8330 HomoDel neurons (Figure 1B, Supplementary Table 1). The expression levels of the *MAGEL2*-neighboring genes *NDN* and *MKRN3* (Figure S1A) remained unchanged in neurons with *MAGEL2* mutations or deletions and were absent in neurons with PWS deletions (Figure S1I, Supplementary Table 1), further supporting the specificity of our CRISPR/Cas9 engineering.

Previous studies suggested that deletion of the paternal *MAGEL2* allele, including its promoter, may exhibit aberrant “leaky” *MAGEL2* expression, driven by the promoter of the otherwise silenced maternal *MAGEL2* allele, potentially contributing to a milder phenotype in PWS compared to SYS.^44,45^ However, both RNA-seq and RT-qPCR data provide no evidence of such leaky *MAGEL2* expression in human cortical neurons, clearly demonstrating absence of *MAGEL2* in either homozygous or heterozygous *MAGEL2* and PWS deletions in both genetic backgrounds (Figure S1J, Supplementary Table 1). Thus, our data argue against compensatory expression of the maternal *MAGEL2* allele as a cause of the phenotypic differences between SYS and PWS. Notably, *MAGEL2* was consistently upregulated in DupC neurons, with a similar trend in DelC, suggesting a specific transcriptional regulatory context in SYS-related *MAGEL2* mutations, the mechanisms of which remain to be explored.

To gain an initial overview of the transcriptional dysregulation observed across all variants, we performed cell type enrichment analysis of the down- and upregulated DEGs for each variant in each genetic background (Figure 1C). This analysis revealed a consistent enrichment of proliferative progenitor signatures in downregulated DEGs (e.g., *PCNA*^46^ and *MCM2*^47^) and neuronal signatures in upregulated DEGs (e.g., *SCN2A*^48^, *SCN3A*^49^, and *GRIA2*^50^) across all variants and backgrounds except MGH DelC (Figure S1K). We next overlapped the DEGs from both backgrounds, resulting in background-overlaps ranging from 219 for downregulated DEGs in the DelC neurons to 2,266 DEGs in DupC neurons (Figure 1D). Querying these against available NDD, ASD, ID, and epilepsy gene databases revealed significant enrichment of associated genes, linking the observed transcriptional dysregulation to the high prevalence of these phenotypes in SYS and PWS (Figure 1E, Supplementary Table 2). The most pronounced enrichment (over 3.5-fold) was found in the shared DEGs across all variants significantly enriched for seven ASD genes, including the synaptic protein *DLG4* (PSD-95)^51,52^ and the neuronal transcription factor *MYT1L*^53–55^.

Overall, although MAGEL2 is not an established transcriptional regulator, both SYS and PWS cortical neurons show common dysregulation of gene expression affecting proliferative progenitor and neuronal signatures, as well as enrichment of genes associated with NDD, ASD, ID, and epilepsy, phenotypes observed in both syndromes, underscoring the biological relevance of these disease cell models.

### Variant-specific and shared dysregulation of transcriptomic pathways in SYS and PWS neurons

To delineate variant-specific and shared transcriptomic changes across all variants, we generated UpSet plots that revealed extensive variant-specific and shared set intersections. For example, among downregulated DEGs, 700 were DupC-specific, 17 exclusively shared between DelC and DupC, 347 PWS-specific, and 6 shared across all variants (Figure 1F-G, Supplementary Table 3). We hypothesized that variability in the number of DEGs reflects distinct gene sets that contribute to related biological processes. Therefore, gene ontology (GO) enrichment analyses were first performed separately for each variant-to-WT comparison and background before comparing the significantly enriched GO terms across backgrounds to identify overlapping biological pathways. This analysis yielded large numbers of significantly enriched GO terms (FDR < 0.05), ranging e.g., for downregulated DEGs from 615 in MGH DelC neurons to 1,949 in 8330 HomoDel neurons (Figure S2A, Supplementary Table 4). UpSet plots of these GO terms (Figure S2B) identified both specific and shared GO terms (Figure 1H-I, Supplementary Table 5) with the most significantly enriched terms shown in Figure 1J-K.

As for downregulated DEGs, DupC-specific GO terms showed dysregulation of TGF-β signaling mediated by the SMAD transcription factor family, EGF signaling, and cell polarity, pathways that may relate to the more severe phenotype observed in patients with this variant. Consistent with prior reports for PWS neurons^42,43,56^, the 37 GO terms specifically shared between PWSTI and PWSTII were linked to translational and ribosomal processes. Their absence in neurons with sole *MAGEL2* deletions and mutations (Figure 1H) indicates key molecular differences between SYS and PWS at the transcriptional level. The PWSTI deletion (Figure S1A) spans 208 coding and non-coding genes^57^, of which 55 (out of 347; 16%) were significantly downregulated specifically in PWS neurons. As the enrichment for ribosome and translation-related GO terms remained after excluding the PWS locus genes (Figure S2C-E, Supplementary Table 6), we speculate that these altered pathways may arise from dysregulation of snoRNA targets. In addition, PWS neurons showed a shared dysregulation of small ncRNAs such as the *RNU* and *RNY* genes involved in RNA-splicing (Figure S2F, Supplementary Table 1), including *RNU4-2*, recently implicated in approximately 0.4% of NDDs.^58,59^ These findings suggest a partially shared pathophysiology of PWS with other splicing-related NDDs, consistent with the known role of SNORD RNAs in splicing.^60–62^ Beyond the DupC- and PWS-specific findings, 46 shared GO terms across all variants except DelC were associated with proliferation, including *PCNA*^63^ and *MCM2*^64^, while 12 GO terms shared across all variants were linked to wound-healing and substrate-related adhesion including *NCAM1*^65^, *VIM*^66^, and *SWAP70*^67^ (Figure 1J, S1K). These data support the prior observation of cell-type enrichment across PWS and SYS (Figure 1C), indicating that these altered pathways contribute to a shift from proliferative progenitor toward more differentiated neuronal signatures.

For upregulated genes, we found 76 enriched DupC-specific GO terms mostly related to metabolic processes, as well as 79 PWS-specific terms widely linked to synaptic receptors (Figure 1K). The 208 terms common to all variants except DelC and the 90 terms overlapping across all variants were primarily related to nervous system and synapse development, including *MYCBP2*^68^, *GRIA2*^50^, *SCN2A*^69^, and *BSN*^70^ (Figure 1K, S1K), further supporting the cell type enrichment results (Figure 1C).

Despite variant-specific differences, including PWS-specific impact on RNA and ribosome biology, all variants exhibited shared dysregulation of adhesion/cytoskeletal pathways in downregulated DEGs and synaptic development in upregulated genes at the transcriptomic level, confirming that distinct DEG sets contribute to similar biological processes across PWS and SYS neurons.

### *MAGEL2* variants disrupt protein homeostasis

Given the extensive transcriptional changes observed across SYS and PWS neurons, as well as the known role of MAGEL2 in protein homeostasis^28,31,33,34^, we next asked whether these altered transcriptional programs are associated with proteomic changes. For this purpose, we performed label-free quantitative mass spectrometry in cortical neurons derived from all 14 hiPSC lines modeling SYS and PWS. This resulted in high coverage, identifying a total of 9,153 proteins, with 6,346 proteins detected across all samples (Figure S3A). PCA separated the samples by genotype (PC1, 27.6 %), with weaker clustering by genetic background (PC2, 17.7%; Figure S3B). The number of differentially expressed proteins (DEPs, FDR < 0.05) was high for each genotype and background, indicating strong proteomic changes in both SYS and PWS neurons, ranging from 3,504 downregulated proteins in 8330 DupC neurons to 2,459 upregulated proteins in MGH DupC neurons (Figure 2A, Supplementary Table 7). Moreover, the numbers remained high in background-overlaps, ranging from 2,011 for downregulated DEPs in the PWSTII neurons to 1,368 for upregulated DEPs in DupC neurons (Figure 2B) further confirming strong proteomic dysregulation across all variants irrespective of genetic background.

**Figure 2:**
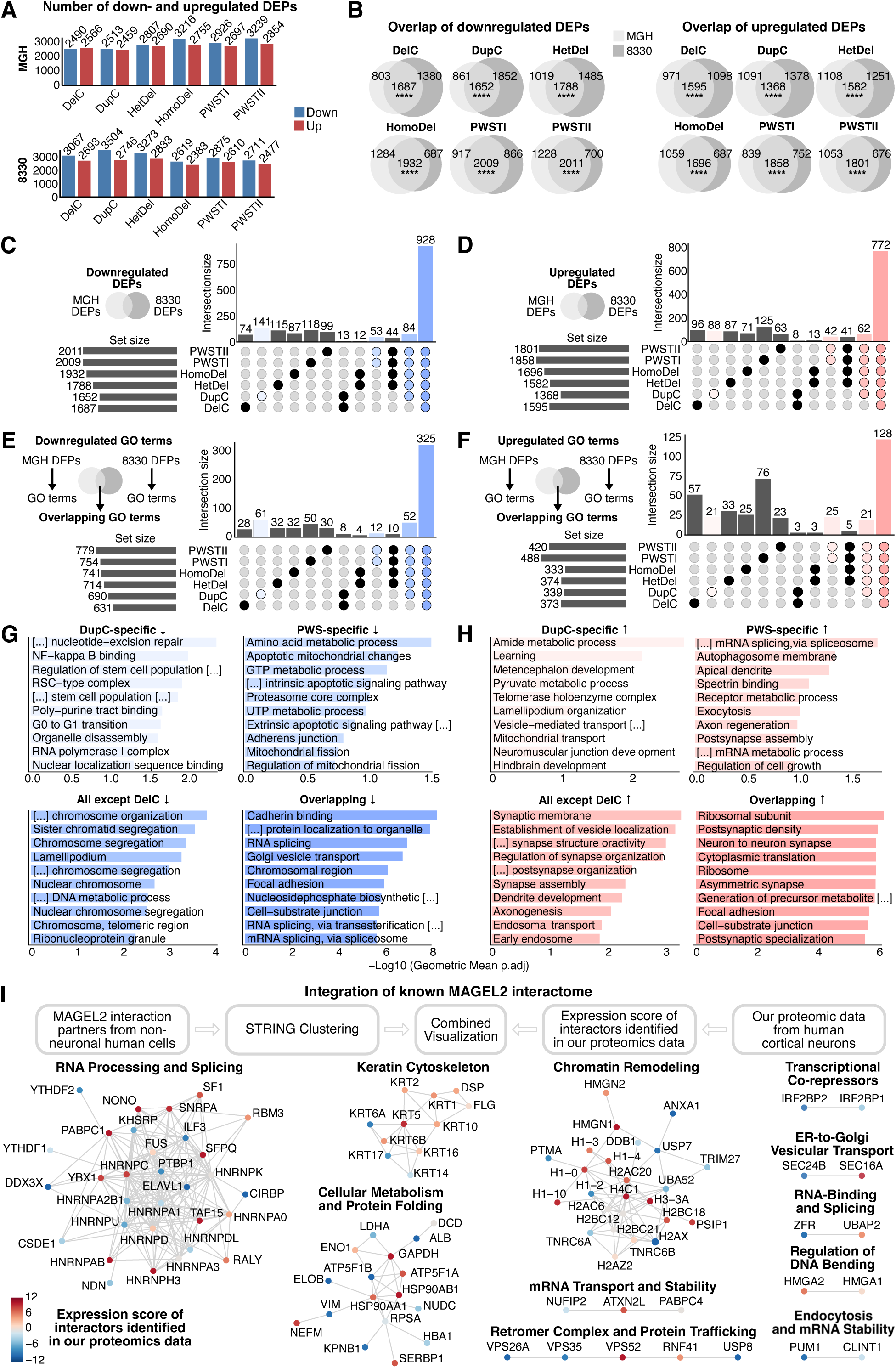
Proteomics data underscore the role of MAGEL2 in protein homeostasis, vesicle trafficking and RNA metabolism. (A) Numbers of down- and upregulated DEPs (FDR < 0.05). (B) Background-overlaps for each variant in down- and upregulated DEPs with significance of overlap determined by a hypergeometric test. (C-D) UpSet plots of background-overlapping downregulated (C) and upregulated (D) DEPs. (E-F) UpSet plots of enriched GO terms for downregulated (E) and upregulated (F) DEPs. (G-H) Enriched GO terms of UpSet plots intersections. (I) STRING PPI network of the MAGEL2 interactome performed using a total of 103 proteins extracted from publicly available data^23,30,31,34^ resulting in 11 sub-networks. Nodes represent proteins in functional communities. Node colors: cumulative "Expression Score" integrating expression of interactors in our proteomics data across all variants (red: shared upregulation, blue: shared downregulation across all variants).

To assess the clinical phenotypes associated with this proteomic dysregulation, we performed human phenotype ontology enrichment (HPO) analysis on the down- and upregulated DEPs shared across variants. Enriched terms included neurodevelopmental delay, abnormal forebrain morphology, hypotonia and abnormal cerebral morphology, thus recapitulating common PWS and SYS phenotypes (Figure S3C, Supplementary Table 8). UpSet plots revealed extensive overlap of DEPs across all variants and genetic backgrounds, with 928 downregulated and 772 upregulated proteins (Fig. 2C-D, Supplementary Table 9), indicating broad and consistent dysregulation shared between SYS and PWS neurons. To uncover unique and shared biological pathways across the variants, analogous to the transcriptomic analysis, we performed GO enrichment analyses for up- and downregulated DEPs separately for each variant-to-WT comparison and genetic background (FDR < 0.05). This yielded extensive lists of enriched GO terms, ranging from 1,286 terms for downregulated proteins in DupC neurons and 542 terms for upregulated proteins in HomoDel neurons (both in 8330 genetic background) (Figure S3E, Supplementary Table 10). We then overlapped enriched terms across genetic backgrounds (Figure S3F) and visualized these intersections using UpSet plots, (Figure 2E–H, Supplementary Table 11).

For downregulated DEPs, 61 DupC-specific terms were primarily related to transcriptional regulation and other DNA associated processes (Figure 2G). In contrast, the 12 PWS-specific terms were enriched in mitochondrial and proteostasis-related processes, whereas 52 terms shared by all variants except DelC were enriched for proliferation-associated terms (e.g., MSI1^71^ and MCM6^72^; Figure S3F). A large set of 325 GO terms common to all variants was enriched for vesicle-mediated transport (e.g., RAB5C^73^), RNA-splicing (e.g., SNRPD1^74^), and cellular metabolism (e.g., SDHA^75^), indicating broad convergence of proteomic changes in these biological pathways.

Upregulated DEPs were associated with 21 DupC-specific GO terms mainly related to metabolic pathways and developmental processes, while 25 PWS-specific terms were linked to splicing and endosomal functions (Figure 2H). In addition, 21 terms shared by all variants except DelC were associated with synaptic organization and signaling (e.g., BSN^70^ and PSD-95^51,52^), and 128 GO terms shared across all variants were related to ribosome biology (e.g., RPL7^76^).

### MAGEL2 interactome provides a link to shared dysregulation of vesicle trafficking, RNA metabolism, and metabolic networks in SYS and PWS neurons

The lack of validated anti-MAGEL2 antibodies suitable for direct pull-down of endogenous MAGEL2 limits our ability to determine its direct interaction partners. Hence, to relate the identified proteomic dysregulation in SYS and PWS neurons to MAGEL2 function, we examined whether and which known MAGEL2 interaction partners are differentially abundant in our proteomics data. We first listed all reported MAGEL2-interacting proteins from four independent studies identified by pull-down and proximity labeling, all of which were obtained after MAGEL2 overexpression in non-neuronal human cell lines.^23,30,31,34^ These interactors were visualized in UpSet plots revealing low overlap between the datasets, indicating that the MAGEL2 interactome varies substantially across experimental conditions (10 datasets, Figure S3G, Supplementary Table 12). Therefore, we focused on 130 WT MAGEL2 interactors, which were mapped onto a protein-protein interaction (PPI) network that included 103 proteins with at least one documented protein interaction. Finally, we integrated our proteomics data by deriving a cumulative "Expression Score" for each interactor, which combines the expression across variants (see Methods) (Figure 2I, Supplementary Table 13).

Eleven functional sub-networks were identified, several of which contained multiple down- and upregulated DEPs shared across variants (Figure 2I). In the vesicular transport and retromer-associated subnetworks, for example, we observed consistent downregulation of VPS26A and the deubiquitinase USP8, together with upregulation of VPS52, indicating that known components of the MAGEL2 trafficking machinery^25,34^ are commonly perturbed in SYS and PWS neurons. Moreover, proteins involved in RNA processing, splicing, and mRNA turnover, including members of the HNRNP and YTHDF families, also showed both down- and upregulated DEPs, strengthening prior evidence from human non-neuronal cells implicating MAGEL2 in the regulation of RNA metabolism.^23,30,31,34^ As MAGEL2 itself contains a PABP-1234 (C-terminal poly(A)-binding) domain^31^, it may bind RNA directly, pending further study.

Consistent with our previous finding, showing enriched GO terms associated with metabolism in down- and upregulated DEPs (Figure 2G-H), the interactome also revealed sub-networks linked to cellular metabolism (Figure 2I), several of which were also differentially expressed, including GAPDH and other proteins commonly used for normalization in experimental protein quantification. While their transcript levels were comparatively more stable, the proteomic data argues against their use as internal protein loading controls for studies of SYS and PWS.

Overall, with this correlative approach, we revealed a dysregulated MAGEL2 interactome associated with vesicle trafficking, RNA metabolism, and metabolic pathways, potentially contributing to the shared SYS/PWS phenotypes observed in our cellular models.

### Ubiquitinome profiling reveals shared dysregulation of metabolism and cytoskeletal pathways

MAGEL2 is a component of the MAGEL2-USP7-TRIM27 (MUST) E3 ubiquitin ligase complex, which primarily regulates K63-linked ubiquitin signaling rather than canonical proteasomal degradation^23,25,26,28,77^, suggesting that *MAGEL2* mutation variants may perturb ubiquitin-dependent regulatory pathways. Hence, we profiled the ubiquitinome across WT and SYS/PWS cortical neurons. Due to technical limitations, the 8330 HetDel line was not included. After pull-down for ubiquitinated proteins, label-free quantitative mass spectrometry identified 5,278 ubiquitinated proteins in total, of which 3,572 were detected across all samples (Figure S4A). PCA of the ubiquitinome separated samples primarily by genetic background (PC1, 24.5%) and less by variant (PC2, 12%; Figure S4B). The number of differentially ubiquitinated proteins (DEUs; FDR < 0.1) varied by variant and genetic background, ranging from 298 upregulated DEUs (8330 DupC) to 1,545 downregulated DEUs (MGH HomoDel; Figure 3A). Across backgrounds, overlapping DEUs ranged from 73 upregulated DEUs for DupC to 738 upregulated DEUs for HomoDel (Figure 3B, Supplementary Table 14).

**Figure 3:**
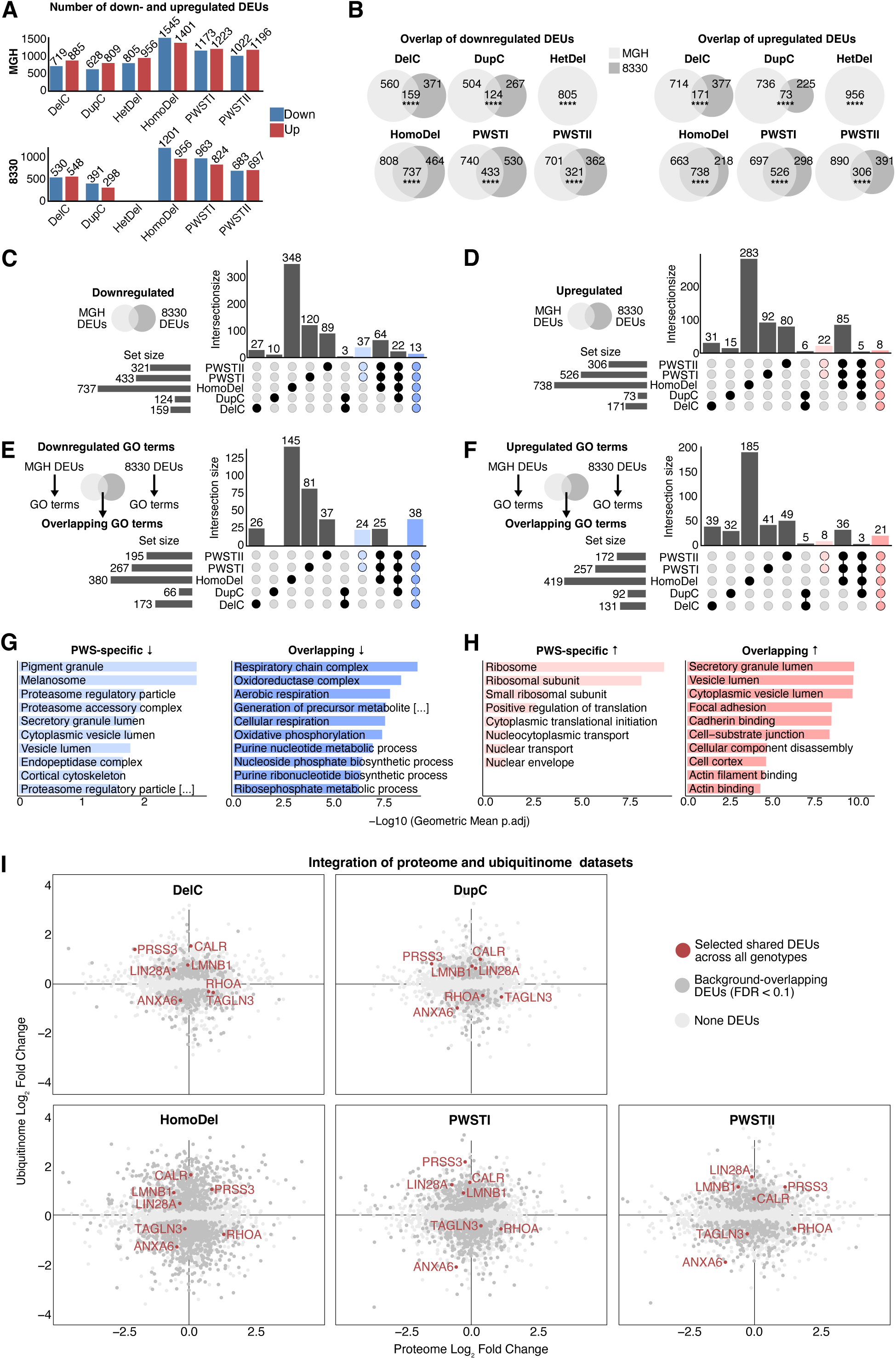
Variants in *MAGEL2* disrupt the neuronal ubiquitinome. (A) Number of up- and downregulated DEUs (FDR < 0.1). (B) Background-overlaps for each variant in down- and upregulated DEUs with significance of overlap determined by a hypergeometric test. (C-D) UpSet plots of background-overlapping downregulated (C) and upregulated (D) DEUs. (E-F) UpSet plots of enriched GO terms for downregulated (E) and upregulated (F) DEUs. (G-H) Enriched GO terms of UpSet plots intersections. (I) Scatter plot showing the log2 fold change of proteins in the proteome (x-axis) versus the ubiquitinome (y-axis).

We then examined distinct versus shared DEUs across genotypes visualized by UpSet plots, identifying 13 downregulated and 8 upregulated DEUs shared across all variants (Figure 3C-D, Supplementary Table 15). To assess whether distinct DEU sets contribute to similar biological processes shared between PWS and SYS neurons, we first performed GO enrichment analyses separately for down- and upregulated DEUs in each variant and background (Figure S4C, Supplementary Table 16). We then identified overlapping GO terms between backgrounds (Figure S4D, Supplementary Table 16) to determine variant-specific and shared terms (Figure 3E-H, Supplementary Table 17). No DupC-exclusive GO terms were enriched among downregulated DEUs, unlike the high numbers previously shown for downregulated DEGs (281 GO terms; Figure 1H-I) and DEPs (61 GO terms; Figure 2E-F). The 38 GO terms shared by all variants were primarily associated with mitochondrial processes and (ribonucleotide) metabolism (Figure 3G, Supplementary Table 17). GO terms associated with upregulated DEUs included eight PWS-exclusive terms related to translation and ribosome as well as spliceosome function (Figure 3H, S4E). Despite the identification of only 21 shared GO terms with affected ubiquitination targets across all variants and backgrounds (Figure 3H, Supplementary Table 17), these terms were enriched for actin cytoskeleton organization, adhesion, and cell-substrate junctions, indicating that SYS and PWS neurons share phenotypes consistent with changes previously observed at the transcriptome and proteome levels, supporting a MAGEL2-specific role in a defined subset of ubiquitin-dependent regulatory pathways.

### Integrating the proteome and ubiquitinome identifies novel potential ubiquitination targets

To contextualize the ubiquitinome data within the overall protein abundance, we integrated data from both modalities. A global comparison of the stoichiometry ratios of ubiquitinated to total protein abundance changes for all 3,572 overlapping proteins revealed similar genome-wide distributions across all variants (Figure S4F, Supplementary Table 18). Thus, our data suggest that MAGEL2 does not contribute to a global shift in protein ubiquitination but rather affects the ubiquitination of a subset of proteins. Hence, for each variant, we plotted the 4,553 proteins (light grey) detected in both proteome and ubiquitinome datasets and selected all background-overlapping DEUs (dark grey). Among these, we identified 22 shared DEUs across variants, of which seven are highlighted in red (Figure 3I, Supplementary Table 15). LIN28A, an RNA-binding protein (RBP) that promotes neural progenitors proliferation and inhibits their differentiation^78,79^ and TAGLN3, an actin-binding protein involved in synaptic organization^80^, showed significant changes in their ubiquitination status without corresponding changes in their total protein levels, indicating a putative non-degradative role for ubiquitination of these targets. In contrast, RHOA, a master regulator of the actin cytoskeleton, neuronal migration, and proliferation^81–85^, showed elevated expression levels across all variants, while its ubiquitination was decreased, indicating RHOA stabilization possibly through reduced ubiquitin-mediated targeting for proteasomal degradation, which may ultimately contribute to the dysregulation of actin dynamics. While MAGEL2 acts within the MUST complex, which primarily regulates K63-linked ubiquitin signaling^23,25,26,28,77^, our pull-down captured both proteasomal and non-proteolytic ubiquitin signaling events, potentially masking a direct MAGEL2-driven targeting. Nevertheless, the low number of ubiquitination-altered proteins is enriched among neurodevelopmental and cytoskeletal proteins, supporting the transcriptomic and proteomic data. However, in contrast to the strong dysregulation in the proteome, these findings suggest that most SYS/PWS-related phenotypes arise primarily from broader proteome-level changes, and only a subset of dysregulated proteins is directly associated with MAGEL2-related ubiquitination.

### Integrating the transcriptome, proteome and ubiquitinome reveals common dysregulation of cell adhesion pathways

To further evaluate the idea of distinct proteomic signatures contributing to the overlapping phenotypes in SYS and PWS, we compared GO terms enriched in the transcriptome, proteome, and ubiquitinome datasets across background-overlapping variants (Figure 4A-B, Supplementary Table 19). Among downregulated DEGs/DEPs/DEUs, only seven transcriptome-specific GO terms related to extracellular matrix organization and general developmental processes were identified (Figure 4A), suggesting a relatively limited impact of the transcriptome on the shared phenotypes between SYS and PWS. In contrast and consistent with our hypothesis, the 94 proteome-specific GO terms enriched in endosomal transport, organelle organization, and metabolic activity underscore the idea that the majority of phenotypes may arise from changes in the proteome, largely independent of transcription or ubiquitination. The 22 terms overlapping between the proteome and the ubiquitinome, linked to metabolism, indicate that MAGEL2-related ubiquitination may selectively tune metabolic pathways at the protein level. Finally, three pathways shared across all omics layers were identified, related to adhesion and migration, suggesting that these processes represent core mechanisms underlying altered neuronal connectivity in SYS and PWS cortical neurons.

**Figure 4:**
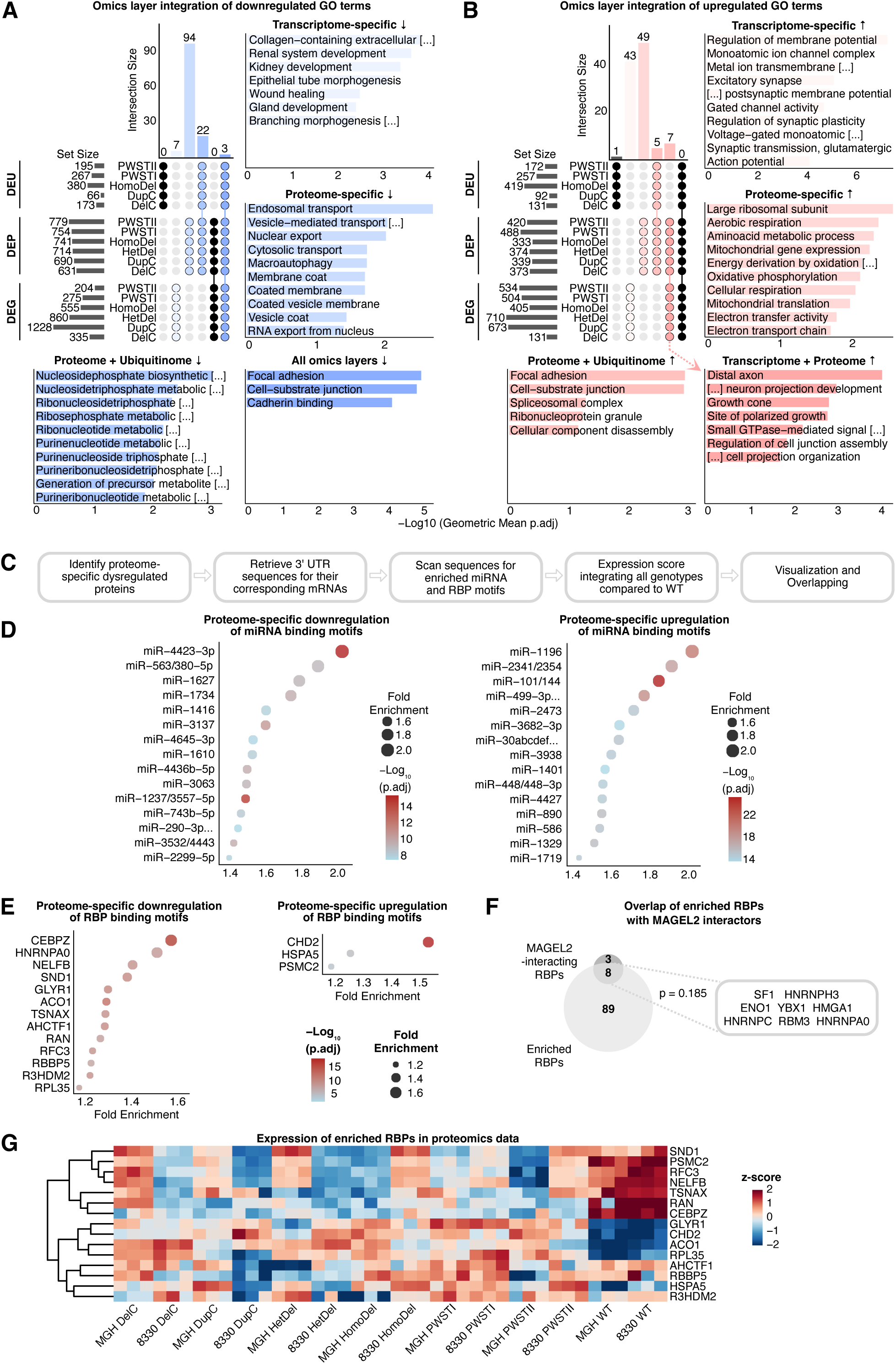
Integrating the transcriptome, proteome and ubiquitinome reveals shared dysregulation of cell adhesion pathways and suggest an important post-transcriptional role of MAGEL2. (A, B) Upset plots integrating enriched GO terms from the transcriptome, proteome, and ubiquitinome for downregulated (A) and upregulated (B) features with enriched GO terms of UpSet plots intersections. (C) Workflow for RBP and miRNA enrichment. (D) Dot plot showing miRNAs whose motifs were enriched in the transcripts of proteins specifically dysregulated on the protein level. (E) Dot plot showing RBPs whose motifs were enriched in the transcripts of proteins specifically dysregulated on the protein level. For (D-E), the statistical significance was determined using a one-sided Fisher’s exact test. (F) Venn diagrams showing the overlap between enriched RBPs and known MAGEL2 interactors with significance determined by a hypergeometric test. (G) Heatmap of proteomics expression of the top enriched RBPs.

For upregulated DEGs/DEPs/DEUs (Figure 4B), 43 transcriptome-specific GO terms emerged, primarily related to neuronal differentiation and synapse development. The majority of the 49 proteome-specific GO terms were linked to metabolic pathways. Seven GO terms shared between the transcriptome and proteome were associated with neuron projections and polarized growth, while five terms shared between the proteome and ubiquitinome were linked to focal adhesion pathways. Unlike the downregulated DEGs/DEPs/DEUs, no GO terms were common across all three omics layers, indicating that the upregulated changes are less uniformly coordinated across all layers.

Taken together, these findings support the view that distinct MAGEL2-dependent proteomic signatures, particularly in cytoskeletal- and adhesion-related pathways, drive the shared SYS and PWS phenotypes, with transcription and ubiquitination acting in a more selective and layer-specific manner.

### Post-transcriptional dysregulation driven by MAGEL2 is linked to adhesion and migration processes

Given that the MAGEL2-related ubiquitination does not primarily regulate protein abundance in SYS and PWS neurons, we postulated that the discordance between transcript and protein levels may indicate changes in mRNA stability, translational efficiency, or protein turnover.^86–88^ Hence, we performed a global comparison of protein-transcript ratios for each variant to select proteome-specific DEPs without a corresponding change in transcriptomic data (Figure S4G, Supplementary Table 20). As miRNAs and RBPs serve as important post-transcriptional regulators that fine-tune protein abundance, we sought to investigate their potential motif enrichment in the transcripts of proteome-specific DEPs (Figure 4C). For each variant, we selected the 8,732 proteins detected in both proteome and transcriptome datasets and identified 725 proteome-specific background and variant-overlapping DEPs (down 362, up 363 DEPs, Supplementary Table 21). Motif enrichment analysis on 3’ UTRs of all proteome-specific DEPs identified 194 miRNA binding motifs in the downregulated DEPs and 512 miRNAs in the upregulated DEPs (Figure 4D; Supplementary Table 22). These miRNAs include regulators of neurogenesis (miR-30/145/101 families), neural progenitor (NP) proliferation (miR-181 family), and neuronal maturation (miR-132/212 family).^89–94^ Of note, while several of these miRNAs are directly linked to NDDs and ASD^95–99^, they remain predictive and require further direct validation.

RBP motif enrichment identified 97 RBP motifs among the 362 proteome-specific, downregulated DEPs, but only three (CHD2, HSPA5, and PSMC2) among the 363 proteome-specific upregulated DEPs (Figure 4E, Supplementary Table 23). Overrepresentation of RBP motifs specifically in downregulated proteins suggests a preferential involvement of RBP-dependent repression mechanisms in neurons with *MAGEL2* mutations and deletions. RBP motifs of the upregulated proteome-specific DEPs exhibited longer motif length with higher sequence specificity and increased GC content compared to downregulated DEPs, suggesting MAGEL2-associated structural specificity influencing the affinity and/or stability of RBP-mRNA interactions (Figure S4H).

The overlap between the motif-identified RBPs and the known MAGEL2 interactors identified eight proteins, including YBX1, essential for NP proliferation and neurogenesis timing^100–102^, the key splicing factor SF1^103^; and RBM3, important for neuronal maturation^104–106^ (Figure 4F). We next examined whether these enriched RBPs showed proteomic dysregulation. Several of the most highly enriched RBPs were indeed dysregulated on the protein level (Figure 4G), including CHD2 (motifs enriched in both up- and downregulated proteome-specific DEPs), linked to NDDs, epileptic encephalopathy, intellectual disability, and autism.^107,108^

The miRNA/RBP analysis predicts how MAGEL2 may orchestrate post-transcriptional regulation of cortical neuronal differentiation, supporting shared changes in transcriptome and proteome signatures that converge on adhesion/migration defects in SYS and PWS neurons.

### Factor analysis identifies core disease signatures linked to the clinical SYS and PWS phenotypes

To complement the differential expression and overlap-based analyses, we applied Multi-Omics Factor Analysis 2 (MOFA2)^109,110^, an unsupervised integration approach that identifies shared, latent axes of variation across modalities without relying on significance thresholds or predefined gene sets. MOFA2 decomposed the variation across all three omics layers into 15 factors, capturing ∼58% of variance in the transcriptome, ∼72% in the proteome, and ∼63% in the ubiquitinome (Figure 5A-B, S5A-B). Since Factor 1 separated all variants from the WT controls (Figure 5C), it represents a core, background-overlapping molecular signature shared by all variants. Moreover, Factor 1 captured substantially more variance for the proteome (∼30%) than the transcriptome (∼5%) or ubiquitinome (∼9%), underscoring that the proteome is the molecular layer most affected by both *MAGEL2* mutations and deletions (Figure 5A). Since the remaining factors largely reflected background- or genotype-specific variation, we prioritized Factor 1 for downstream interpretation. As shown in the heatmaps, top-weighted features for Factor 1 include *DCX* linked to neuronal migration^111–113^ and *SEZ6* involved in synaptic plasticity^114,115^ in the transcriptome, ribosomal proteins such as RPL13, RPL39 and RPL7^116^ in the proteome, and the cytoskeleton regulator RHOA^81–85^ in the ubiquitinome (Figure S5C-E).

**Figure 5:**
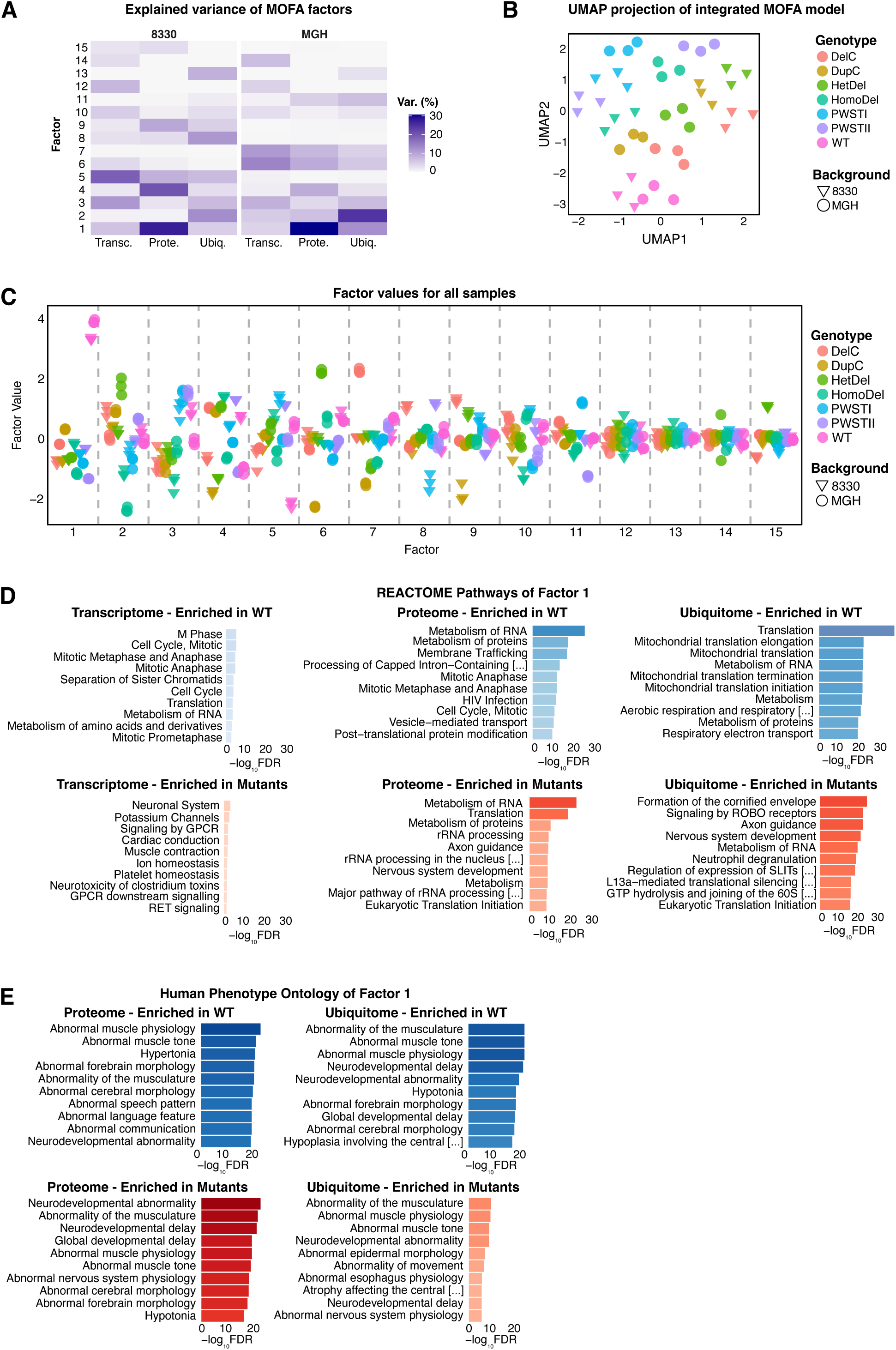
MOFA integration uncovers core disease signatures. (A) Variance decomposition by each of the 15 identified factors across the different omics layers. (B) UMAP projection of the integrated MOFA model with samples colored by genotype, shaped by genetic background (n=3). (C) Factor values for all samples colored by genotype and shaped by genetic background. (D, E) Bar plot showing the significant enrichment of REACTOME (D) and HPO (E) terms for Factor 1. For both (D) and (E), significance was determined using g:Profiler2 (FDR < 0.05).

Using gene set enrichment analysis on feature weights from each omics layer, we identified REACTOME pathways significantly associated with Factor 1 (Figure 5D, Supplementary Table 24). Positive weights (towards the WT) were enriched in proliferation-related programs in both the transcriptome and proteome, in addition to mitochondrial pathways in the ubiquitinome. In contrast, negative weights (towards all *MAGEL2* variants) were enriched for neurogenesis and axon guidance in the transcriptome and proteome, whereas general nervous system development terms were enriched in the ubiquitinome. Gene sets related to RNA metabolism and translation were enriched among both positively and negatively weighted features, suggesting simultaneous increases and decreases across different genes within the same functional modules. The clinical relevance of these findings was assessed by HPO enrichment of the Factor 1 weights (Figure 5E, Supplementary Table 25). While no significant enrichment was found for the transcriptome features, proteome and ubiquitinome features were strongly enriched for terms matching the clinical spectrum of PWS and SYS, including neurodevelopmental delay, neuromuscular phenotypes, abnormalities of brain morphologies, and speech delay.^9–11,117^

MOFA2 identified a major factor separating all *MAGEL2*-variants from WT samples, capturing a shared, proteome-driven disease signature enriched for proliferation, translation, and neurodevelopmental pathways, consistent with our hypothesis of proteome-dominant dysregulation underlying the SYS and PWS phenotype.

### Shared early cytoskeletal and proliferative deficits in SYS and PWS neurons

Our multi-omics data show that hiPSC-derived cortical neurons carrying *MAGEL2* mutations and deletions recapitulate key PWS and SYS characteristics and revealed aberrant migration, adhesion, and proliferation emerging as one of the most striking molecular signatures shared across all six variants. To validate these data, we chose the MGH background, which generally showed milder changes compared to 8330, thereby providing a more rigorous test of our findings. Since focal adhesion kinase (FAK) autophosphorylation at Y397 is a central event regulating focal adhesion turnover and cytoskeletal organization^118,119^, we performed Western blotting to semi-quantitatively assess pFAK/FAK ratios, which were reduced across all variants (Figure 6A, S5F). We hypothesized that early cytoskeletal and adhesion changes could precede the altered adhesion and migration phenotype identified by multi-omics analysis at d30 in SYS and PWS neurons. Hence, subsequent analyses of migration, adhesion and proliferation were performed at an intermediate stage between d8 and d19 (depending on the assay), comprising a mixed population of dividing neural progenitors and immature neurons. Using a wound-healing assay and live-cell imaging (d12 to d17), we revealed significantly reduced wound closure in all variants, with the DupC neurons displaying the most severe phenotype (Figure 6B-C). Consistent with the dysregulation of migration-associated genes and proteins (Figure S1K, S3C), single-cell tracking showed that all variants migrated more slowly, covering less distance with reduced average speed and total displacement.

**Figure 6:**
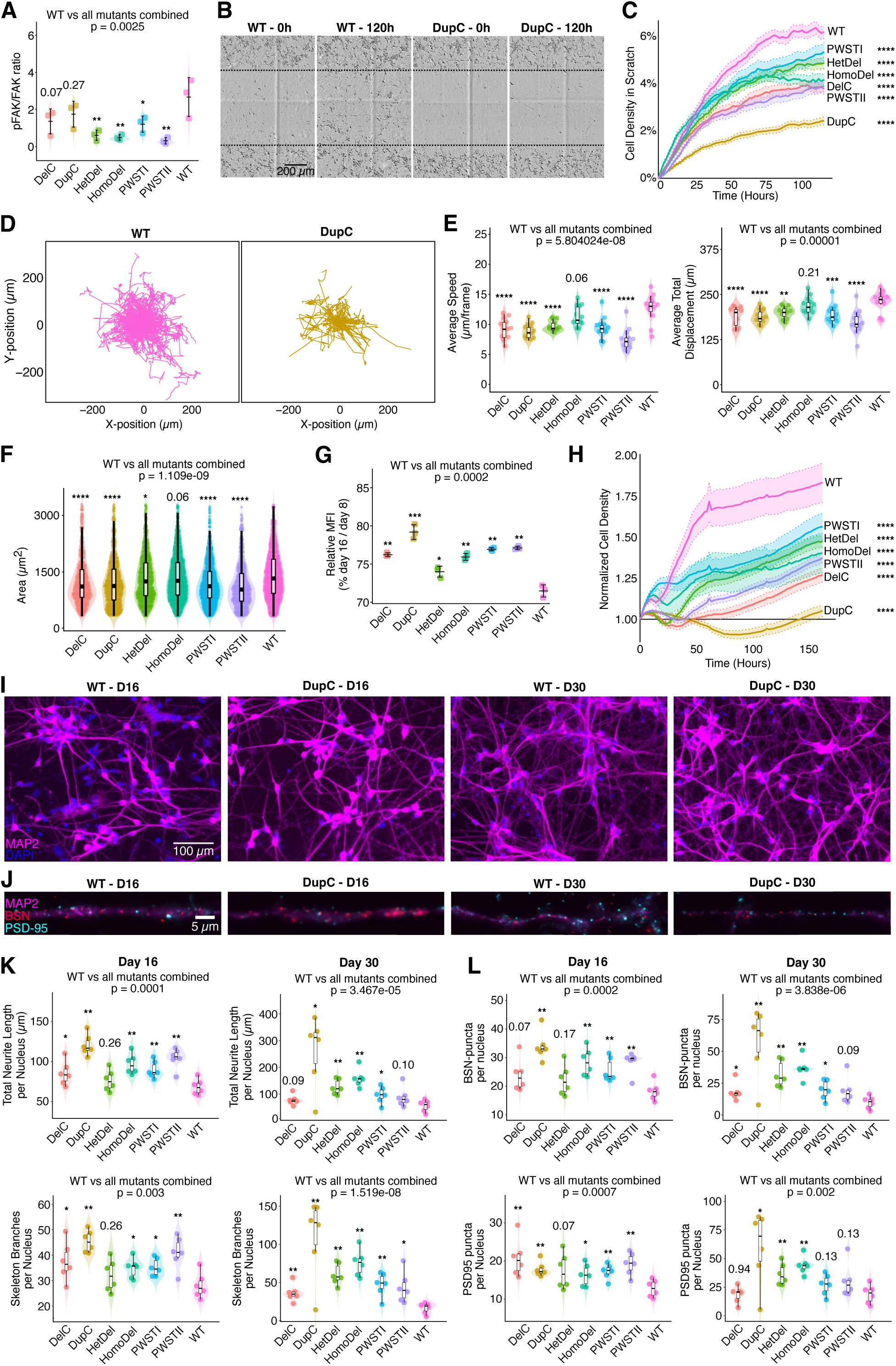
Characterization of dysregulated migration, proliferation and neurogenesis. (A) Western blot semiquantitative analysis of PFAK/FAK ratio (n=3) on d30 of differentiation with significance determined by ANOVA followed by Dunnett’s test. (B) Representative images of the migration assay from d12 until d17 of neuron differentiation. (C) Live cell imaging quantification of the percentage of scratch area covered by migrating cells (± SEM, n=12). (D-E) Representative spider plot of single-cell trajectories from migration analysis (D) with quantification of the average speed and total displacement of individual migrating neurons (E; n=12). (F) Quantification of spreading assay showing the average area per cell 120 minutes after seeding on d8 of differentiation (n=3). (G) Relative mean fluorescent intensity (MFI) from the flow cytometry Cytopainter dilution assay on d16 relative to d8 (n=3). (H) Live cell imaging quantification of normalized cell density from d9 until d16 of differentiation (± SEM, n=12). (I-J) Representative images of immunofluorescence staining for MAP2 and nuclei (I) and MAP2, BSN and PSD-95 (J) in WT and DupC neurons. (K) Quantification of total neurite length and skeleton branches per nucleus. (L) Quantification of BSN and PSD-95 puncta per nucleus. For (C) and (H), significance was determined by a Linear Mixed-Effects Model (LMM) followed by pairwise tests vs WT using Tukey’s correction. For (E), (F), (G), (K) and (L), significance was determined by an ANOVA, followed by pairwise Welch’s t-tests vs. WT. Error bars represent SEM (E, F, K, L) or SD (A, G), and asterisks denote statistical significance (*p < 0.05, **p < 0.01, ***p < 0.001, and ****p < 0.0001).

To further examine actin cytoskeleton organization, we quantified cell spreading areas^120^ by seeding cells at d8, fixing them 120 minutes later, and staining actin filaments (Figure S5H). As all MAGEL2-variant neurons showed significantly reduced spreading areas (Figure 6F), the dysregulated pathways identified by multi-omics analyses at d30 in SYS and PWS neurons appear to manifest already during intermediate stages of neuronal differentiation.

Given these early cytoskeletal alterations, we next asked whether they are accompanied by changes in proliferation, as seen across transcriptomics and proteomics analyses (Figure 1C, J, 5D). Using a fluorescent proliferation dye and flow cytometry, we observed higher fluorescence intensity, indicating reduced proliferation in all variants from d8 to d16, including DelC (Figure 6G). Live-cell imaging from d9 to d16 confirmed this finding, revealing reduced cell density, which was most pronounced in DupC, similar to the wound-healing results (Figure 6H, S5I). Of note, no proliferation defects were observed in undifferentiated hiPSC, in which *MAGEL2* is not expressed, further reinforcing that the phenotype is specific to NP and immature cortical neurons with both mutated and deleted *MAGEL2* (Figure S5J).

Consistent with our previous multi-omics findings, these data confirm that dysregulation of migration, adhesion, and proliferation manifests early during differentiation in SYS and PWS neurons.

### Both SYS and PWS neurons exhibit accelerated neuronal development and synapse development

We next set out to experimentally assess the upregulation of neuronal maturation pathways identified in our multi-omics data by analyzing dendritic architecture and synaptic marker distribution. Immunofluorescence (IF) staining of the dendritic marker MAP2, the presynaptic active zone protein Bassoon (BSN), and the postsynaptic scaffold protein PSD-95 (DLG4) at d16 and 30 of the differentiation was performed (Figure 6I). Maximum intensity projections identified BSN and PSD-95 puncta localized within dendrites and showed significantly increased MAP2-positive dendritic neurites (total length and branches per nucleus) in all variants across both timepoints (Figure 6J), indicating their enhanced morphological complexity. A similar trend was observed for BSN and PSD-95 puncta (Figure 6J), suggesting an increased number of synapses in SYS and PWS neurons. Quantification of synaptic puncta abundance and density, and BSN/PSD-95 colocalization (Figure S6) indicates an altered balance in pre-and post-synaptic organization in SYS and PWS neurons, consistent with the disrupted synaptic pathways revealed by the multi-omics analysis.

To test whether this accelerated structural maturation and synaptic development is accompanied by increased neuronal activation, we measured the ratio of phosphorylated CREB (pCREB) to total CREB following potassium chloride (KCl)-induced depolarization.^121,122^ Quantitative IF analysis revealed an elevated pCREB/CREB ratio at d16 across all variants, which was no longer evident at d30 (Figure 7A, B). These data indicate early premature activation or sensitization of the CREB signaling pathway during differentiation, which may contribute to the accelerated neuronal differentiation inferred from our integrated analyses.

**Figure 7:**
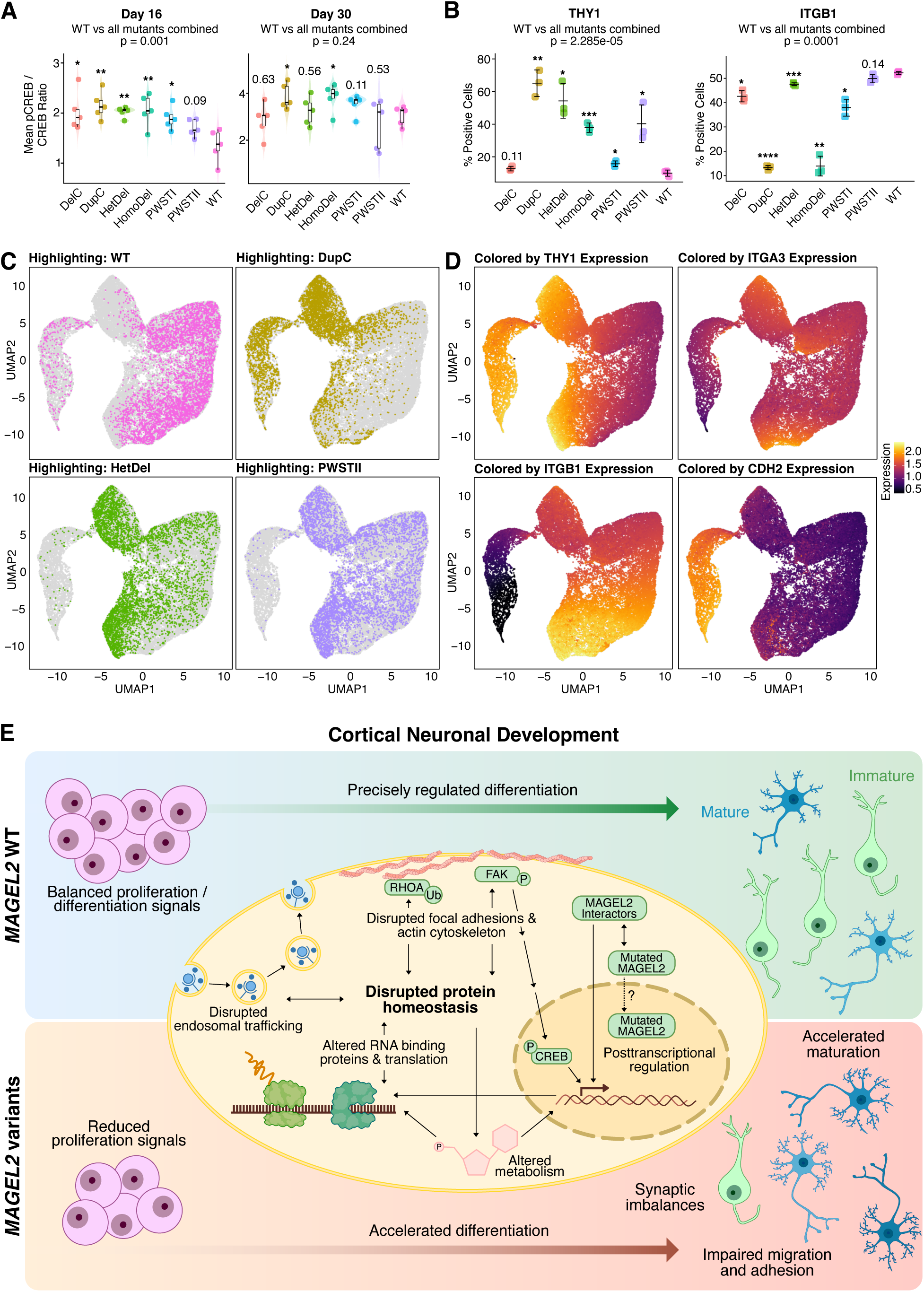
*MAGEL2* variants accelerate neuronal differentiation. (A) IF quantification of the pCREB/CREB ratio at d16 and d30 after KCl treatment (n=5). (B) Percentage of positive cells for the adhesion marker ITGB1 and the maturation marker THY1. (C, D) UMAP projection of single-cell data from differentiated d30 neurons, showing the distribution of cells based on the genotype (C, n=3) and expression of surface markers (D). (E) Proposed model illustrating how *MAGEL2* variants disrupt the balance of neuronal progenitor expansion and differentiation. For (A) and (B), significance was determined by an overall ANOVA followed by pairwise Welch’s t-tests vs. WT. Error bars represent SEM (A) or SD (B), and asterisks denote statistical significance (*p < 0.05, **p < 0.01, ***p < 0.001, and ****p < 0.0001).

We next examined whether the accelerated maturation was reflected in cell surface marker levels on a single-cell level by quantifying the expression of membrane proteins associated with neuronal maturation, adhesion, and migration, selected from dysregulated markers identified in the proteome using flow cytometry (Figure S7A). Hence, d30 neurons were stained for integrin beta-1 (ITGB1), integrin alpha-3 (ITGA3), Thy-1 cell surface antigen (THY1), and N-cadherin (CDH2). Uniform manifold approximation and projection (UMAP) of these single-cell data revealed a distinct marker-based cell distribution for each genotype (Figure 7C-D). Overall, WT neurons were enriched for ITGB1/ITGA3 associated with adhesion and migration (Figure 7C-D, S7B-C), consistent with roles of α3β1 integrin in neuron-glial interactions during cortical migration^123,124^ and showed lower THY1/CDH2 expression linked to postmitotic neurons^125^ and cadherin-linked adhesion remodeling.^126,127^ In contrast, the variants were enriched for THY1/CDH2 and displayed lower ITGA3/ITGB1 levels compared to WT, further indicating that SYS and PWS neurons undergo premature structural and functional maturation.

To facilitate further exploration of our unique and powerful datasets, we developed an interactive, open-access web portal: the MAGEL2 Multi-Modal-Omics Targeting Hubs (MAMMOTH; Figure S7D), providing the scientific community with complete and user-friendly access to the transcriptome, proteome, and ubiquitinome datasets obtained in this study. MAMMOTH serves as a comprehensive community resource, enabling researchers to independently perform additional analyses on our data and test novel hypotheses to accelerate progress in understanding the molecular basis of MAGEL2-related NDDs.

## DISCUSSION

Building upon the established *PWSTI* and *PWSTII* hiPSC lines^28^, we engineered SYS *c.1996dupC* and *c.1996delC* mutations, as well as heterozygous and homozygous *MAGEL2* deletions, within the same two backgrounds. This strategy enabled robust multi-omics cross-validation while minimizing donor-dependent variability of cortical neurons derived from a total of 14 hiPSC lines (six variants, including an isogenic WT control per background). Integrative analyses combining transcriptomics, proteomics, and ubiquitinomics, together with functional assays, identified *MAGEL2* as a central regulator of human cortical neuron differentiation, primarily driven by major changes in the proteome. Disruption of *MAGEL2* in both SYS and PWS models shared phenotypes, including reduced proliferation, accelerated neuronal maturation, and impaired adhesion and migration, partially converging in hallmarks of other NDDs.

Among disease-specific findings, GO terms related to ribosome biology and splicing emerged in PWS-specific downregulated DEGs. PWS-specific DEPs were enriched for mitochondrial, proteostatic, neuronal, and synaptic perturbations, while DEUs were related to ribosomal biology. These findings strongly align with previous studies of the PWS locus demonstrating the role of *SNORD*/*SNRPN* in RNA processing and translation^42,43,60–62^, *NDN* in neuronal differentiation^128,129^, and *MKRN3* in ubiquitin-mediated homeostasis^130^, thereby confirming the robustness of our multi-omics data.

Strikingly, DupC neurons showed by far the highest number of specific DEGs, with GO terms enriched for developmental and proliferative signaling, including pathways associated with nuclear function. Subsequent functional studies uncovered the most severe defects in proliferation, migration and neurite outgrowth in DupC neurons compared to the other *MAGEL2* variants, which is consistent with the more severe phenotype in SYS patients.^11,117,131^ As we did not observe any “leaky” *MAGEL2* expression from the silenced maternal allele in the neurons with heterozygous *MAGEL2* deletion lines, our data support the hypothesis that the DupC-specific phenotype may result from the predicted truncated MAGEL2 protein, potentially mis-localizing to the nucleus and associated with aberrant RNA metabolism^30,31,132,133^. This idea is further supported by the observation that the *MAGEL2* transcript was consistently upregulated in DupC neurons, with a similar trend in DelC, suggesting a distinct posttranscriptional regulatory context in SYS-related *MAGEL2* mutations, the mechanisms of which remain to be elucidated.

In addition to these disease-specific signatures, we identified shared features between SYS and PWS models, characterized by reduced proliferative capacity, accelerated neuronal maturation, and impaired adhesion and migration, underscoring the significant contribution of MAGEL2 to human cortical neuron differentiation. These data are consistent with the high, transient *MAGEL2* expression in outer radial glia cells during the first trimester of pregnancy.^134^ Similar effects on proliferation and differentiation have been reported in cells with *SNORD116* deletion, which notably also show decreased *MAGEL2* expression despite intact *MAGEL2* sequence and imprinting^42,43,135,136^, supporting a previously reported mutual regulatory relationship between these two loci.^42^ However, the RNA-seq depth of 25 million reads per sample used in this study generally did not allow for reliable quantification of *SNORD* gene expression levels, which are typically low-abundance and non-polyadenylated.^137^ Consequently, we could not assess whether *MAGEL2* deletions also alter expression at the *SNORD*/*SNRPN* locus. As fetal neural stem cells from a paternal 15q11-13 duplication mouse model including duplication of paternal *MAGEL2* show opposite effects, such as increased proliferative capacity and reduced neuronal differentiation^138^, a dosage-dependent role of *MAGEL2* is likely required in balancing the proliferation-differentiation ratio. Moreover, dysregulated genes shared between SYS and PWS were significantly enriched in genes associated with NDD, ASD, ID, and epilepsy, consistent with the high prevalence of these phenotypes in both syndromes.^8,117,131,139–142^

In agreement with previous studies^23–29,34^, our proteome data confirm the known role of MAGEL2 in protein homeostasis, which was commonly disrupted in SYS and PWS neurons and strongly linked to their phenotypes such as neurodevelopmental delay, hypotonia and abnormal cerebral morphology. The MAGEL2 interactome data indicate that these phenotypes are linked to dysregulated vesicle trafficking, RNA metabolism, and metabolic networks. Dysregulated proteins were significantly enriched for miRNA and RBP motifs previously implicated in ASD and NDD.^95–103,107,108^ Thus, our results highlight MAGEL2 as a regulator of post-transcriptional regulation and further support its role in RNA metabolism.^31^

Despite the technical variability of pull-down based ubiquitinomics^143^, our ubiquitinome profiling of SYS and PWS neurons revealed shared dysregulation of metabolism and cytoskeletal pathways, consistent with our transcriptome and proteome findings. Integrating proteome and ubiquitinome data, we identified dysregulated ubiquitination of LIN28A, involved in RNA regulation of neural progenitor cells^78,79^, TAGLN3 and RHOA both linked to cytoskeleton biology^81–85^, indicating selective post-translational control of cytoskeletal and signaling proteins by MAGEL2 in SYS and PWS neurons, providing novel candidates for future studies. Our findings, however, argue against a primary role for MAGEL2 in ubiquitin-mediated protein homeostasis, highlighting its role in post-transcriptional processes and RNA metabolism.^31^

Most strikingly, the dysregulated proteome dominated the shared signature across all *MAGEL2* variants, with 1,700 shared dysregulated proteins and 453 enriched GO terms. MOFA2 independently confirmed this dominance with latent factor 1 accounting for ∼30% of proteome variance (compared with ∼5% for the transcriptome and ∼9% for the ubiquitinome) across all M*AGEL2*-variant neurons. In line with the MOFA framework, which interprets leading factors as the main shared sources of biological variation across omics layers^109,110^, we focused on Factor 1 as other factors mainly captured background variation without clear variant relevance. Overall, factor 1 displayed top loadings, including DCX, related to neuronal migration, ribosomal proteins RPL13/39/7, and RHOA. Consistently, pathway enrichment revealed changes in neurogenesis and axon guidance deficits toward mutants, mitochondrial shifts toward WT, and bidirectional alteration in RNA and translation. HPO analysis directly matched SYS and PWS clinical phenotypes specifically in proteome and ubiquitinome, further confirming our previous layer-specific and integrative multi-omics analyses.

Functional assays validated reduced proliferation, accelerated neuronal maturation, and altered synaptic morphology together with impaired adhesion and migration, underscoring a strong contribution of *MAGEL2* to cortical development defects shared between SYS and PWS neurons. Moreover, we revealed that defects in adhesion and migration assessed by migration speed and cell spreading area arise early during differentiation (d8 - d19). In addition, we also found increased BSN/PSD-95 puncta and neurite branches, indicating an altered pre- and post-synaptic organization linking the ASD-associated synaptic scaffold protein PSD-95, encoded by the *DLG4* gene^51,52^ to MAGEL2. In contrast to our study, reduced dendrite length was observed in patient-derived SYS iNeurons driven by NGN2 overexpression^37^ and a lower number of PSD-95-positive synapses was reported in cortical neurons differentiated using a minimally guided protocol derived from hiPSC carrying a CRISPR/Cas9 engineered *MAGEL2* frameshift mutation that introduces a premature stop codon^35^. iNeurons^37^ undergo an accelerated differentiation that largely bypasses a progenitor-like phase, while the minimally patterned cortical protocol includes only FGF2 and EGF as patterning molecules. Our data, however, were obtained from neurons generated by a protocol that relies on dual SMAD inhibition and small-molecule induced patterning yielding layer VI-enriched cortical neurons.^41^ Hence, we speculate that this discrepancy may result from differences in neuronal subtype composition, each potentially influencing cytoskeletal and synaptic architecture in distinct ways.

Our data are in line with PWS/SYS clinical manifestations such as decreased brain volume and altered cortical morphology observed in PWS patients^135,144–150^, linking the reduced progenitor pool in SYS and PWS neurons to aberrant brain morphology. As reported in Tuberous Sclerosis Complex and other NDDs^151–154^, disrupted neuronal migration showed in SYS and PWS may underlie cortical malformation and cognitive impairment. Moreover, accelerated neuronal maturation in SYS and PWS neurons is consistent with emerging evidence that aberrant expression of ASD-associated genes leads to common asynchronous neuronal development through different but converging mechanisms^155–161^, further confirming the relevance of our data to broader neurodevelopmental mechanisms associated with ASD. A recent study further identified a disbalance between proliferative and neurogenic processes, together with disruption of microtubule biology and RBPs, as convergent features of ASD-related neurodevelopmental disruption.^157^ These findings underscore the biological relevance of our data, suggesting that SYS and PWS share a common MAGEL2-dependent mechanism that converges on pathways governed by other ASD-associated genes, thereby affecting the timing of neuronal maturation, migration and circuit assembly. Thus, based on our data, we propose a model in which MAGEL2 acts as a coordinator of protein homeostasis of key factors regulating progenitor proliferation, cortical neuronal differentiation, and cytoskeletal processes such as migration and focal adhesion. Disruption of these processes in SYS and PWS neurons results in a MAGEL2-dependent imbalance in the regulation of an accelerated transition from proliferative progenitors to differentiating neurons.

Collectively, our multi-omics analysis of cortical neurons derived from 14 CRISPR/Cas9-engineered hiPSC lines modeling SYS and PWS, complemented by functional assays, revealed both syndrome-specific and common molecular programs, thereby establishing MAGEL2 as a central driver of shared pathophysiology. Consistent with growing evidence that genetic NDDs converge on shared biological pathways, our data show that MAGEL2 variants affect core processes implicated in NDDs. Together with the multi-omics data accessible through the open-access MAMMOTH platform, these findings provide a valuable basis for future research and translational exploration, facilitating the identification of snoRNA-specific targets and their regulatory crosstalk with MAGEL2, the elucidation of variant-specific effects, and the evaluation of candidate therapeutic strategies such as antisense oligonucleotides (ASOs) or MAGEL2-activation from the silenced maternal allele.

### Limitations of our study

We generated multi-omics data from six isogenic variants and their isogenic controls in two different genetic backgrounds, enabling background-overlapping analyses per genotype. Thus, these analyses strengthen our confidence in a shared role of MAGEL2 in SYS and PWS cells in driving an accelerated transition from proliferative progenitors to differentiating neurons. Functional tests confirmed these defects and highlighted aberrant adhesion, migration and synaptic morphology in SYS and PWS neurons. However, to better exclude inter-individual variability as a confounding factor^162–164^, the inclusion of CRISPR/Cas9 engineered *MAGEL2* variants in additional genetic backgrounds in future studies would be beneficial.

Our current work focused on shared molecular features across SYS and PWS. Given that the SYS-associated *MAGEL2* mutations studied here, DupC and DelC, showed limited overlap in their signatures, the availability of our data now enables more targeted, mutation-specific investigations into their distinct molecular mechanisms and potential differential contributions to disease pathogenesis, with implications for developing mutation-tailored therapeutic strategies.

Moreover, studies of additional SYS-associated *MAGEL2* mutations in our cellular system would help to better define genotype-phenotype relationships across the broader SYS spectrum. Since our work was restricted to cortical neurons, analyses of other disease-relevant brain regions, particularly the hypothalamus, will reveal whether the phenotypes observed in cortical neurons generalize across neurodevelopmental contexts relevant to SYS and PWS.

Our work is based on 2D neurons that represent a powerful system for studying human cortical development.^41^ However, it does not fully recapitulate the cellular diversity and 3D architecture of the developing brain. Hence, studying 3D neural organoids combined with single-cell and spatial resolution will provide additional insight into the migration defects and accelerated maturation in SYS and PWS neurons. While the multi-omics data, including transcriptome, proteome, and ubiquitinome, strongly indicate dysregulated protein homeostasis as a major driver of the SYS and PWS pathophysiology, we only exemplarily tested phosphorylation of FAK and CREB as an additional layer of signaling-relevant post-transcriptional regulation. Comprehensive phospho-proteomics, as well as metabolomics will provide additional insights into the role of *MAGEL2* in cortical development. Following our identification of MAGEL2 interactors and top multi-omics hits as candidate targets, CRISPRi/a perturbations of these factors in future studies will help functionally validate the implicated pathways in SYS and PWS, confirm interactome-based predictions, and prioritize potential therapeutic targets.

## RESOURCE AVAILABILITY

### Lead contact

Further information and requests for resources and reagents should be directed to and will be fulfilled by the Lead Contact, Magdalena Laugsch

## Acknowledgements

This work was supported by Foundation of Prader-Willi Research grants to M.L., C.P.S., and M.E.T., an R01 grant HD096326 by the U.S. National Institutes of Health to M.E.T., and a Marsilius Fellowship by Heidelberg University to C.P.S. We thank Core Facilities at Heidelberg University: the FFCF for providing equipment and support for the flow cytometry experiments, the ZMBH Imaging Facility and Holger Lorenz for equipment and support with the imaging experiments as well as the Deep Sequencing Core Facility and David Ibberson for performing the RNA sequencing. In addition, we acknowledge the Baden-Württemberg High Performance Computing cluster for supporting the computational analyses. We thank Daniela Mauceri for her advice on the CREB signaling analyses and Michaela Bartusel for her for her discussion and feedback on the manuscript and Aylin Camgöz for her support with the initiation of the flow cytometry studies.

## Contributions

M.L., C.P.S., and J.B. conceptualized the study. Experimental data were generated by J.B. with contributions from B.E.G., A.S.A., T.W., T.B., F.H.S., K.H., M.S., and S.T.; bioinformatic analyses were carried out by M.E., J.B., B.E.G., and A.S.A.; hiPSC lines harboring the MAGEL2 point mutation and PWS deletions were provided by C.E.d.E., D.J.C.T., and M.E.T.; J.B. assembled all figures; M.L., J.K, and C.P.S. contributed to the interpretation of results; M.L. and J.B. wrote the manuscript; all authors reviewed and approved the final version of the manuscript.

## DECLARATION OF INTERESTS

The authors declare no conflicts of interest.

## STAR METHODS

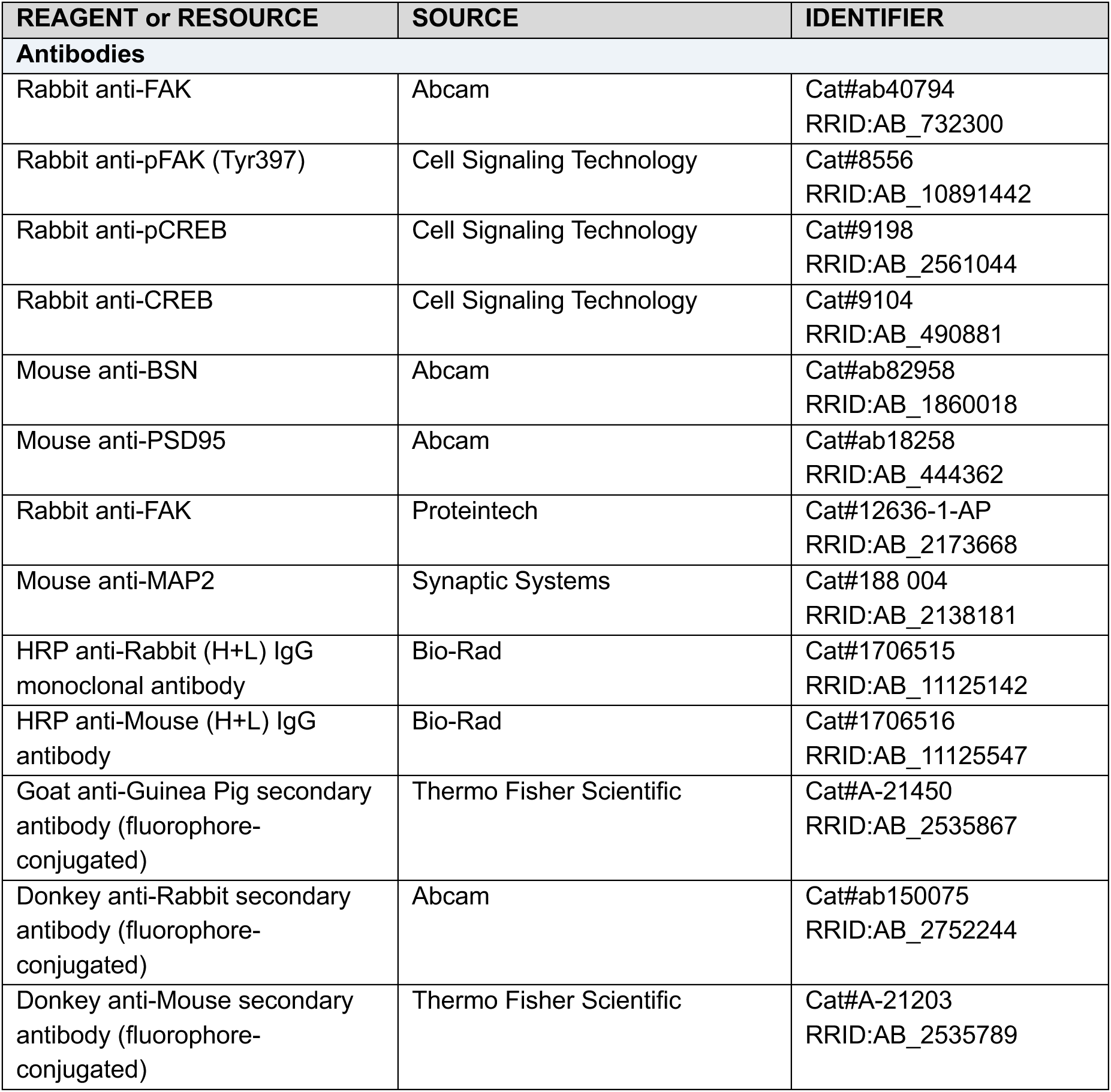
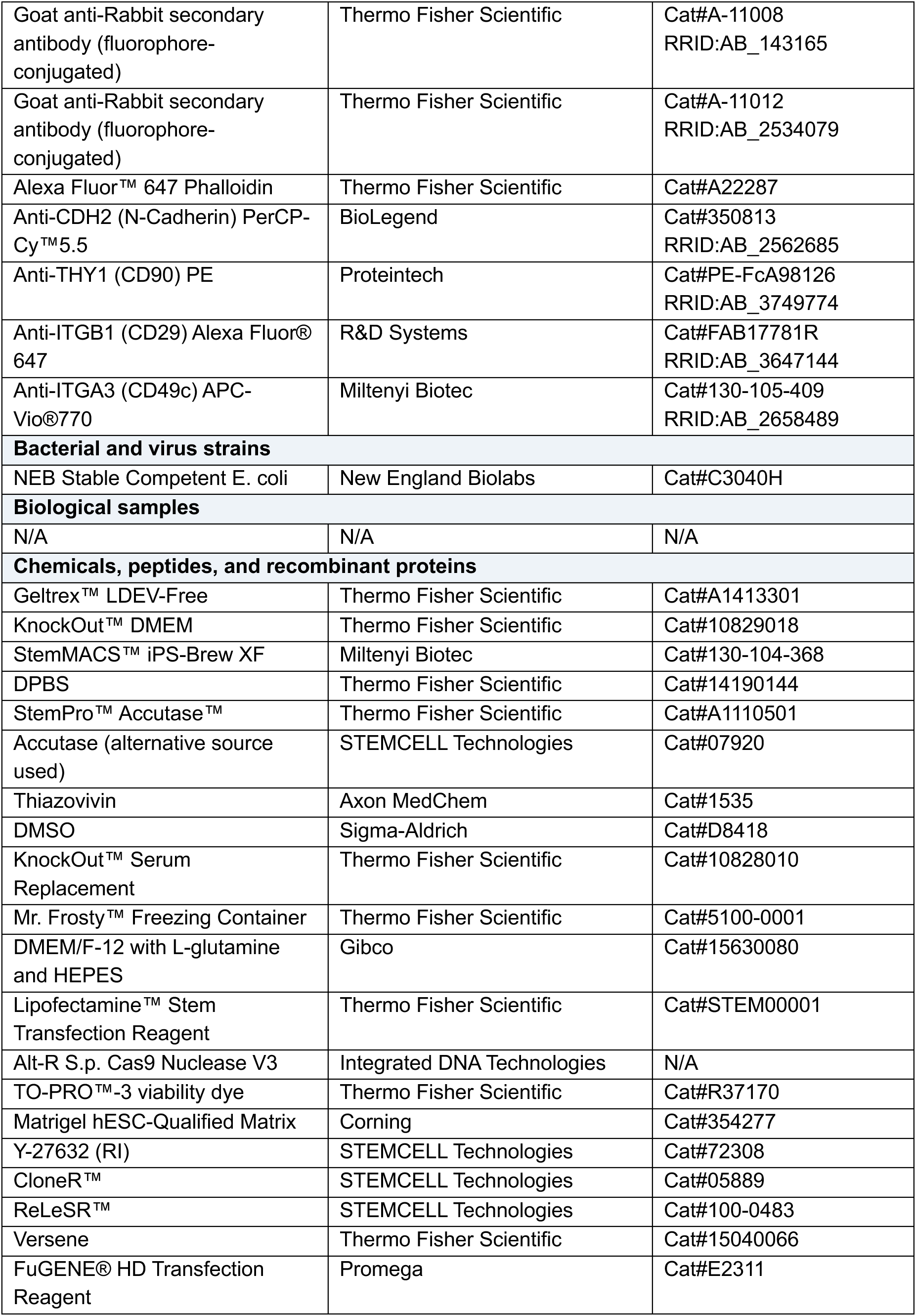
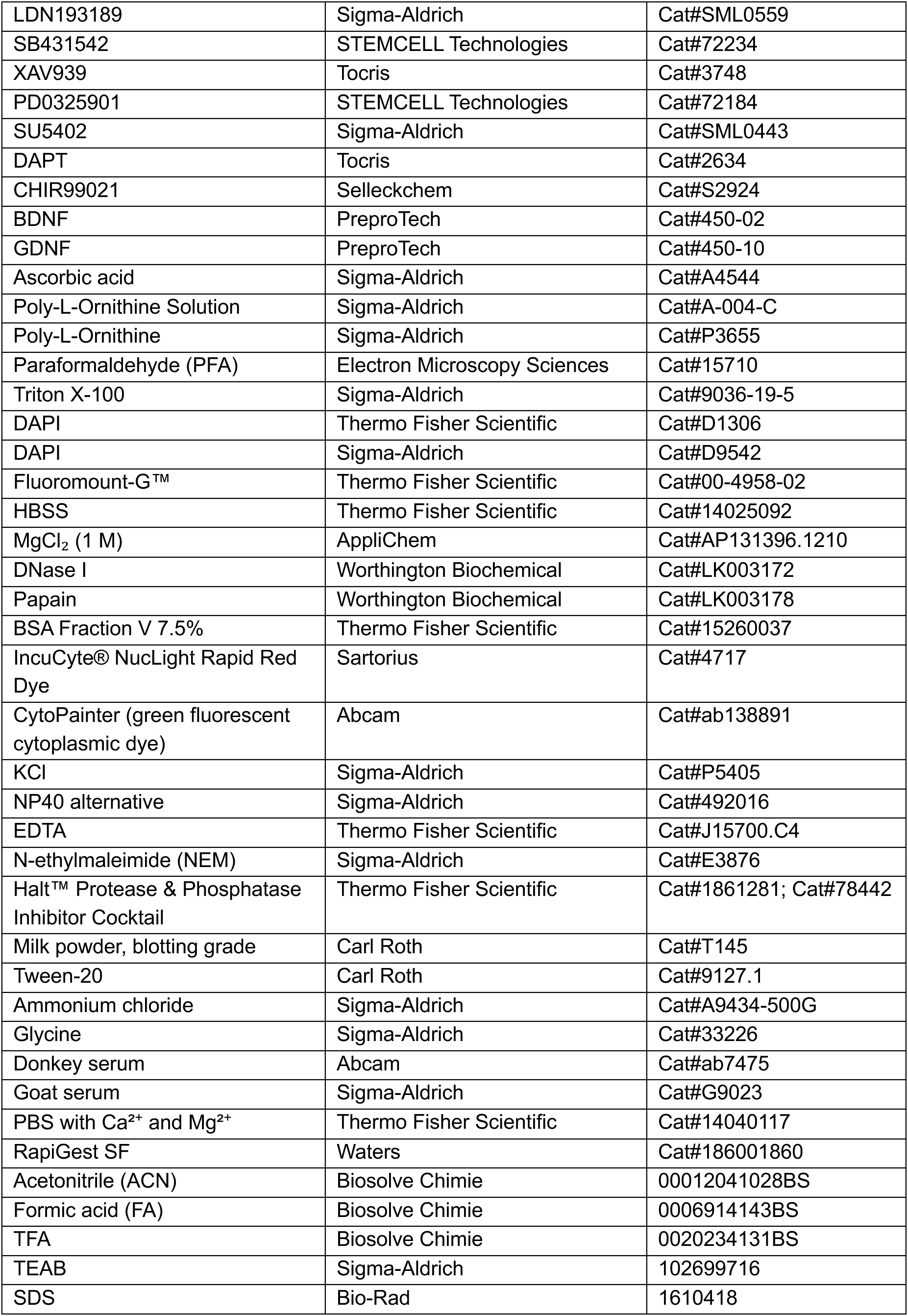
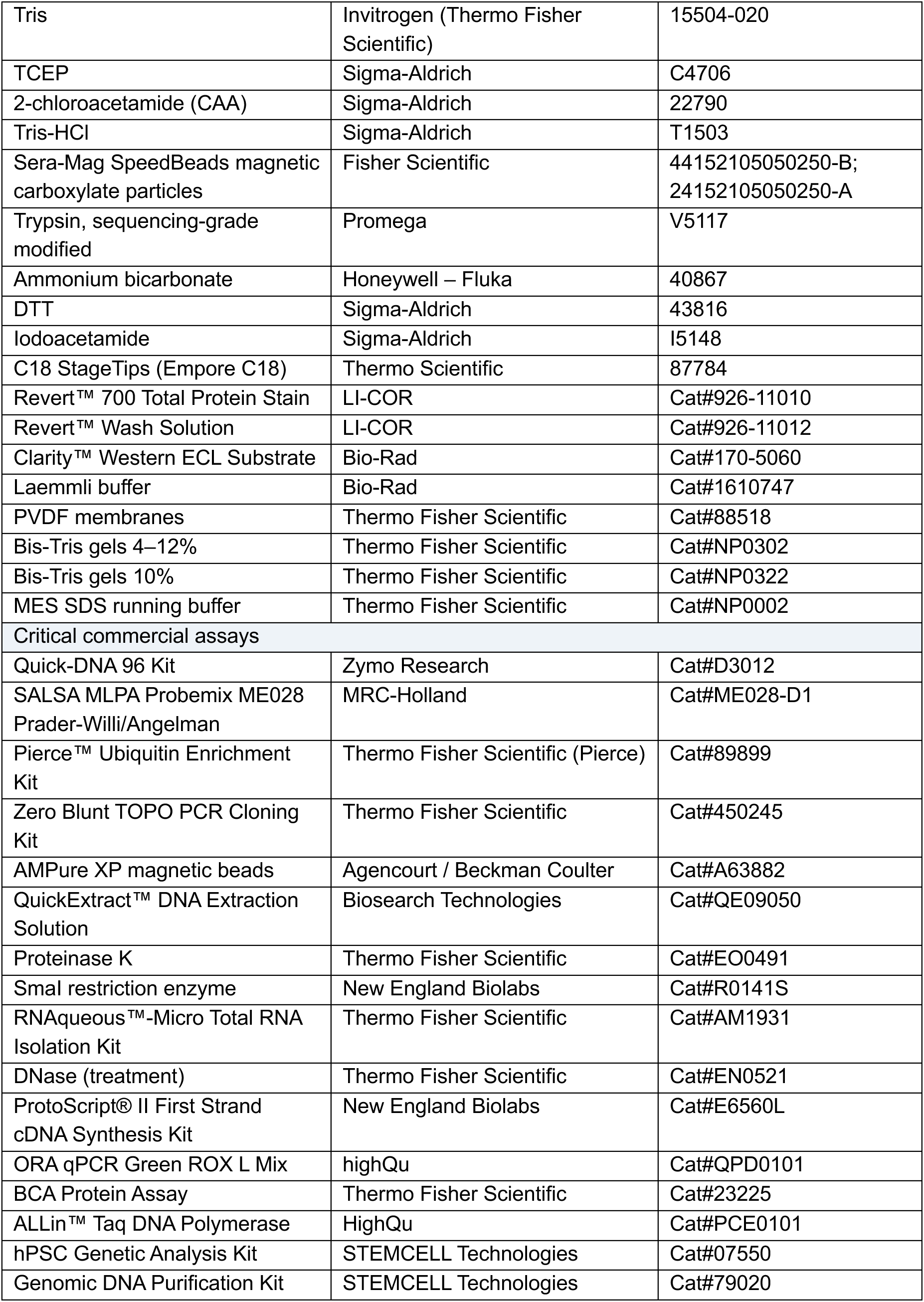
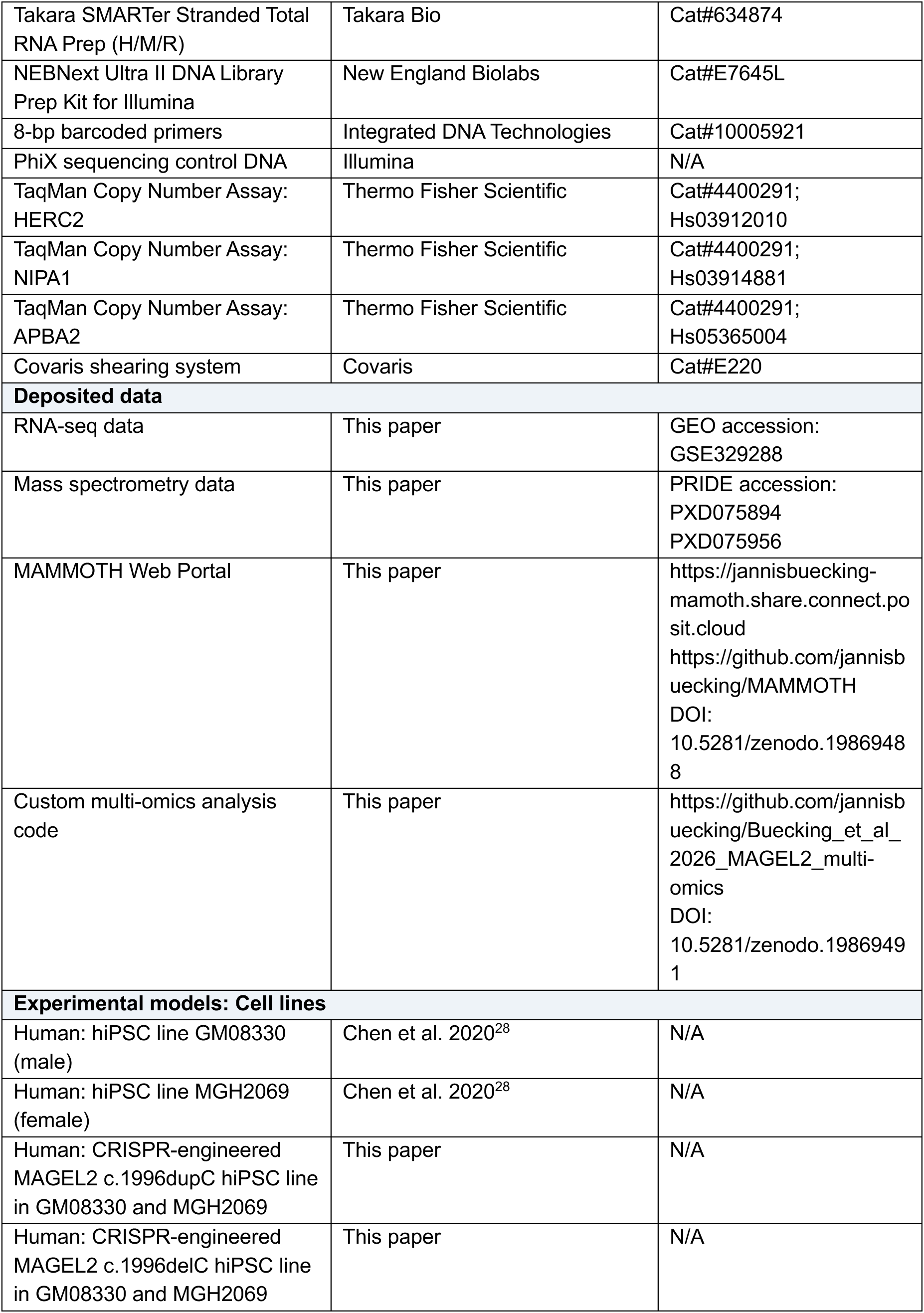
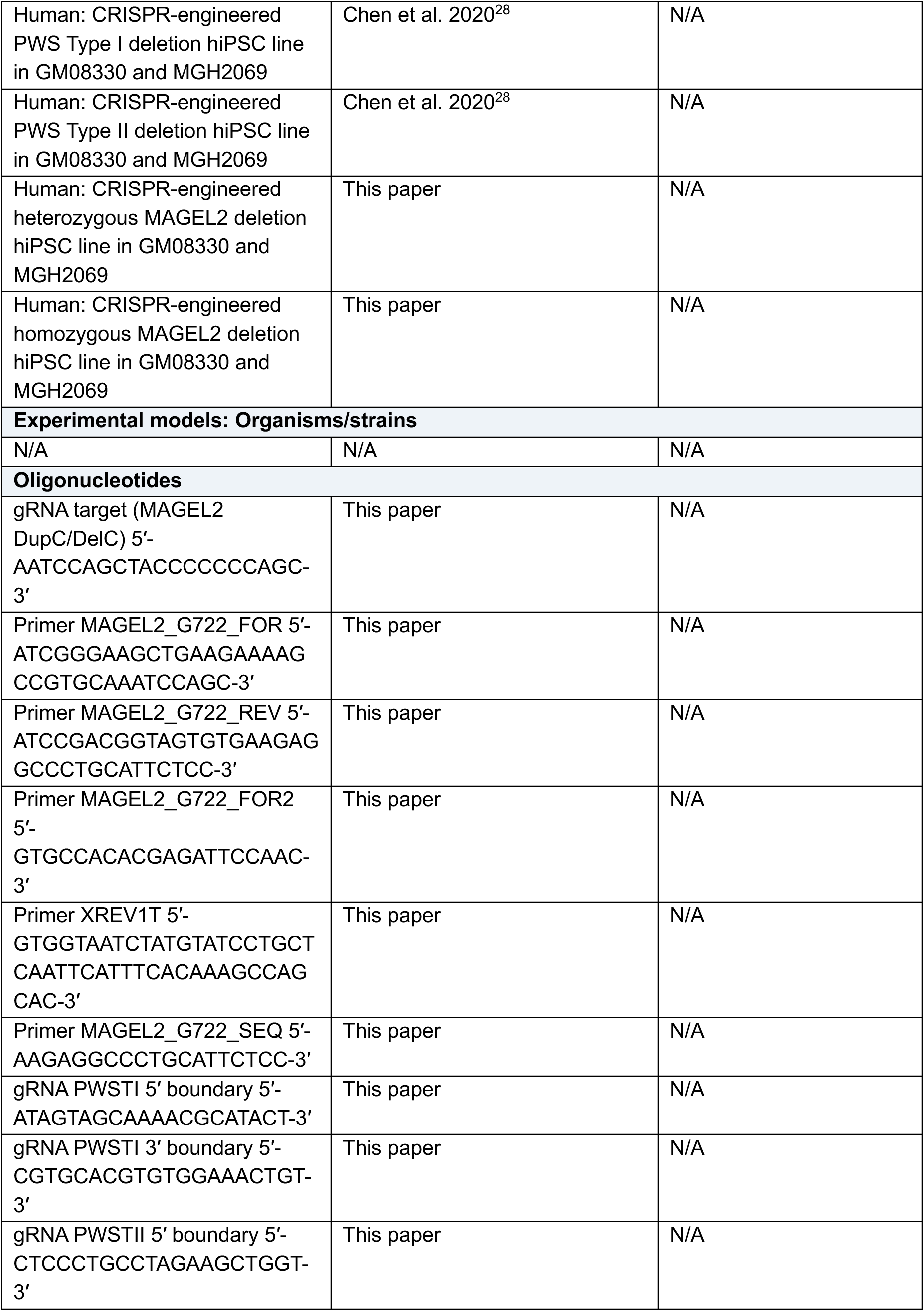
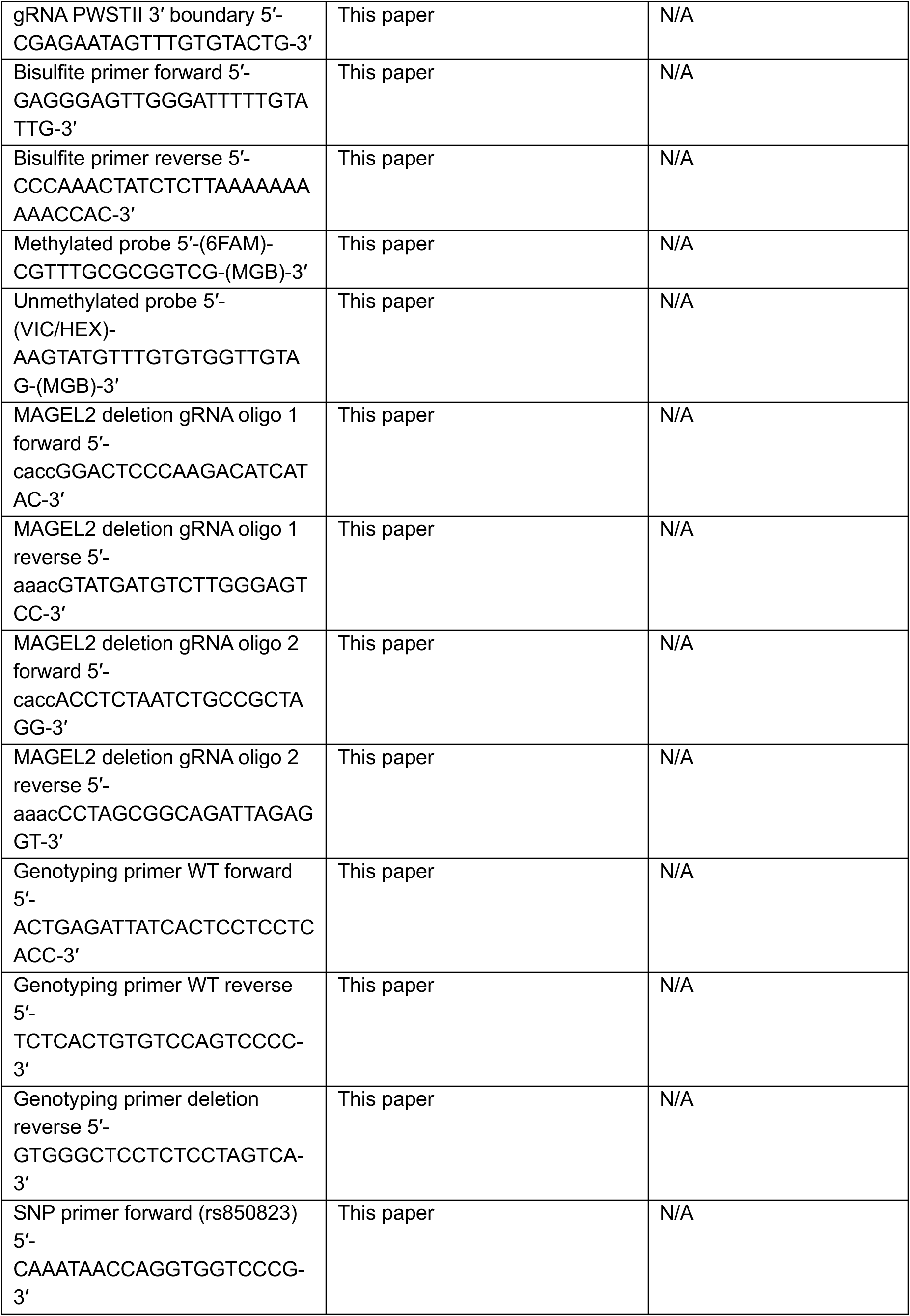
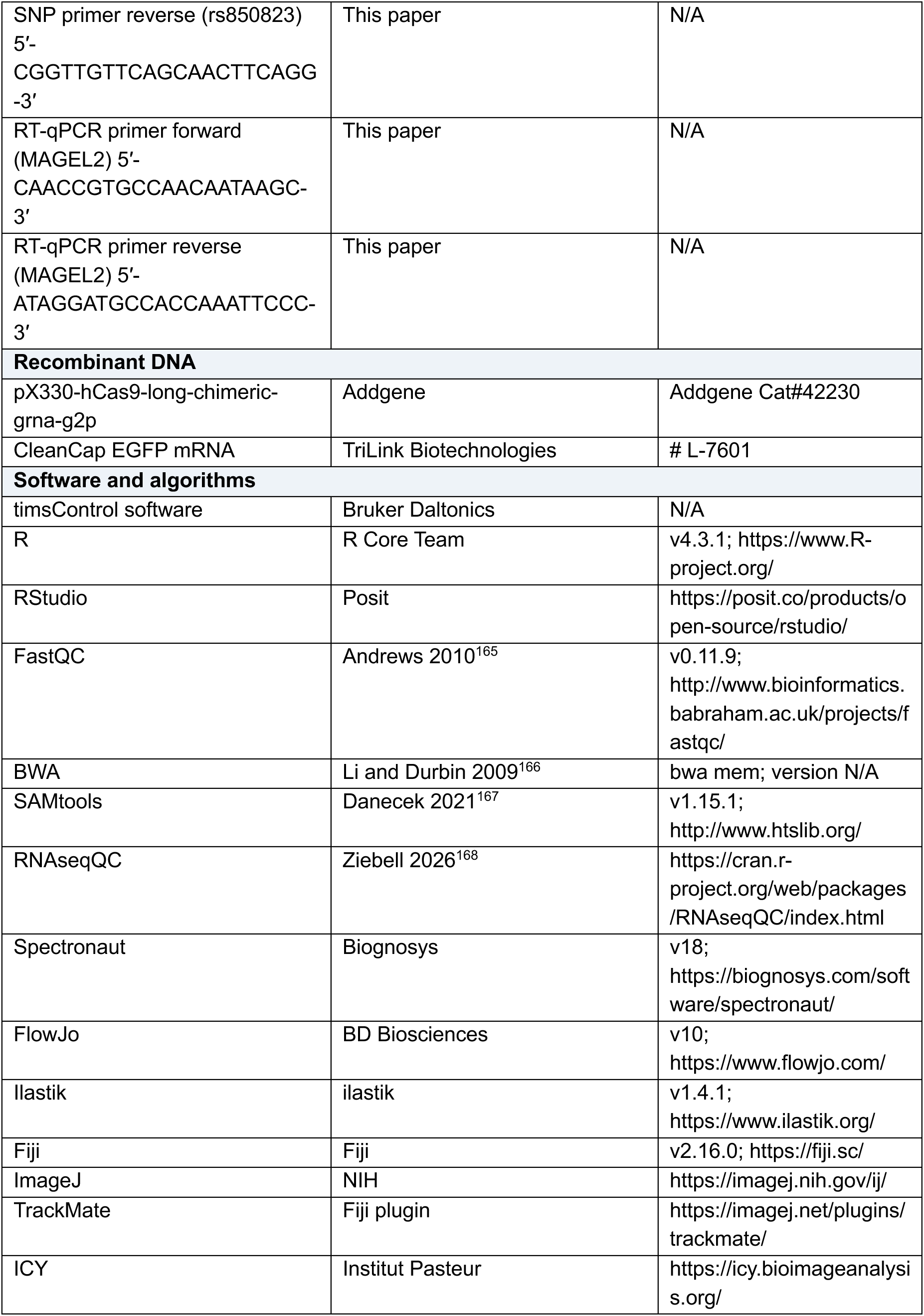
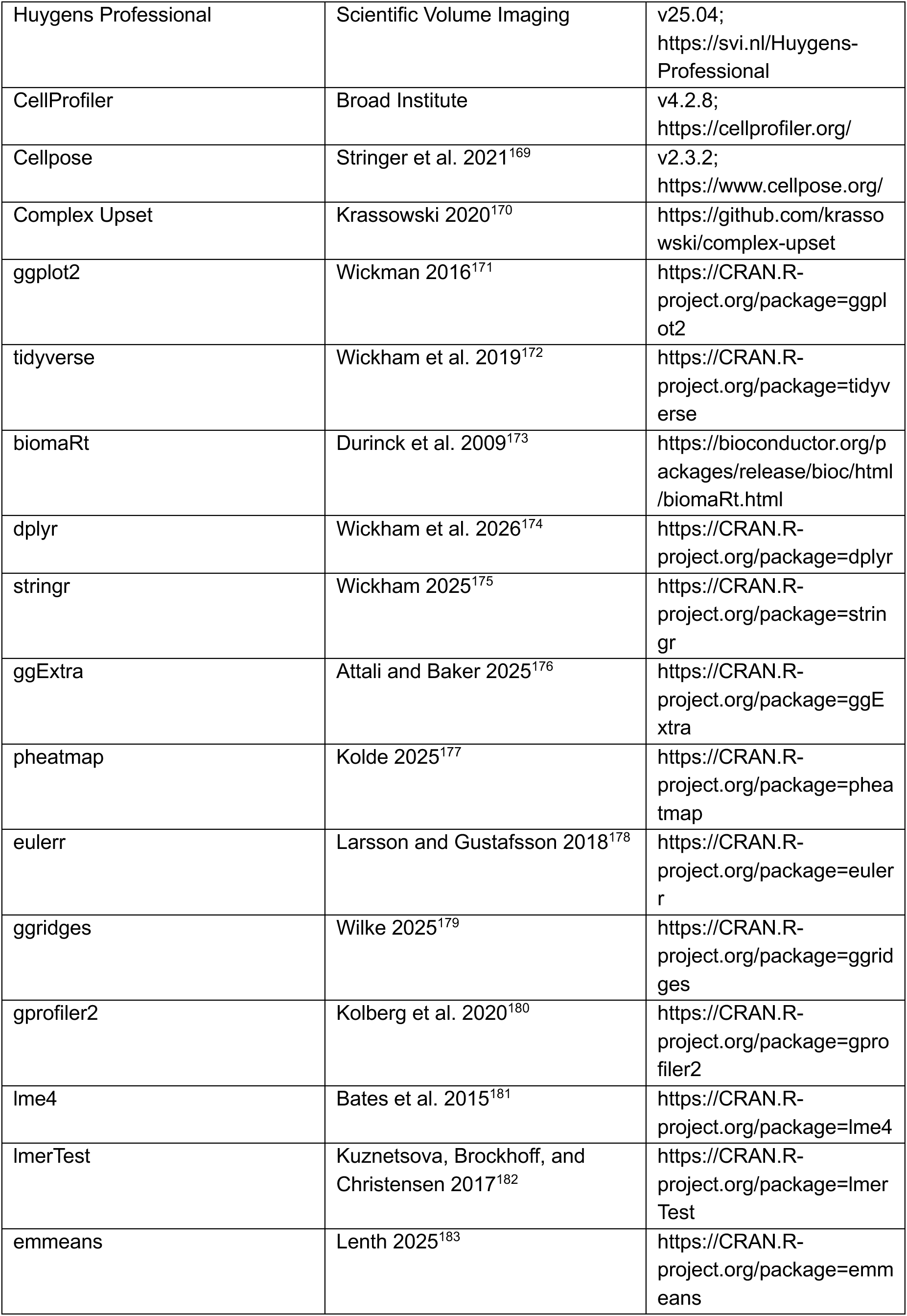
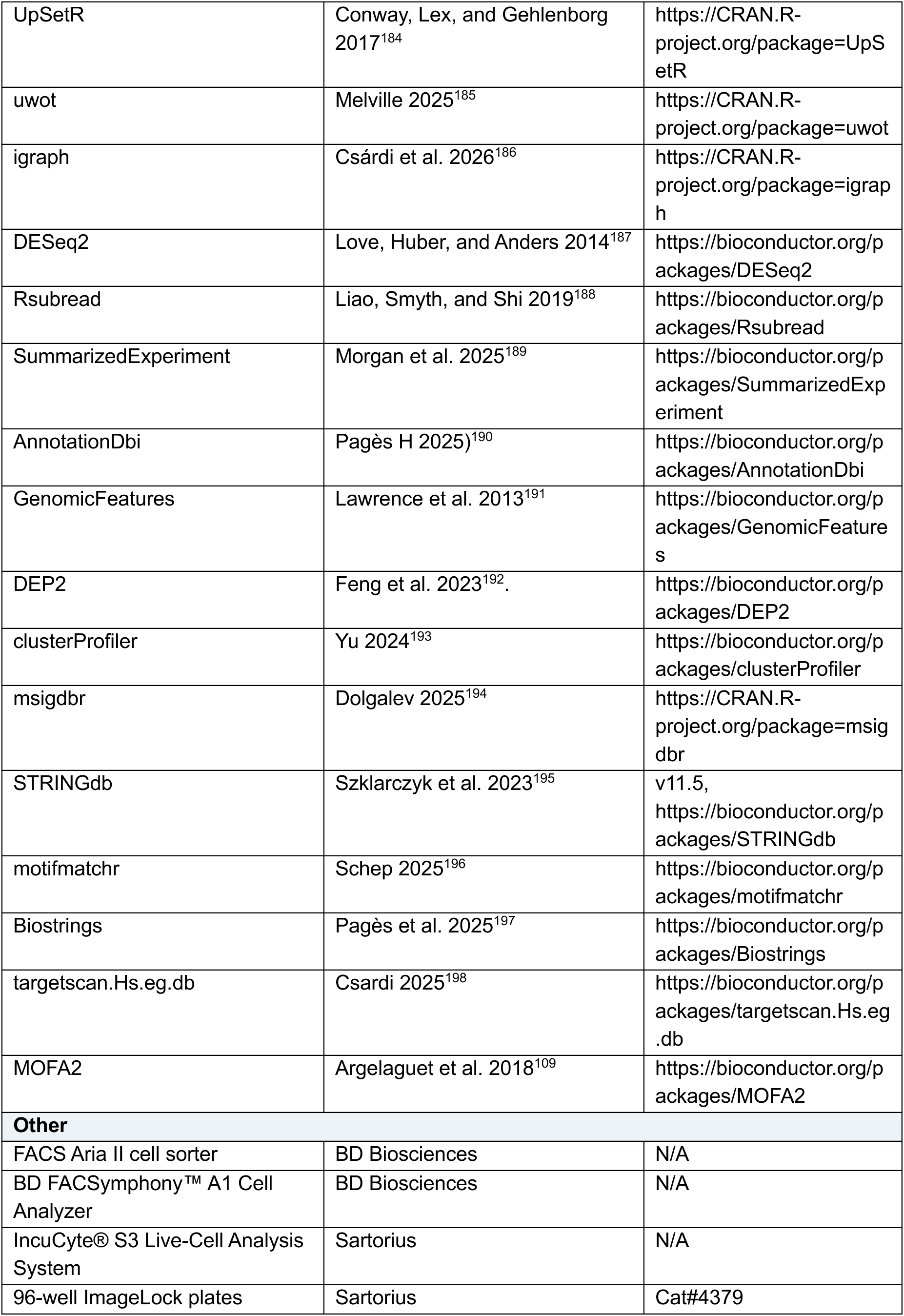
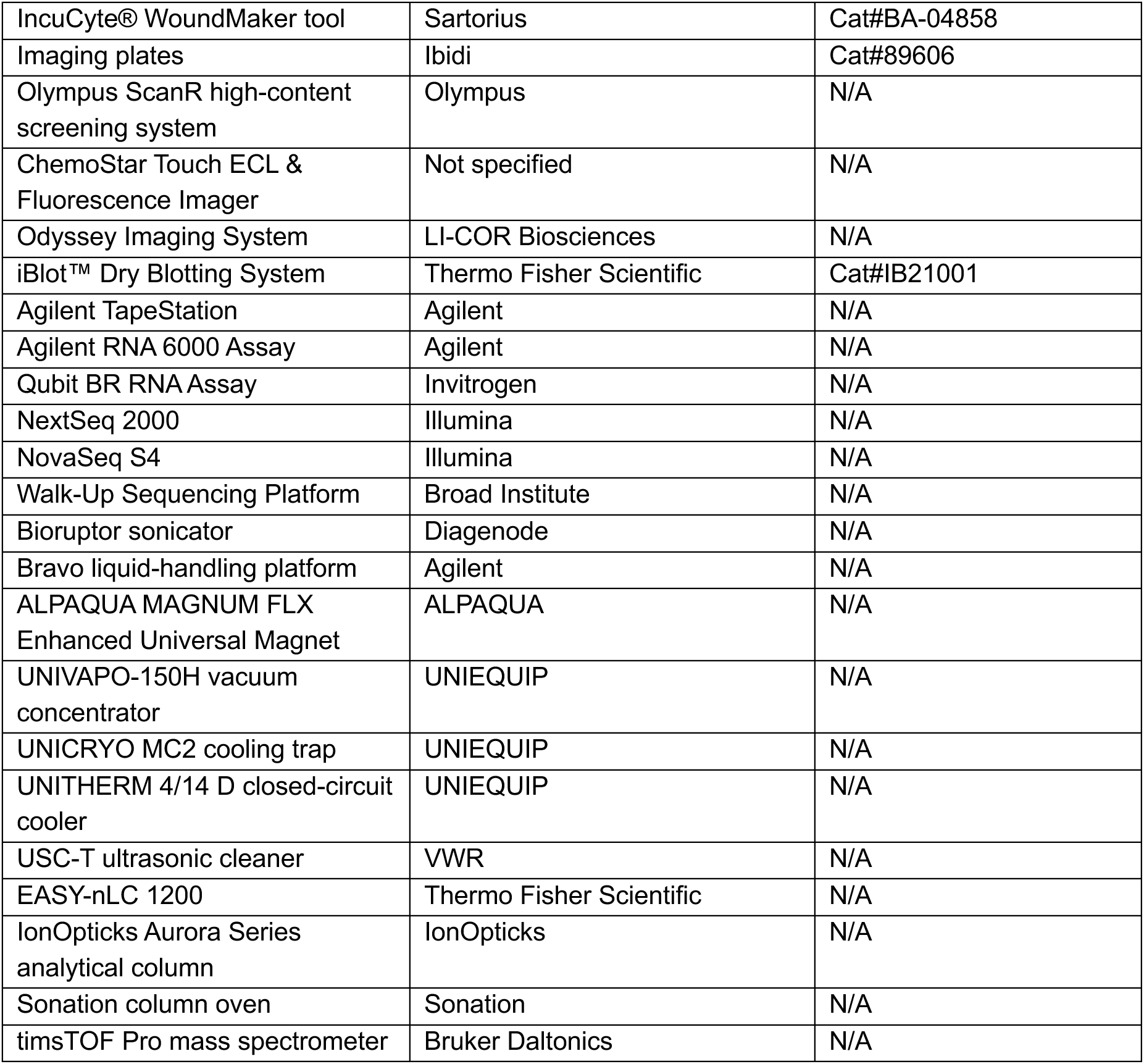

### hiPSC culture

hiPSC (background male GM08330 and female MGH2069) were grown in dishes coated for at least 1 hour at 37°C with Geltrex™ LDEV-Free (ThermoFisher #A1413301) diluted 1:100 in KnockOut™ DMEM (ThermoFisher #10829018) and cultured in StemMACS™ iPS-Brew XF (Miltenyi Biotec #130-104-368) media. Once the cells were ∼80% confluent, the medium was aspirated and cells were washed once with DPBS (ThermoFisher #14190144), following incubation with StemPro™ Accutase™ (ThermoFisher #A1110501) for 7 min at 37°C. Cells were resuspended in StemMACS™ iPS-Brew XF supplemented with 2 µM Thiazovivin (Axon Med Chem #1535), counted with 0.4% trypan blue and seeded between 2 × 10^4^ cells/cm^2^ to 5 × 10^4^ cells/cm^2^. Thiazovivin was removed after 24 hours. To freeze, cells were detached with Accutase and resuspended in freezing medium (10% DMSO (Sigma-Aldrich #D8418), 30% iPS-Brew XF and 60% KnockOut™ Serum Replacement (ThermoFisher #10828010)), before transferring to a cryovial for long-term storage. Filled cryovials were transferred to Mr. Frosty™ Freezing Container (ThermoFisher #5100-0001) and placed inside a -80°C freezer for gradual cooling. After 24 hours at -80°C, cryovials were transferred to -150°C for long-term storage. To thaw, hiPSC were placed at 37°C in a water bath until a small ice crystal remains, mixed with 13 mL DPBS and centrifuged for 5 min at 300g on room temperature (RT). The supernatant was aspirated, and cells were resuspended in iPS-Brew XF supplemented with 2µM Thiazovivin. Cells were counted with 0.4% trypan blue and seeded between 2 × 10^4^ cells/cm^2^ to 5 × 10^4^ cells/cm^2^. Thiazovivin was removed after 24 hours.

### Generation of CRISPR/Cas9-engineered DupC and DelC mutation hiPSC lines

*MAGEL2* coding sequence was parsed to identify all candidate *S. pyogenes* Cas9 gRNA sequences, 20 bp sequences preceding a canonical 3’ NGG protospacer adjacent motif (PAM). All candidates were scored for predicted efficiency using Azimuth 2.0^199^ and for predicted off-target editing potential using Cas-OFFinder^200^ with the human reference genome GRCh38/hg38, allowing up to five mismatches (not counting mismatches at the degenerate first PAM position). Cas-OFFinder output was analyzed to count off-target sequences with 0-5 mismatches to each candidate, and for each off-target sequence, the Cutting Frequency Determination score was calculated. Candidate guides targeting sequences near the locus of interest were further assessed using inDelphi^201^, which provides predictions of editing outcomes for each guide design. One guide (DNA target: 5’-AATCCAGCTACCCCCCCAGC-3’) was manually selected for editing experiments after comparing candidates’ locations relative to the *MAGEL2* indels to be generated, predicted on-target activities, predicted targeting specificities, and predicted editing outcomes. One day prior to transfection, cells were seeded into a new Matrigel-coated well in a 12-well tissue culture plate by following the procedure above for passaging, except resuspending cells in working medium supplemented with 10 µM ROCK inhibitor (RI, Y-27632) and pipetting resuspended cells a few additional times to produce smaller cell aggregates. The volume of resuspended cell aggregates transferred to the new well was calculated to yield ∼20% confluency upon plating. The next day, cells were transfected with 1 µg Alt-R S.p. Cas9 Nuclease V3 (Integrated DNA Technologies), 250 ng gRNA (Synthego), and 125 ng CleanCap EGFP mRNA (TriLink Biotechnologies) using Lipofectamine Stem reagent (ThermoFisher #STEM00001) according to the manufacturer’s protocol. Cells were exposed to transfection complexes for 24 hours, then medium was refreshed. Two days after lipofection, cells were prepared for sorting by treating them with working medium supplemented with 10 µM RI for three hours prior to dissociation. To create a single-cell suspension, cells were washed with DPBS and treated with Accutase (STEMCELL Technologies #07920) at 37°C for 10 minutes, followed by pipetting several times to break cell aggregates into single cells. The dissociation reaction was halted using Dulbecco’s Modified Eagle Medium/Nutrient Mixture F-12 with L-glutamine and HEPES (Gibco #15630080), and cells were pelleted by centrifugation. The pellet was then resuspended in DPBS supplemented with 10 µM RI. Cells were stained with 1 µL TO-PRO-3 viability dye (ThermoFisher #R37170), filtered through a 35 µm cell strainer, and sorted under sterile conditions using a FACS Aria II cell sorter (BD Biosciences) equipped with a 100 µm nozzle at a pressure of 20 psi. The sorter was configured to sort live (TO-PRO-3-negative), GFP-positive, single iPSCs, depositing one cell into each well of a 96-well Matrigel-coated plate filled with 100 µL working medium supplemented with 10% CloneR (STEMCELL Technologies #05889). Plates filled with sorted cells were transferred to the 37°C cell culture incubator and left undisturbed for 48 hours after sorting. Cloning medium was refreshed two days after FACS, 25 µL of cloning medium were added to each well three days after FACS, and then starting on d4 post-FACS, medium changes using working medium were performed every other day. On d14 post-FACS, visible colonies were passaged using ReLeSR treatment (STEMCELL Technologies #100-0483) to consolidate them into new 96-well Matrigel-coated plates. Cells from each well were split into replicate plates, one for propagation and one for DNA extraction. The Quick-DNA 96 Kit (Zymo Research #D3012) was used to extract DNA from clonal iPSC lines growing in 96-well plates. Colonies were detached using ReLeSR as above and resuspended in genomic lysis buffer. Cell lysis, DNA isolation, washing, and final elution were performed according to the manufacturer’s protocol. PCR to amplify the *MAGEL2* c.1996 locus was performed on DNA from each recovered iPSC line using primers MAGEL2_G722_FOR (5’-ATCGGGAAGCTGAAGAAAAGCCGTGCAAATCCAGC-3’) and MAGEL2_G722_REV (5’-ATCCGACGGTAGTGTGAAGAGGCCCTGCATTCTCC-3’). PCR products were then used as templates for a second round of PCR with primers that uniquely barcoded each sample and added Illumina sequencing adapters. Cleanup, library preparation, and massively parallel sequencing was then performed as previously described^202,203^, except PhiX sequencing control DNA (Illumina) was spiked in to the final library at 15% and sequenced together with barcoded PCR product to ensure sufficient nucleotide diversity for sequencing. Demultiplexing of sequencing data based on sample of origin was performed by the Walk-Up Sequencing Platform at the Broad Institute, taking advantage of barcodes sequenced as index sequencing reads. Analysis of sequencing data identified iPSC lines having acquired heterozygous indels of interest affecting the length of the cytosine homopolymer within the *MAGEL2* coding sequence. To determine whether indels of interest occurred on the maternal or paternal haplotype, long-range PCR was performed on DNA from iPSC lines identified above using primers MAGEL2_G722_FOR2 (5’-GTGCCACACGAGATTCCAAC-3’) and XREV1T (5’-GTGGTAATCTATGTATCCTGCTCAATTCATTTCACAAAGCCAGCAC-3’). Resulting PCR products were cloned into plasmids using the Zero Blunt TOPO PCR Cloning Kit (ThermoFisher #450245), followed by transformation into NEB Stable Competent *E. coli* (New England Biolabs #C3040H) using the manufacturer’s heat shock protocol. Transformations were then plated on standard LB agar plates with kanamycin. Long-range PCRs were then performed using the same long-range PCR primers as above on individual colonies picked from the agar plates. Colony PCR products were cleaned up using AMPure XP magnetic beads (Agencourt #A63882) following the manufacturer’s protocol and Sanger sequenced using primers MAGEL2_G722_SEQ (5’- AAGAGGCCCTGCATTCTCC-3’) and XREV1T. Sequencing results linked each indel to one allele of a nearby single nucleotide variant in each iPSC line. Analysis of WGS data from PWS deletion lines revealed the inheritance status (maternal or paternal) of each allele at this locus. Thus, comparison of the Sanger data and the Whole-Genome Sequencing (WGS) data^204^ was performed to identify iPSC lines with paternal *MAGEL2* c.1996delC and c.1996dupC indels. WGS was performed for each starting, unedited iPSC line (two total) using a modified NEBNext Ultra II DNA Library Prep Kit for Illumina (New England BioLabs #E7645L). Briefly, 1 µg of DNA from the unedited iPSC line was fragmented to approximately 350 bp using Covaris shearing (Covaris #E220). DNA fragments were end repaired and A-tailed using Ultra II reagents. Illumina stubby-Y adapters were ligated to fragments and finished with PCR using 8-bp barcoded primers (Integrated DNA Technologies #10005921) using Ultra II reagents. Individual libraries were pooled and sequenced on NovaSeq S4 using paired-end 150 bp chemistry on the Walk-Up Sequencing Platform at the Broad Institute. In addition, *SmaI* digestion (New England Biolabs #R0141S) with subsequent PCR allowed for amplification of only the methylated maternal allele. Digested and undigested PCR products were sanger sequenced to confirm the paternal allele of point mutation, as described previously^8^. The intact methylation pattern was confirmed by methylation-sensitive multiplex ligation-dependent probe amplification, as also previously described^20^.

### Generation of CRISPR/Cas9-engineered *PWSTI* and *PWSTII* deletions

Using the SCORE strategy^205^, two isogenic hiPSC backgrounds (MGH2069 female^206^ and GM08330 male^207^) were used to create paternal deletions of the 15q11-q13 locus. Guide RNAs (gRNAs) were designed against segmental duplication blocks upstream and downstream of the canonical boundaries of PWS Type I and Type II deletions using the GRCh38 reference genome. Induced Cas9 double-strand breaks followed by imperfect repair were expected to create deletions in isogenic iPSCs that mimic clinical PWS alleles. gRNAs targeting the 5′ and 3′ boundaries of the Type I deletion were 5′-ATAGTAGCAAAACGCATACT-3′ (PAM: AGG) (chromosome 15:22,588,838–22,588,857) and chromosome 15:28,683,290–28,683,309, together with 5′-CGTGCACGTGTGGAAACTGT-3′ (PAM: AGG). gRNAs targeting the 5′ and 3′ boundaries of the Type II deletion were 5′-CTCCCTGCCTAGAAGCTGGT-3′ (PAM: TGG) (chromosome 15:23,198,458–23,198,477) and chromosome 15:28,490,195–28,490,214, together with 5′-CGAGAATAGTTTGTGTACTG-3′ (PAM: GGG). To increase specificity and efficiency, high-fidelity Cas9 was delivered as an RNP complex together with synthetic gRNAs^208^ into hiPSC, and single cells were isolated by FACS and clonally expanded. Individual clonal lines were screened by ddPCR for genomic copy number changes spanning the PWS critical region. Genes in the deleted interval (*NIPA1*, *HERC2*) and outside the interval (*APBA2*) were measured as an internal control, using Applied Biosystems TaqMan assays (*HERC2*, Hs03912010_cn; *NIPA1*, Hs03914881_cn; *APBA2*, Hs05365004_cn). To determine the parent-of-origin of the deleted allele, quantitative methylation analysis of bisulfite-treated DNA was performed in the SNURF–SNRPN exon 1/promoter region which is maternally methylated and paternally unmethylated. Primers were designed without CpG sites to avoid amplification bias (forward: 5′-GAG GGA GTT GGG ATT TTT GTA TTG-3′; reverse: 5′-CCC AAA CTA TCT CTT AAA AAA AAC CAC-3′), together with hydrolysis probes specific for either the methylated maternal allele (5′-(6FAM)-CGT TTG CGC GGT CG-(MGB)-3′) or unmethylated paternal allele (5′-(VIC/HEX)-AAG TAT GTT TGT GTG GTT GTA G-(MGB)-3′)^209^. The intact methylation pattern was confirmed by methylation-sensitive multiplex ligation-dependent probe amplification, as previously described^20^.

### Generation of CRISPR/Cas9-engineered heterozygous and homozygous *MAGEL2* deletions

The template sequence to generate the *MAGEL2* deletions was obtained from the UCSC Genome Browser. gRNAs flanking the *MAGEL2* gene were designed with Integrated DNA Technologies (IDT), ChopChop, and Benchling and ordered from IDT. Primer pairs capturing the gRNA cuttings were designed with NCBI Primer-BLAST, ordered from IDT, and tested on WT gDNA. The gRNA pairs (5’-caccGGACTCCCAAGACATCATAC-‘3 and 5’- aaacGTATGATGTCTTGGGAGTCC-‘3 as well as 5’-caccACCTCTAATCTGCCGCTAGG-‘3 and 5’- aaacCCTAGCGGCAGATTAGAGGT-‘3) were cloned into the pX330-hCas9-long-chimeric-grna-g2p plasmid (Addgene #42230) as previously described. ^210^ To confirm correct transformation, plasmids were sequenced to ensure correct integration of gRNAs. hiPSC (male GM08330 and female MGH2069) were transfected by non-liposomal formulation with FuGene HD transfection reagent (Promega #E2311) during cell passaging. Before starting the transfection, FuGENE was mixed with iPS-Brew XF without supplements, and the plasmid DNA and the FuGENE HD Transfection Reagent were incubated at RT for 15 min in a ratio of 100 µL mTeSR: 5 µL FuGENE HD: 1 µg plasmid DNA. After washing with DPBS, cells were incubated with Versene (ThermoFisher #15040066) for 7 min. After incubation, Versene was removed and the cells were resuspended in pre-warmed StemMACS™ iPS-Brew XF supplemented with 2 µM Thiazovivin. The cell suspension was added to the FuGENE mixes in a ratio of 900 µL cells: 100 µL mTeSR: 5 µL FuGENE HD: 1 µg plasmid DNA, inverted gently at room temperature for 5 min and then seeded into the Geltrex-coated wells. Medium was changed after 24 h to remove Thiazovivin. Transfection efficiency was evaluated via fluorescence microscopy 24 h post-transfection. The desired deletion was confirmed in the bulk population by PCR-based genotyping with the ALLin™ Taq DNA Polymerase (HighQu #PCE0101). Each 25 µL PCR reaction contained 1X ALLin Taq Mastermix, 0.4 µM of each primer (WT primer pair: 5’-ACTGAGATTATCACTCCTCCTCACC-3’ and 5’-TCTCACTGTGTCCAGTCCCC-3’; Deletion primer pair: 5’-ACTGAGATTATCACTCCTCCTCACC-3’ and 5’-GTGGGCTCCTCTCCTAGTCA-3’), and 100 ng of gDNA (at a final concentration of 4 ng/µL). Thermal cycling was performed with an initial denaturation at 95°C for 1 min; followed by 40 cycles of denaturation at 95°C for 15 s, annealing at 59.5°C for 15 s, and extension at 72°C for 15 s; and a final extension at 72°C for 3 min. The resulting products were confirmed by Sanger sequencing. To generate single-cell clones, the targeted bulk population was dissociated with Accutase and resuspended in StemMACS™ iPS-Brew XF supplemented with 2 µM Thiazovivin. A serial dilution was performed (∼1 cell per 100 µL) and cells were seeded on 96-well plates coated with Geltrex (1:33) with 100 µL of media. The medium was changed 2-3 days after seeding and following that, every other day. Once cell colonies reached confluency, they were expanded to 24-well plates. To isolate the gDNA, the cell suspension was transferred into a microcentrifuge tube and centrifuged at 8000 rpm for 3 minutes. The supernatant was aspirated and the cells were resuspended in 100-300 µL QuickExtract^TM^ DNA Extraction Solution (Biosearch Technologies #QE09050) containing 1 µL Proteinase K (600U/ml, ThermoFisher #EO0491) This solution was incubated at 65°C overnight and subsequently at 98°C for 5 min on the next day to inactivate Proteinase K, following PCR-based genotyping using the same primers as for the bulk population.

By switching the primer pairs, clones were identified containing an inverted and re-inserted *MAGEL2* gene and excluded from future use. With the use of heterozygous SNPs found in the MGH and 8330 WT (rs850823 G>A/C/T), amplified by the primer pair 5’-CAAATAACCAGGTGGTCCCG-‘3 and 5’-CGGTTGTTCAGCAACTTCAGG-‘3, the absence of mosaics of genetically different clones was confirmed. For the allele identification, gDNA was isolated with the Genomic DNA Purification Kit (STEMCELL Technologies #79020) according to the manufacturer’s instructions*. SmaI* digestion (New England Biolabs #R0141S) with subsequent PCR allowed for amplification of only the methylated maternal allele, resulting in the distinction between paternal and maternal heterozygous *MAGEL2* deletions^8^. For the detection of methylation status and copy number variations, methylation-sensitive multiplex ligation-dependent probe amplification (MS-MLPA) was performed, as previously described^20^. In brief, seven *MAGEL2*/*SNRPN* methylation-specific probes were utilized that are expected to show 50% methylation in WT hiPSC (0% on the paternal allele and 100% on the maternal allele). In case of paternal PWS / *MAGEL2* deletions, these probes are fully methylated (due to deletion of the paternal allele, resulting in 100% methylation from the maternal allele), whereas in samples with a deletion of the maternal allele, there is no methylation. MS-MLPA was applied according to the manufacturer’s instructions (MRC-Holland #ME028-D1). In hiPSC with heterozygous *MAGEL2* deletion, the remaining *MAGEL2* allele was fully methylated, indicating that this is the maternal allele. At the *SNRPN* locus, the methylation pattern was normal, confirming an isolated deletion excluding *SNRPN* or other potential imprinting defects in this region (Figure 2A). Karyotyping was done using the hPSC Genetic Analysis Kit (STEMCELL Technologies #07550). Clones were expanded to a 6-well plate and used for subsequent experiments.

### Generation of hiPSC-derived neurons

This protocol is an adaptation from Qi et al., 2017 ^41^. hiPSC grown to a confluency of 80% were dissociated into single cells with Accutase (8 min at 37°C), collected in a falcon, and plated with StemMACS™ iPS-Brew XF supplemented with 2 µM Thiazovivin at 2.5*10^5^ cells/cm^2^ or 2.4*10^6^ cells/well of a 6-well plate. The following day, the medium was completely aspirated, and cells were washed once with DPBS. The differentiation was started in 5 mL of KSR medium per well (84% DMEM/F-12, GlutaMAX™ Supplement (ThermoFisher #31331093), 14% KnockOut™ Serum Replacement (ThermoFisher #10828010), 1% MEM Non-Essential Amino Acids Solution (ThermoFisher #11140050), and 50 mM β-Mercaptoethanol (Sigma Aldrich #M3148)). Medium was changed daily. Starting from d4, this was gradually switched to N2 medium (95% Neurobasal™ Medium (ThermoFisher #21103049), 1% GlutaMAX™ Supplement (ThermoFisher #35050087), 1% MEM-NAC, 1% N-2 supplement (ThermoFisher #17502048), 2% B-27™ Supplement (ThermoFisher #17504044). From d0 until d5, the medium was supplemented with 250 nM LDN193189 (Sigma Aldrich #SML0559), 10 µM SB431542 (STEMCELL Technologies #72234), and 5 µM XAV939 (Tocris #3748). In addition, from d2 until d7, further medium supplementation included 1 µM PD03259013 (STEMCELL Technologies #72184), 5 µM SU5402 (Sigma Aldrich #SML0443), and 10 µM DAPT (Tocris #2634). To passage the cells on d8, the medium was aspirated, and the cells were washed once with DPBS, following dissociation into single cells with Accutase (45 min at 37°C). After collection in a falcon and centrifugation at 200g for 5 min, cells were counted and resuspended in BCA medium (97% Neurobasal™ Plus Medium (ThermoFisher #A3582901), 1% GlutaMAX™ Supplement (ThermoFisher #35050087), and 2% B-27™ Plus Supplement (ThermoFisher #A3582801)) supplemented with 8 µM PD0325901, 10 µM SU5402, and 3 µM CHIR99021 (Selleckchem #S2924) until d16. Further supplementation with 10 µM DAPT, 20 ng/ml BDNF (PreproTech #450-02), 10µg/ml GDNF (PreproTech #450-10), and 200 µM ascorbic acid (Sigma Aldrich #A4544) was continued until d30. Finally, the cells were plated at 2,5*10^5^ cells/cm^2^ on plates pre-coated with Poly-L-Ornithine Solution (diluted 1:5 in water; Sigma Aldrich #A-004-C) followed by Geltrex™ LDEV-Free diluted 1:50 in KnockOut™ DMEM. 2 µM of Thiazovivin was added to the medium which was replaced the next day followed by medium change every 3-4 days in the remaining differentiation. Cells were harvested on d16 or d30 of differentiation.

### RNA isolation, DNase treatment, cDNA synthesis, and RT-qPCR

Total RNA was isolated using the RNAqueous™-Micro Total RNA Isolation Kit (ThermoFisher #AM1931) according to the manufacturer’s instructions. RNA was first DNase-treated (ThermoFisher #EN0521) and then transcript into cDNA with the ProtoScript® II First Strand cDNA Synthesis Kit (NEB #E6560L) according to the manufacturer’s instructions. RT-qPCRs were performed on the QuantStudio 5 (ThermoFisher #A28574) using the ORA qPCR Green ROX L Mix (highQu #QPD0101) following the system’s run mode “Fast” protocol. cDNA was diluted 1:5 and custom *MAGEL2* RT-qPCR primers (5’-CAACCGTGCCAACAATAAGC-’3 and 5’-ATAGGATGCCACCAAATTCCC-‘3), Integrated DNA Technologies) were used with a final concentration of 0.25 µM. Gene expression fold changes were normalized to the housekeeping genes *ACTB* and *GAPDH* calculated using the ΔΔCt method.

### Pull-down for ubiquitinomics

Lysis buffer (25 mM Tris-HCl pH 7.5, 150 mM NaCl, 1% NP40 alternative (Sigma-Aldrich #492016), 1 mM EDTA (ThermoFisher #J15700.C4), 5% Glycerol, 100 mM NEM (Sigma-Aldrich #E3876)) was supplemented with a Halt™ Protease & Phosphatase Inhibitor Cocktail (1x, ThermoFisher #1861281, #78442). Cells were removed from the incubator and immediately put on an ice-water mix for lysis. Cells were washed once with ice-cold DPBS. Then, ice-cold lysis buffer was added to the cells and incubated for 5 min on ice. Lysates were then stirred with a cell-scratcher and incubated for another 10 min on ice. Lysates were transferred to a tube, resuspended thoroughly, and incubated for one hour on ice and then centrifuged at 13100g at 4°C. Supernatants were transferred to a new tube, and debris was resuspended in lysis buffer for storage. Samples were flash-frozen in liquid nitrogen and stored at -80°C. For the ubiquitin enrichment, the Pierce™ Ubiquitin Enrichment Kit (ThermoFisher #89899) was used. Lysates were thawed on ice and centrifuged for 15 min. at 13100g at 4°C to reduce debris contamination further. Bicinchoninic Acid Assay (ThermoFisher #23225) was performed with supernatants before enrichment. 150 µg protein was loaded into the column and diluted 1:1 with Tris-buffered saline (TBS) buffer (pH 7.2) provided by the kit. 20 µL of the anti-polyubiquitin (anti-poly-Ub) resin was added to the sample to precipitate polyubiquitinated (poly-Ub) proteins. Samples were incubated overnight on a rotator at 4°C. Unbound poly-Ub proteins were centrifuged at RT at 5000g for 15 sec. Samples were washed 3x with 300 µL wash buffer by centrifugation at RT at 5000g for 15 sec. Wash buffer consisted of a 1:10 dilution of lysis buffer in TBS. Wash fractions were stored for analysis. After the third wash, samples were eluted with 1x Laemmli buffer (diluted with TBS, BioRad #1610747) supplemented with 355 mM BME by denaturing at 95°C for 5 min. Eluates were collected into tubes by centrifugation at 5000g for 30 sec. 20 µL of the eluates were used for western blot. Samples from unbound and wash fractions were diluted in a ratio of 3:1 with 4x Laemmli buffer supplemented with 10% BME and denatured at 95°C for 5 min. Denatured samples were loaded onto 4-12% or 10% Bis-Tris SDS-polyacrylamide gels (ThermoFisher #NP0302 and #NP0322) with MES SDS running buffer for SDS-PAGE (ThermoFisher #NP0002) at 120 V for 1:30 h. Polyvinylidene fluoride (PVDF) membranes (ThermoFisher #88518) were washed once with PBS and blocked with 5% BSA-PBST for 1 h at RT on a rocking table. The anti-poly-Ub antibody was diluted 1:7500 in the respective blocking buffer and incubated overnight at 4°C on a rocking table. Membranes were washed three times with PBST, then incubated for one hour at RT on a rocking table with 1:5000 diluted anti-Rabbit (H+L) IgG monoclonal antibody HRP-conjugate (BioRad #1706515) in respective blocking buffer. Membranes were washed 3 × 5 min. with PBST and 1 × 2 min. with PBS. Membranes were transferred to clear film, and 1 mL of Clarity™ Western ECL Substrate (BioRad #170-5060) 1:1 reaction mixture was dribbled over the membrane. The clear film was closed, and the chemiluminescence reaction was allowed to react for 5 minutes in the dark at RT. Imaging was performed by ChemoStar Touch ECL & Fluorescence Imager.

### RNA-seq

Input RNA was initially QC’d with the Agilent TapeStation, using the RNA 6000 Assay to check for integrity, and quantified with the Qubit BR RNA Assay from Invitrogen. 240ng of Total RNA were then processed with the Takara SMARTer Stranded Total RNA Prep (H/M/R) to generate sequencing libraries compatible with the Illumina Sequencing Instruments. Sequencing was performed on a P3 100 cycle flowcell with the NextSeq 2000 .

### RNAseq preprocessing

Good quality of all sequencing data was ensured by quality control using FastQC (0.11.9)^165^. Reads were then aligned to the hg38 reference genome (refdata_gex_GRCh38-2020-A) employing BWA^166^, a local aligner with soft clipping, in mode *bwa mem*. Resulting *SAM* files were converted to *BAM* files through the SAMtools (1.15.1)^167^ view -bS call. A gene x sample count matrix was generated in R (version 4.3.1) using *featureCounts()* from the Rsubread package^188^ with the in-built hg38 annotation in paired-end mode. Comprehensive QC metrics were generated using the RNAseqQC package.^168^ We evaluated total library sizes, library complexity (fraction of genes detected), the number of detected genes per sample, and the distribution of gene biotypes to ensure sample consistency and data integrity

### Protein Extraction and Digestion for Proteomics

Whole-cell proteome pellets were lysed in 0.1% (w/v) RapiGest SF (Waters, #186001860) prepared in 50 mM ammonium bicarbonate buffer (Sigma-Aldrich) and incubated at 80 °C for 15 min. Lysates were subsequently sonicated using a Bioruptor system (Diagenode) for 15 cycles of 30 s ON/30 s OFF at 10 °C. Protein disulfide bonds were reduced by incubation with 5 mM dithiothreitol (DTT) at 60 °C for 15 min, followed by alkylation with 10 mM iodoacetamide for 30 min at room temperature in the dark. Proteins were digested overnight at 37 °C using sequencing-grade modified trypsin (Promega) at 1:40 an enzyme-to-protein ratio appropriate for complete digestion. Proteolytic digestion was quenched by acidification with trifluoroacetic acid (TFA) to a final concentration of 0.2% (v/v), and RapiGest was precipitated by further incubation at 37 °C for 40 min. Samples were centrifuged at 12,000 rpm for 20 min at 4 °C, and the clarified supernatants containing tryptic peptides were collected. Peptides were desalted using self-packed C18 StageTips containing four discs of Empore C18 material (Merck Supelco). StageTips were sequentially conditioned with 100% acetonitrile (ACN; Biosolve Chimie), solvent B (50% (v/v) ACN and 0.1% (v/v) formic acid (FA) in ddH₂O), and solvent A (0.1% (v/v) FA in ddH₂O). After loading, peptides were washed and eluted according to standard protocols. Peptide concentrations were determined using the bicinchoninic acid (BCA) assay (Pierce, Thermo Scientific). Purified peptide samples were stored at −80 °C until further mass spectrometric analysis.

### Ubiquitinome Sample Preparation

Proteins eluted from immunoprecipitation (IP) were diluted in 50 mM triethylammonium bicarbonate (TEAB; pH 8.5; Sigma-Aldrich) supplemented with 1% (v/v) sodium dodecyl sulfate (SDS; Bio-Rad). Samples were denatured at 95 °C with agitation at 600 rpm for 10 min. Subsequently, trifluoroacetic acid (TFA; Biosolve Chimie) was added to a final concentration of 2% (v/v), and the reaction was quenched after 1 min by neutralization with 3 M tris(hydroxymethyl)aminomethane (Tris; AppliChem). Due to limited sample input, protein digestion and peptide cleanup were performed using an automated single-pot solid-phase–enhanced sample preparation (SP3) workflow implemented on a Bravo liquid-handling platform (Agilent), enabling parallel processing of up to 96 samples. The platform performs protein reduction, alkylation, magnetic bead handling, SP3-based protein cleanup, on-bead digestion, and peptide recovery. Sera-Mag SpeedBeads magnetic carboxylate particles (Fisher Scientific) were washed and resuspended to a working concentration of 100 μg/μL. For protein binding, 20 μL of 100% acetonitrile (ACN) was added to each sample, followed by incubation off the magnetic rack for 18 min with alternating agitation cycles (1,500 rpm for 30 s and 100 rpm for 90 s). Samples were then placed on a magnetic rack (UNIEQUIP MAGNUM FLX Enhanced Universal Magnet) for 5 min to capture bead-bound proteins. Bound proteins were reduced and alkylated directly on beads using 150 μL of a solution containing 10 mM tris(2-carboxyethyl)phosphine (TCEP) and 40 mM 2-chloroacetamide (CAA) in 100 mM Tris-HCl (pH 8.0). Following washing steps, beads were resuspended in 50 μL of 100 mM ammonium bicarbonate (pH 8.0). Proteolytic digestion was performed on-bead by addition of 2 μL sequencing-grade trypsin (Promega; 0.5 μg/μL in 50 mM acetic acid) and incubation for 16 h at 37 °C in a thermal shaker. Plates were sealed throughout the procedure using VersiCap 96-well flat cap strips. Following digestion, peptide solutions were transferred to a fresh 96-well plate and concentrated using a UNIVAPO-150H vacuum concentrator connected to a UNICRYO MC2 cooling trap and a UNITHERM 4/14 D closed-circuit cooler (UNIEQUIP). For peptide cleanup, Sera-Mag SpeedBeads were diluted to 100 μg/μL in 10% (v/v) formic acid, and 5 μL of this suspension was added to each dried sample. Peptides were bound to beads by addition of 195 μL ACN and incubation for 18 min with orbital shaking at 100 rpm. After magnetic separation, the supernatant was removed in two steps, and beads were washed twice with 180 μL ACN and air-dried. Peptides were eluted by resuspension of beads in 20 μL of 0.1% (v/v) formic acid in water, followed by sonication for 10 min in an ultrasonic cleaner (USC-T; VWR). The eluates were transferred to a fresh plate, diluted in 0.1% formic acid, and used for LC–MS/MS analysis. Peptide concentrations were determined using the bicinchoninic acid (BCA) assay (Pierce, Thermo Scientific).

### Data Acquisition (LC–MS/MS)

Peptide samples (400 ng) were injected in a volume of 1 µL onto an analytical column (IonOpticks Aurora Series, 25 cm × 75 µm i.d., CSI, 1.6 µm C18) using an EASY-nLC 1200 system (Thermo Fisher Scientific). Peptides were separated at a flow rate of 250 nL/min with the column temperature maintained at 50 °C using a column oven (Sonation). Sample loading was performed at a maximum flow rate of 1 µL/min and a maximum pressure of 980 bar. For full proteome analysis, peptides were separated using a 120 min active gradient starting at 4.0% (v/v) acetonitrile (ACN; Biosolve Chimie) in ddH₂O containing 0.1% (v/v) formic acid (FA), increasing to 17.0% ACN over 88 min, to 25.0% ACN over 104 min, and to 35.0% ACN over 120 min. For ubiquitinome analysis, peptides were separated using a 90 min active gradient starting at 4.0% ACN with 0.1% FA, increasing to 17.0% ACN over 58 min, to 25.0% ACN over 74 min, and to 35.0% ACN over 90 min. Mass spectrometric data were acquired on a timsTOF Pro mass spectrometer (Bruker Daltonics) in data-independent acquisition parallel accumulation–serial fragmentation (diaPASEF) mode using timsControl software. Ions were accumulated in the trapped ion mobility spectrometry (TIMS) analyzer at a constant accumulation potential corresponding to an inverse reduced mobility (1/K₀) of 1.5 V·s/cm². Mobility separation was achieved by linear ramping from 1.25 to 0.65 V·s/cm² over 100 ms with a locked duty cycle of 100%. The TIMS ramp rate was set to 9.51 Hz, and ion charge control was disabled. Full MS1 scans were acquired over an m/z range of 100–1,700. For diaPASEF acquisition, precursor ions within an m/z range of 375–925 and an ion mobility range of 0.66–1.22 1/K₀ were selected for fragmentation. This 550 m/z precursor mass range was partitioned into isolation windows of 25 Th width and approximately 0.15 V·s/cm² ion mobility height. The windows were distributed across 8 diaPASEF MS/MS scans per cycle, resulting in a total cycle time of approximately 0.95 s. Fragmentation was performed in the collision cell using ion mobility–dependent collision energies, linearly ramped from 45 eV at 1/K₀ = 1.30 V·s/cm² to 27 eV at 1/K₀ = 0.75 V·s/cm². Collision RF was set to 1500 Vpp. The quadrupole ion energy was set to 5.0 eV with a low-mass cutoff of 200 m/z. Transfer optics were operated with Funnel 1 RF at 350 Vpp, Funnel 2 RF at 200 Vpp, multipole RF at 200 Vpp, and a deflection delta of 70 V. The TOF analyzer was operated with a transfer time of 60 µs and a pre-pulse storage time of 12 µs.

### Raw Data Processing

Raw diaPASEF data were processed using a library-free data-independent acquisition (DIA) workflow in Spectronaut (version 18; Biognosys). Peptide identification was performed using an in silico–generated spectral library derived from the UniProtKB reference proteome of *Homo sapiens* (downloaded September 2021; 20,386 reviewed protein entries). Trypsin/P was specified as the proteolytic enzyme, allowing up to two missed cleavages per peptide. Peptide length was restricted to 7–50 amino acids, and precursor charge states were limited to 2–4. Carbamidomethylation of cysteine residues was set as a fixed modification, while methionine oxidation and protein N-terminal acetylation were specified as variable modifications.

Protein inference was performed by grouping protein isoforms according to their corresponding protein names as defined in the FASTA file. Precursor-level quantification was performed using the “any LC (high precision)” quantification strategy. The match-between-runs (MBR) feature was enabled for all experiments to improve identification consistency across samples. False discovery rate (FDR) control was applied at 1% at the precursor level using Spectronaut’s default target–decoy approach. Protein-level FDR filtering was applied according to default Spectronaut settings. All remaining parameters were kept at default values unless stated otherwise.

### Principal Component Analysis

All computational analyses were conducted using R (version 4.3.1) with the dplyr^174^ and tidyverse^172^ packages for data manipulation. The human genome annotation database org.Hs.eg.db (version 3.17.0) and transcriptome database TxDb.Hsapiens.UCSC.hg38.known-Gene were used for all gene-level annotations interfaced using the AnnotationDbi^190^ and GenomicFeatures^191^ packages. To visualize the relationships between samples and identify the primary sources of variation within the transcriptomics, proteomics, and ubiquitinomics data, we performed principal component analyses (PCAs) and generated plots using ggplot2.^171^ Data cleaning and string manipulations were performed using stringr.^175^ For transcriptomics, raw count data was loaded into a DESeqDataSet object. To normalize for differences in library size and stabilize variance, we first applied a variance-stabilizing transformation (vst) using the DESeq2 package^187^, with the blind = TRUE setting. PCA was then performed using the R prcomp function from the stats package^211^ on the transposed, scaled (scale. = TRUE) matrix of transformed expression values. For proteomics and ubiquitinomics, data were managed using the SummarizedExperiment package^189^ and unified to enable direct comparison of samples across backgrounds. Missing values were characterized as Missing Not At Random (MNAR), attributed to protein abundances falling below the limit of detection. Accordingly, missing data were imputed using the Probabilistic Minimum Imputation (MinProb) method from the DEP2 package. For each data type, the imputed abundance matrices rom the MGH and 8330 backgrounds were filtered to retain only the commonly detected features, which were defined as the intersection of features present in both backgrounds. These filtered matrices were then concatenated into a single, combined data matrix. PCA was subsequently performed on this unified, transposed, and scaled (scale. = TRUE) matrix using the prcomp function. Marginal density distributions were visualized with the ggExtra package.^176^

### Identification of Differentially Expressed Genes and Proteins

To identify differentially expressed genes (DEGs), differentially expressed proteins (DEPs), and differentially expressed ubiquitinated proteins (DEUs) we compared each of the six genetic conditions (DelC, DupC, HetDel, HomoDel, PWSTI, PWSTII) to WT controls. For the RNA-seq data, DEG lists were generated from the DESeq2 result objects. Genes were considered significantly dysregulated if they had an adjusted p-value (p.adj) calculated using the Benjamini-Hochberg (BH) method of less than 0.05. These genes were then further stratified into upregulated (log2fold-change>0) or downregulated (log2fold-change<0) sets for each comparison. For the proteomics data, differential expression analysis was performed using the DEP2 package^192^. Proteins were deemed significantly dysregulated if their corresponding p.adj (BH) was less than 0.05. These significant DEPs were then categorized as upregulated (log2fold-change>0) or downregulated (log2fold-change<0) for downstream analysis. For the ubiquitinomics data, differentially expressed ubiquitinated proteins (DEUs) were identified from the differential expression results tables similar to the proteomics data. Given the higher technical variability of pull-down-based ubiquitinomics^143^, a relaxed statistical threshold (FDR < 0.1) was used to identify differentially ubiquitinated proteins (DEUs) relative to their isogenic WT. These significant DEUs were then categorized as upregulated (log2fold-change>0) or downregulated (log2fold-change<0) for downstream analysis. This process was performed independently for both the MGH and 8330 cell line backgrounds, resulting in distinct sets of up- and downregulated DEGs, DEPs, and DEUs for each experimental condition and background.

### Cell type Enrichment Analysis

To identify cell-type-specific signatures within our differential expression results, we performed an over-representation analysis using the clusterProfiler R package.^193^ We tested for enrichment against the MSigDB C8 cell type signature gene set collection, retrieved using the msigdbr package.^194^ Similar to the GO enrichment, we performed the analysis on the significantly upregulated (p.adj < 0.05, log2FoldChange > 0) and downregulated (p.adj < 0.05, log2FoldChange < 0) genes for each genotype in each background separately. The enricher function was used with gene symbols to calculate enrichment significance based on a hypergeometric test, and p-values were adjusted for multiple comparisons using the BH method with a p.adj cutoff of 0.05. For visualization, results from all conditions were combined into a single data frame. We then generated dot plots by filtering this combined data for specific subsets of interest, including gene sets from the ZHONG_PFC and FAN_EMBRYONIC_CTX datasets, as well as a final curated list of key neurodevelopmental cell types.

### Disease Database Enrichment Analysis

This analysis was adapted from the approach described by Gordon et al., 2026.^161^ To identify high-confidence molecular signatures robust to genetic background effects, we intersected the DEGs from both backgrounds for each genotype and only included DEGs with the same direction of change (upregulated in both or downregulated in both). Additionally, an overlapping list was generated by intersecting the DEGs across all genotypes. Next, we compiled a panel of six curated gene sets:

1. SFARI ASD database including syndromic, high and strong confidence genes (release April 3, 2025).^212^
2. SPARK ASD database (release September 2022).^212^
3. High Confidence NDD genes obtained from GeneTrek (release April 26, 2024).^213^
4. Systemic Intellectual Disability genes from the SysID database.^213^
5. ASD risk genes identified via large-scale exome sequencing.^214^
6. ASD risk genes identified via large-scale whole genome sequencing.^215^
7. EPPAN epilepsy genes (Mayo Clinic laboratories) obtained from GeneTrek (release April 26, 2024)^213^

A one-tailed hypergeometric test was performed to determine the statistical significance of the overlap between our overlapping DEG list, and the disease gene lists, with the universe being defined as all detected genes in the RNA-seq dataset (>10 counts).

### Functional Enrichment and Gene Ontology (GO) Analysis

To elucidate the biological functions associated with the observed transcriptomic and proteomic changes, we performed Gene Ontology (GO) enrichment analysis on each set of significant DEGs, DEPs and DEUs. Prior to analysis, gene symbols were converted to Entrez IDs using the bitr function from the clusterProfiler package^193^. The enrichment analysis was conducted using the enrichGO function, testing against the "all" ontology (encompassing Biological Process, Cellular Component, and Molecular Function). We used the BH method for multiple testing correction. GO terms were considered significantly enriched if they had a p.adj of less than 0.05. The resulting lists of significantly enriched GO term descriptions were extracted for each comparison, creating separate lists for the upregulated and downregulated genes/proteins in both the MGH and 8330 backgrounds. These lists formed the basis for subsequent overlap and pathway-level comparisons.

### Cross-Background and Inter-Condition Overlap Analysis

To identify robust molecular signatures that are independent of the cellular background, we quantified the overlap of DEGs, DEPs and DEUs between the MGH and 8330 backgrounds for each genetic condition. Overlaps for upregulated, downregulated, and the union of all dysregulated genes/proteins were calculated and visualized as area-proportional Euler diagrams using the eulerr package^178^. The statistical significance of the overlap between the two backgrounds for any given condition was assessed using the one-tailed hypergeometric test using the phyper function^211^ with the universe defined as all genes/proteins detected in the experiment. An overlap was considered statistically significant if the resulting p-value was less than 0.05. To investigate the relationships between the molecular signatures of the different genetic conditions, we first defined a core set of commonly dysregulated genes/proteins for each condition by taking the intersection of the MGH and 8330 DEG/DEP/DEU lists. We then constructed an overlap matrix quantifying the number of shared DEGs, DEPs and DEUs between every pair of genetic conditions. These matrices were visualized as heatmaps using the pheatmap package^177^.

### Comparative Functional Analysis and Intersection Visualization

To compare the functional consequences of genetic perturbations across different conditions and data modalities, we analyzed the intersections of both dysregulated molecules and their associated enriched GO terms. First, we overlapped the enriched GO terms between the MGH and 8330 backgrounds for each condition. The overlapping GO terms were visualized in UpSet plots generated via the UpSetR^184^ and Complex Upset^170^ packages to compare the intersections across variants. To prioritize enriched GO terms for specific intersections of interest (e.g., genotype-exclusive or shared pathways), GO terms were ranked based on the significance of their enrichment across the relevant experimental contrasts. For each UpSet intersection, we calculated the geometric mean of the BH p.adj from each comparison of that intersection and used those to visualize the top 10 most significant terms per intersection.

### Analysis of PWS-Exclusive Signatures

To distinguish between the direct effects of gene loss within the PWS locus and the subsequent downstream consequences, we performed a filtered enrichment analysis. All genes residing within the canonical PWS locus on chromosome 15 (hg38, chr15:22,480,000-28,500,000) were identified using the TxDb.Hsapiens.UCSC.hg38.-knownGene^216^ and biomaRt^173^ packages. These locus-residing genes were then subtracted from the PWS-specific DEG list. GO enrichment analysis was subsequently re-run on this filtered, non-locus DEG set to identify biological pathways specifically dysregulated as a downstream effect of the PWS-like deletions.

### Identification and Characterization of Shared Proteomics Signatures

A core set of consistently up- and downregulated proteins common to all six variants was identified by taking the intersection of the respective DEP lists. This overlapping protein signature was then subjected to further functional characterization. In addition to GO enrichment, we also performed enrichment analysis for disease-associated phenotypes, using the gprofiler2 package, querying the Human Phenotype Ontology (HP) database.^180^ Terms with a BH p.adj < 0.05 were considered significant.

### Curation and Visualization of a Consensus MAGEL2 Interactor List

We first compiled a comprehensive list of putative MAGEL2-interacting proteins from several sources. This included affinity purification-mass spectrometry (AP-MS) and BioID proximity labeling experiments performed with different MAGEL2 constructs under basal and stress conditions^23,30,31,34^. To understand the relationships between these diverse ten datasets, the overlap and uniqueness of proteins identified in each experiment were visualized using an UpSet plot, generated with the UpSetR package.^184^ We then defined a consensus list of interactors by selecting the most stringent interactors reported in each individual study: The basal condition from Sanderson et al^31^ (34 proteins), the co-IP interactors from Heimdörfer et al.^30^ (91 proteins) and all interactors from Hao et al.^24^ (6 proteins) and Wijesuriya et al.^34^ (6 proteins). This curated list of 130 unique proteins formed the basis for all subsequent network analyses.

### Protein-Protein Interaction (PPI) Network Construction and Analysis

To explore the functional landscape of the MAGEL2 interactome, we constructed a protein-protein interaction (PPI) network. Using the curated consensus list of 130 MAGEL2 interactors, known interactions were retrieved from the STRING database (v11.5) via the STRINGdb package^195^. To ensure high fidelity, only interactions with a confidence score > 700 were included in the final network. The resulting network was analyzed using the igraph package^186^. Densely connected functional modules, or communities, were identified using the fast-greedy clustering algorithm (cluster_fast_greedy). The full network and its constituent community subgraphs were visualized using the Kamada-Kawai layout algorithm to spatially resolve functional clusters. Of the initial 130 proteins, 103 proteins demonstrated at least one interaction at the specified confidence score and were incorporated into the network.

### Integration of Proteomic Expression Data onto the PPI Network

To determine how the MAGEL2 interactome is affected by the genetic perturbations studied in this work, we integrated our differential proteomics data with the network structure. An "Expression Score" was calculated for each protein in the network. This score quantifies the overall direction and consistency of a protein’s change across all experiments. For each of the twelve comparisons (6 variants × 2 backgrounds), a protein was assigned a +1 if significantly upregulated (p.adj < 0.05) or a -1 if significantly downregulated (p.adj < 0.05). The Expression Score is the sum of these values, providing a metric where a large positive score indicates consistent upregulation and a large negative score indicates consistent downregulation across the panel of perturbations. This score was then mapped as a color gradient onto the nodes of the PPI network communities.

### Global Comparison of the Proteome and Ubiquitinome Data

This analysis was performed on the normalized abundance data (prior to differential expression calculation). First, the proteome and ubiquitinome data were filtered to retain only proteins detected in both experiments. For each experimental condition and background, the mean abundance was calculated across biological replicates for every protein. A stoichiometry ratio (Mean Ubiquitinome Abundance / Mean Proteome Abundance) was then calculated for each protein, providing an estimate of the fraction of that protein pool that is ubiquitylated. The genome-wide distribution of these stoichiometry ratios was visualized for each condition using ridgeline plots, generated with the ggridges package^179^.

### Comparative Analysis of Differential Proteome and Ubiquitinome Abundance

We first aimed to identify proteins where changes in ubiquitinome were decoupled from changes in the proteome. For each genotype vs the WT, the Log2 Fold Change and p.adj for each protein were extracted from both the proteomics and ubiquitinomics data for both genetic backgrounds. These data were then merged into a single comprehensive table and filtered for proteins that exists in all proteome and ubiquitinome data sets. The results from the MGH background were visualized as scatter plots, with the Log2 Fold Change from the total proteome on the x-axis and the Log2 Fold Change from the ubiquitinome on the y-axis. Then, for each genotype, we colored all background-overlapping DEUs in dark grey and selected DEUs shared across variants in red.

### Analysis of Post-Transcriptional Regulation

First, we assessed the global relationship between protein and mRNA levels across all conditions. RNA-seq counts were normalized for library size and variance using a variance-stabilizing transformation (vst), while imputed and normalized log2-intensity values were used for the proteomics data. To enable direct comparison, a mapping table was created to link each proteomics sample to its corresponding RNA-seq sample based on background and genotype. For each gene commonly detected in a matched sample pair, we calculated the log2-transformed ratio of protein to RNA abundance (log2(Protein) - log2(RNA)). The genome-wide distribution of these ratios for each experimental condition was then visualized using ridgeline plots generated with the ggridges package. To identify genes likely subject to post-transcriptional changes, we first created a master data table integrating the differential expression results (log2 fold-change and p.adj) from both the RNA-seq and proteomics datasets for all 12 comparisons (6 variants × 2 backgrounds) and filtered for genes/proteins detected in all RNAseq and proteomics datasets. The results from the MGH background were visualized as scatter plots using ggplot2, with the Log2 Fold Change from the transcriptome on the x-axis and the Log2 Fold Change from the proteome on the y-axis. Genes were then categorized into two "buckets" based on discordant regulation: Bucket 1 (Post-transcriptional Upregulation): This set included genes with a significant increase in protein abundance (Protein_p.adj < 0.05, Protein_Log2FC > 0) that was not accompanied by a corresponding significant increase in mRNA abundance (mRNA_p.adj > 0.05). Bucket 2 (Post-transcriptional Downregulation): This set included genes with a significant decrease in protein abundance (Protein_p.adj < 0.05, Protein_Log2FC < 0) that was not accompanied by a corresponding significant decrease in mRNA abundance (mRNA_p.adj > 0.05). For robust downstream analysis, a high-confidence foreground gene set was created for each bucket by taking the intersection of genes that met the criteria across all 12 experimental comparisons. The background gene set for all enrichment analyses was defined as all genes for which protein differential expression data were available in the master table. Next, we performed motif enrichment analysis for both RNA-Binding Proteins (RBPs) and microRNAs (miRNAs). For both foreground and background gene sets, 3’ UTR sequences were retrieved from the Ensembl database using the biomaRt package^173^. A curated, high-confidence human RBP motif database was first prepared. The full MotifDb was filtered to include only motifs corresponding to proteins annotated as RBPs in the RBP2GO database (https://rbp2go.dkfz.de, retrieved on September 3, 2025) with a confidence score ≥ 10. The foreground and background 3’ UTR sequences were then scanned for matches to this curated motif set using the motifmatchr package^196^. The statistical significance of the enrichment for each RBP motif in the foreground set relative to the background was determined using a one-tailed Fisher’s exact test, with p.adj corrected using the BH method. The background set was defined as all genes for which protein differential expression data were available. For the miRNA Target Site Enrichment, a comprehensive list of miRNA 8mer seed motifs was extracted from the targetscan.Hs.eg.db package.^198^ The 3’ UTR sequences were scanned for exact matches to these seed motifs using the Biostrings package^197^. As with the RBP analysis, enrichment significance was calculated using a one-tailed Fisher’s exact test with BH correction. For both analyses, enriched motifs (p.adj < 0.05) were reported, and the top 15 results were visualized as dot plots. To visualize the overlap and uniqueness of elements between different sets, we generated area-proportional Euler diagrams using the R package eulerr^178^. The sets of significantly enriched RBP motifs and miRNA families identified in "Bucket 1" (post-transcriptional upregulation) and "Bucket 2" (post-transcriptional downregulation) were compared to identify common and distinct regulatory signatures. The list of significantly enriched RBPs was intersected with a curated list of known MAGEL2 interactors to visualize the overlap between statistically identified regulators and known binding partners. The statistical significance of the overlap between any two sets was determined using a one-tailed hypergeometric test with the phyper function from the stats package^211^, with the universe defined as the total pool of items from which the sets were drawn. An overlap was considered statistically significant if the resulting p-value was less than 0.05.

### MOFA Training

To integrate transcriptomic, proteomic, and ubiquitination datasets, we utilized Multi-Omics Factor Analysis 2 (MOFA2).^109^ Data preparation and integration were performed in R Studio using a custom pipeline. Raw counts from RNA-seq and intensity values from proteomics and ubiquitinomics were processed separately for two genetic backgrounds (MGH and 8330). Transcriptomic data were normalized using the Variance Stabilizing Transformation (vst) from the DESeq2 package^187^. Proteomic and ubiquitination data were Z-score normalized by feature to ensure comparability across modalities and to minimize the impact of differing dynamic ranges. To handle technical replicates across backgrounds, feature values were averaged across corresponding samples, only if the feature was detected in at least two replicates per dataset. To prevent the model from being dominated by the higher dimensionality of the transcriptomic data, we implemented a feature selection strategy as suggested by Argelaguet et al., 2018. RNA-seq data were filtered to retain the top 9,000 most variable genes (HVGs) based on variance. For the final integrated model, RNA features were further subset to match the total number of features in the proteomic dataset, ensuring balanced weight across the different omics views. Features were uniquely labeled with their respective modality suffix (e.g., _RNA, _Protein, _Ubiq) to prevent naming conflicts. The MOFA input was constructed by merging long-format dataframes, and the model was trained using the create_mofa and run_mofa functions from the MOFA2 package. Missing values were handled using the framework’s built-in unsupervised latent factor inference.

### MOFA downstream analysis

The pre-trained MOFA model object was further analyzed using the MOFA2 package. The contribution of each latent factor to the total variation in the dataset was quantified using the plot_variance_explained function, assessing variance both per data view and per sample group. To visualize the relationships between samples in the low-dimensional space, we employed two methods. First, the values for each of the top latent factors were plotted for all samples, grouped by genotype. Second, we performed Uniform Manifold Approximation and Projection (UMAP) on the full set of inferred factors using the run_umap function from the MOFA2 package to generate a two-dimensional embedding of all samples. In this UMAP representation, samples were colored by genotype and shaped by cellular background to reveal the primary drivers of sample clustering. The independence of the inferred factors was confirmed by visualizing their pairwise correlations. To interpret the biological processes captured by each latent factor, we analyzed the feature weights and performed systematic pathway enrichment analyses. For each factor, the features (genes, proteins, or ubiquitinated proteins) driving the variation were identified by their corresponding weights. The top 10 features with the largest positive and negative weights were visualized for each data view. To further inspect the expression patterns of these driver features, we generated heatmaps of the model’s denoised data for the top 25 features per factor. The data in these heatmaps was scaled per feature (scale = "row") to highlight relative changes across samples. To systematically annotate the biological meaning of each factor, we performed GSEA on the full, ranked lists of feature weights. For each factor and each view, features were separated into those with positive and negative weights, and these two sets were tested for enrichment independently. GO Enrichment (MSigDB) enrichment was performed against the GO gene sets from the MSigDB C5 collection using the run_enrichment function within the MOFA2 package Pathway and Phenotype Enrichment (g:Profiler) comprehensive enrichment was performed using the gprofiler2 package. This analysis tested for enrichment against the Reactome (REAC), Human Phenotype Ontology (HP), and Gene Ontology (GO:BP, GO:CC, GO:MF) databases. For all enrichment analyses, pathways with p.adj < 0.05 were considered statistically significant.

### Protein Isolation & Western Blot

For total protein isolation and Western blot analysis, cells were lysed as previously described in Pull-Down section. Protein concentrations were determined using the Bicinchoninic Acid (BCA) Assay (ThermoFisher #23225). Samples were then denatured by boiling in 4x Laemmli Sample Buffer (Bio-Rad #1610747) containing 10% ß-mercaptoethanol at 95°C for 5 min. Denatured samples were loaded onto 4-12% Bis-Tris SDS-polyacrylamide gels (ThermoFisher #NP0302). Electrophoresis was performed using MES SDS running buffer (ThermoFisher #NP0002) at 120V for 1.5 h. Subsequently, proteins were transferred to PVDF membranes (ThermoFisher #88518) using iBlot^TM^ Dry Blotting System standard program (ThermoFisher #IB21001), consisting of a three-step voltage gradient: 20 V for 1 min, followed by 23 V for 4 min, and 5 V for 2 min, for a total transfer time of 7 minutes. For total protein normalization, membranes were rinsed with ultrapure water and stained with Revert™ 700 Total Protein Stain (LI-COR #926-11010) for 5 min before the blocking step. Membranes were washed twice with Revert™ Wash Solution (LI-COR #926-11012) and imaged in the 700 nm channel using an Odyssey Imaging System (LI-COR Biosciences). Following total protein imaging, membranes were rinsed with PBST and blocked with 5% Milkpowder (blotting grade, Carl Roth #T145) in PBST for 1 hr at RT and incubated with following primary antibodies overnight at 4°C, anti-FAK (1:1000, Abcam #ab40794) and anti-pFAK (Tyr397; 1:2000 Cell Signaling Technology #8556. After washing three times with PBST, membranes were incubated with HRP-conjugated anti-rabbit or anti-mouse secondary antibodies (1:5000, Bio-Rad #1706515 or #1706516) for 1 hr at RT. Chemiluminescent signals were detected using Clarity™ Western ECL Substrate (Bio-Rad #170-5060) and captured with a ChemoStar Touch ECL & Fluorescence Imager. Target band intensities were normalized to the total protein signal of the respective lane using Fiji (ImageJ) according to Revert™ 700 Total Protein Stain (LI-COR #926-11010) protocol.

### Incucyte Imaging for Proliferation Assay

For live-cell imaging, hiPSC-derived neurons were seeded on d8 of differentiation into 96-well ImageLock plates (Sartorius #4379) at a density of 50,000 cells per well. The plates were pre-coated with Poly-L-Ornithine followed by Geltrex. On d9, the cells were stained to track proliferation. IncuCyte® NucLight Rapid Red Dye (Sartorius #4717) was diluted 1:1500 in fresh neuronal differentiation medium and added to each well. The plate was incubated for 1 h and then immediately placed inside the IncuCyte® S3 instrument. The plate was monitored continuously for a total of 7 days inside the IncuCyte® S3 Live-Cell Analysis System. Phase-contrast and red fluorescence images were acquired every 2 hours using a 10x objective with 3 images were captured per well. We validated the findings in two independent neuronal differentiations and imaging runs to account for potential variability.

### Analysis of Incucyte Proliferation Assay

Time-lapse images acquired from the IncuCyte S3 were exported and processed using the density counting workflow in the machine learning toolkit Ilastik (v1.4.1). The model was trained blinded on representative images using Random Forest regression algorithm with all available features. The resulting probability maps for the "live cell" class were used to quantify the area occupied by living cells (cell density) in batch processing mode for each image at every time point to assess proliferation over time. For further analysis, the results were exported from Ilastik and imported into R Studio. To account for an initial cell settling period, the first two time points (four hours) for each well were discarded as fluorescence intensity peaked at 4h after staining. The cell density at each subsequent time point was then normalized by dividing it by the density of the third time point (the new t=0) to generate a fold-change value. The resulting growth curves, representing the mean normalized density ±SEM for each condition, were plotted over time. The subsequent time-course data was analyzed using a Linear Mixed-Effects Model (LMM), fitted with the lme4 R package.^181^ This approach was chosen to properly account for the repeated-measures nature of the longitudinal data. The raw CellDensity was modeled as the response variable. The model’s fixed effects included the interaction between genotype and time, where time (TotalHours) was modeled with both linear and quadratic terms to capture potentially non-linear growth patterns. To account for baseline variability between individual wells, each unique well was included as a random intercept. The overall significance of differences in the proliferation curves between conditions was assessed using a Type III ANOVA with Satterthwaite’s method for p-value approximation from the lmerTest package.^182^ For significant interactions, post-hoc pairwise comparisons of the estimated linear growth rates (slopes) between each mutant cell line and the WT control were performed using estimated marginal means emmeans package, using the Tukey method from the emmeans package.^183^

### Incucyte Imaging for Migration Assay

First, hiPSC-derived neurons were seeded at d8 of differentiation into 96-well ImageLock plates (Sartorius #4379) at a density of 50,000 cells per well. The plates were pre-coated with Poly-L-Ornithine and Geltrex. Cells were then cultured for four days to allow for the formation of a confluent monolayer. On d12 of differentiation, a uniform, cell-free scratch was created in the center of each well using the IncuCyte® WoundMaker tool (Sartorius, #BA-04858). Immediately following the scratch, a full medium change was performed. The fresh neuronal differentiation medium was supplemented with IncuCyte® NucLight Rapid Red Dye (Sartorius, #4717, 1:1500 dilution) to label cell nuclei. The plate was immediately placed into an IncuCyte® S3 Live-Cell Analysis System and monitored for a total of 5 days. Both phase-contrast and red fluorescence images were acquired every 2 hours using a 10x objective.

### Population-level Analysis of Incucyte Migration Assay

Time-lapse images from the IncuCyte scratch assay were processed to quantify cell migration over a 5 days. Similar to the proliferation assay, the density module from Ilastik was used to quantify the number of cells over time but in different areas of the image: The entire field of view, to measure overall cell confluence and the initial scratch area only, to specifically measure cells moving into the wound. To obtain a metric for cell migration that corrects for potential differences in cell proliferation, we calculated a "Relative Wound Density" for each time point. First, to correct for any cells or debris remaining after the scratch was made, the cell density within the scratch at t=0 was subtracted from the scratch density at all subsequent time points for each well. This background-corrected scratch density was then normalized by dividing it by the total cell density of the full image at the corresponding time point and expressing it as a percentage. The resulting time-course data, representing the mean Relative Wound Density ± SEM for each condition, were visualized as line plots. To statistically compare the migration dynamics between cell lines, we fitted a Linear Mixed-Effects Model (LMM) using the lme4 R package. This approach was chosen to properly account for the multiple measurements from the same wells over time. The calculated “Percent In Scratch” was modeled as the response variable. The model’s fixed effects included the interaction between Genotype and time (Total Hours), where time was modeled with both linear and quadratic terms to capture non-linear migration patterns. To account for baseline variability between individual wells, each well was included as a random intercept. The overall significance of differences in the migration curves was assessed using a Type III ANOVA with Satterthwaite’s method for p-value approximation (lmerTest package). This was followed by pairwise comparisons of the estimated linear migration rates (slopes) between each mutant cell line and the WT control, performed using estimated marginal means (emmeans package), using the Tukey method.

### Single-cell-level Analysis of Incucyte Migration Assay

To analyze migration dynamics at the single-cell level, the same time-lapse images used for population-level analysis were further processed. For tracking, the wound area was cropped to focus specifically on cells actively migrating into the scratch zone. Image processing and quantification were performed using Fiji (ImageJ). Time-lapse image stacks were generated from frames captured every 2 hours at 10x magnification. Single cells with IncuCyte® NucLight Rapid Red Dye (Sartorius, #4717, 1:1500 dilution) staining were tracked using the TrackMate plugin.^217^ Within TrackMate, a Laplacian of Gaussian (LoG) filter was used for spot detection, followed by the Linear Assignment Problem (LAP) Tracker. The tracking parameters were set to a maximum frame-to-frame linking distance of 60 pixels. Track segment gap closing was permitted with a maximum distance of 100 pixels and a maximum gap of 2 frames. The resulting trajectories were exported to ICY software^218^ for the quantification of average cell speed and average total displacement. Finally, motility data and spatial coordinates were imported into R and used to visualize individual cell trajectories. For statistical analysis, we used a linear mixed-effects model with genotype as a fixed effect and well as a random intercept, thereby treating wells as the replicates and accounting for the non-independence of multiple tracked cells measured within the same well. WT was specified as the reference group, and each mutant line was compared directly with WT using model-based contrasts.

### Spreading assay

For the cell spreading assay, hiPSC-derived neurons at d8 of differentiation were detached using Accutase (ThermoFisher #A1110501) and seeded at a density of 50,000 cells into 96-well ImageLock plates pre-coated with Poly-L-Ornithine (Sigma-Aldrich #P3655) followed by Geltrex (ThermoFisher #A1413302) per well. Cells were allowed to adhere and spread for 120 minutes inside the incubator at 37°C. Following this incubation, the medium was immediately removed, and cells were washed with one time with PBS to remove dead and unattached cells and fixed with 4% paraformaldehyde (PFA; Electron Microscopy Sciences #15710) for 10 minutes at room temperature to capture the spreading morphology. Following fixation, cells were washed twice with PBS and permeabilized with 0.5% Triton-X-100 (Sigma-Aldrich #9036-19-5) for 10 minutes. Fixed cells were blocked for 1 hour using a blocking buffer containing 3% BSA (Roth #8076.2). To visualize the cytoskeleton and nuclei for spreading quantification, cells were stained with Alexa Fluor^TM^ 647 Phalloidin (ThermoFisher #A22287) for F-actin and DAPI (ThermoFisher #D1306) for 30 minutes. After staining, cells were washed three times with PBS and mounted using fluoromount-G^TM^ (Thermofisher #00-4958-02). Images were automatically acquired using an Olympus ScanR high-content screening system based on an IX81 inverted microscope platform, equipped with a Hamamatsu Orca Flash 4.0 camera. For each well, nine fields of view were captured using a UPLSAPO 20x/0.75 Air. To quantify the cell areas, the raw image segmentation was performed by uploading the images to a Google Colab cloud environment and utilizing a specialized 2D prediction notebook Cellpose (v2.3.2).^169^ Individual cell boundaries were defined using the ’cyto2’ pre-trained model, with using the F-actin staining as the primary morphological source. The resulting 16-bit label maps were imported into Fiji (ImageJ) for ROI-based analysis. To convert the label maps into individual Regions of Interest (ROIs), the SCF plugin (SCF -> Segmentation -> LabelMap to ROI Manager 2D) was utilized. The total cell surface area was calculated for each ROI, and single, non-overlapping cells were isolated using size-thresholding to exclude debris or cell clusters. Significance was determined by one-way ANOVA with a Welch’s t-test between each mutant line and the WT control.

### Flow Cytometry Proliferation Staining in hiPSC-derived Neurons

We measured the proliferative capacity of the neuronal lines using a fluorescence-based dye dilution assay. The procedure was initiated on d8 of differentiation, during the routine passaging step. Neuronal cultures were first dissociated into a single-cell suspension by incubating them with Accutase for 45 min at 37°C. The cells were then pelleted at 200g for 5 minutes, counted and then resuspended in fresh neuronal differentiation medium. For the staining, this single-cell suspension was incubated with a 1:1000 dilution of a green-fluorescent cytoplasmic dye (CytoPainter, Abcam #ab138891) for 20 min at 37°C. After staining, the cells were pelleted by centrifugation to remove excess dye and resuspended in fresh medium. At this point, a sample aliquot was taken for immediate analysis to establish the baseline fluorescence (t=0) of the undivided population. The remaining cells were re-plated for continued culture. The cells were harvested on d16 to measure the dilution of the fluorescent signal (t=1) as described previously^219^. An FC washing buffer was prepared by combining 30 mL of HBSS (Thermo Fisher Scientifc, #14025092) with 150 µL of 1 M MgCl₂ (AppliChem #AP131396.1210), 6 µL of Thiazovivin, and 1.5 mL of reconstituted DNase I (Worthington Biochemical, #LK003172; each vial diluted in 500 µL HBSS). The neuron dissociation solution was prepared by mixing 5 mL of Accutase (Thermo Fisher Scientific #A1110501) with 5 mL of HBSS containing two vials of reconstituted papain (Worthington Biochemical #LK003178), supplemented with 3 µL of Thiazovivin and 500 µL of reconstituted DNase I. In preparation, the centrifuge was pre-cooled to 4°C. The HBSS and neuron dissociation solution were warmed to 37°C for 5-10 minutes. Culture medium was removed from neurons cultured in 6-well plates, and the cells were washed once with the warmed HBSS. 400 µL of the warmed dissociation solution was added to each well, and the plates were incubated for 20 min at 37°C. The reaction was stopped by adding 400 µL of FC wash buffer, causing the neurons to lift off as a single sheet. This sheet was transferred to a microcentrifuge tube and vigorously triturated with a P1000 pipette, then centrifuged at 2300 RPM for 7 minutes. After discarding the supernatant, the pellet was resuspended in 150 µL of FC wash buffer and further triturated with a 1 mL pipette tip to create a single-cell suspension. For both the d8 and d16 time points, the final cell pellets were resuspended in the FC washing buffer, kept on ice, and analyzed on the BD FACSymphony™ A1 Cell Analyzer.

### Flow Cytometry Proliferation Data Processing and Analysis

Single-cell fluorescence data was exported from FlowJo (v10) after initial gating on live (FSC vs. SSC) and single cells (FSC-A vs. FSC-H). The resulting .csv files were imported into R, and metadata for cell line, replicate, and time point were extracted from the filenames. All fluorescence intensity values for the CytoPainter dye were transformed using the inverse hyperbolic sine (arsinh)function with a cofactor of 150 to normalize the data distribution. To quantify the overall intensity of the CytoPainter dye, the Median Fluorescence Intensity (MFI) was calculated from the arsinh-transformed data for each biological replicate. These replicate MFI values were visualized using box plots to compare expression levels across all cell lines. To determine statistical significance, the MFI of each mutant cell line was compared to the WT control using a one-way ANOVA with a Welch’s t-test between each mutant line and the WT control.

### Immunofluorescence

For immunofluorescence analysis, hiPSC-derived neurons cultured on 96-well imaging plates (Ibidi, #89606) and fixed on d16 and d30 of differentiation. The culture medium was removed, and cells were washed once with PBS supplemented with Ca²⁺ and Mg²⁺ (ThermoFisher Scientific, #14040117). Fixation was performed by incubating the cells in 4% paraformaldehyde (PFA; Electron Microscopy Sciences #15710) for 10 min at room temperature. Following fixation, cells were washed twice with PBS. For permeabilization, cells were treated with 0.5% Triton-X-100 (Sigma-Aldrich, #9036-19-5) in PBS for 10 minutes. To quench autofluorescence from the aldehyde fixative, cells were subsequently incubated in a solution of ammonium chloride (Sigma Aldrich, #A9434-500G) and glycine (Sigma Aldrich #33226) for 5 min. After washing, non-specific antibody binding was blocked by incubating the cells for 1 hour in a blocking buffer consisting of 2% Bovine Serum Albumin (BSA; Roth #8076.2), 4% Donkey Serum (Abcam, #ab7475), and 4% Goat Serum (Sigma Aldrich #G9023) in PBS. Cells were then incubated overnight at 4°C with the following primary antibodies diluted in blocking buffer: pCREB (1:500, Cell Signaling Technologies #9198), CREB (1:1000, Cell Signaling Technologies #9104), BSN (1:200, Abcam #ab82958), PSD95 (1:500, Abcam #ab18258), pFAK (1:200, Cell Signaling Technologies #8556), FAK (1:400, Proteintech #12636-1-AP), MAP2 (1:500, Synaptic Systems #188 004). The following day, cells were washed three times for 5 min each with PBS containing 0.05% Tween-20 (Roth #9127.1) Secondary antibody incubation was performed for 1 hour at room temperature in the dark using fluorophore-conjugated secondary antibodies, diluted 1:500 in blocking buffer (Goat anti-Guinea Pig (Thermo Fisher #A-21450), Donkey anti-Rabbit (Abcam #ab150075), Donkey anti-Mouse (Thermo Fisher #A-21203), Goat anti-Rabbit (Thermo Fisher #A-11008) and Goat anti-rabbit (ThermoFisher #A-11012)) For the phalloidin staining, Phalloidin Labeling Probe (1:400, Thermo Fisher #A22287) was added during the secondary antibody incubation. Finally, cells were washed three times for 10 min each with 0.05% Tween-20 in PBS. During the second wash, nuclei were counterstained with DAPI (1:1000, Sigma-Aldrich, #D9542). The final wash was performed with PBS without detergent. The plates were sealed and stored at 4°C in the dark until imaging. Images were automatically acquired using an Olympus ScanR high-content screening system on an IX81 inverted microscope platform, equipped with a Hamamatsu Orca Flash 4.0 camera. At least 12 images per well were captured using a UPLSAPO 20x/0.75 Air (pCREB/CREB, no z-stacks) or UPLSAPO 40x/0.95 Air (FAK/pFAK and MAP2/BSN/PSD95, with z-stacks) objective lens, and the system was controlled via the Olympus ScanR (v2.5.0) software. Representative images for MAP2 and pCREB/CREB figures were caputured using UPLSAPO 20x/0.75 Air objective lens and gamma adjusted equally across images for better visibility.

### Image Analysis

Deconvolution in Huygens Professional (v25.04) was performed using the Classic Maximum Likelihood Estimation algorithm in the workflow processing mode. The resulting 3D deconvolved stacks were exported and then imported into FIJI (v2.16.0), where Maximum Intensity Projections (MIPs) were generated for subsequent 2D analysis in CellProfiler. Images stained for pCREB/CREB were analyzed directly from the 2D acquisitions without deconvolution. Quantitative analysis of 2D images for pCREB/CREB was performed using CellProfiler (v4.2.8). First, nuclei were segmented from the DAPI channel using the IdentifyPrimaryObjects module. A quality control step was then applied using the FilterObjects module to remove potential artifacts (e.g., out-of-focus or over-saturated nuclei). Using the MeasureObjectIntensity module, the Mean Intensity for both the pCREB and CREB channels were measured within each "NucleiFiltered" object. Finally, the ExportToSpreadsheet module was used to export the per-nucleus measurements. Exported CSV files were imported into R for visualization and downstream statistical analysis. The MIPs of the four-channel (DAPI, MAP2, BSN, PSD-95) were imported into CellProfiler. Nuclei were segmented from the DAPI channel using the IdentifyPrimaryObjects module with an Otsu global threshold and then filtered to exclude segmentation artifacts and debris. For neurite morphology, the MAP2 channel was thresholded using a Minimum Cross-Entropy method to generate a binary mask of the dendritic arbor. This mask was processed using the MorphologicalSkeleton module to calculate the total neurite length and number of branches per image (MeasureImageSkeleton), while total dendritic area was quantified using MeasureImageAreaOccupied. Synaptic puncta (BSN and PSD-95) were identified using a "Speckle" enhancement filter followed by segmentation using a Minimum Cross-Entropy threshold and then filtered to exclude artifacts. To restrict analysis to dendritically localized synapses, only puncta overlapping with the MAP2-positive binary mask were retained (MaskObjects). Functional synapses were defined as the physical overlap between presynaptic (BSN) and postsynaptic (PSD-95) puncta. This was quantified using the RelateObjects module, establishing a parent-child relationship between BSN and PSD-95 objects; a synapse was defined as a BSN punctum containing at least one PSD-95 punctum (and vice-versa). The resulting quantification were exported and processed in R. Data were aggregated at the well level (biological replicate) by summing the values from all images within a well (n = 12 images per well; 6 wells per genotype). Metrics were normalized to assess distinct biological features: Total neurite length, branch count, and total synaptic puncta counts were divided by the total number of nuclei in the well to assess the total output per cell. Synaptic puncta counts were divided by the total neurite length (in µm) of the well to quantify the density per µm neurite length. Mean puncta area and mean integrated intensity were calculated by averaging all objects within a well. Colocalization ratios were calculated as the percentage of BSN puncta positive for PSD-95 and the percentage of PSD-95 puncta positive for BSN. Statistical Analysis Statistical comparisons were performed on the aggregated well-level data. Differences between each mutant cell line and the isogenic WT control were assessed using Welch’s t-test (two-sided), to account for unequal variances between groups. P-values < 0.05 were considered statistically significant.

### Potassium chloride (KCl)-induced depolarization

For the KCl-induced depolarization, hiPSC-derived neurons cultured on 96-well imaging plates (Ibidi, #89606). On d16 and d30 of differentiation, the neuronal maturation medium was fully removed and replaced with fresh neuronal maturation medium supplemented with 50mM KCl (Sigma-Aldrich, #P5405). After 60min of incubation at 37°C, cells were fixed for immunofluorescence applications (see section below).

### Flow Cytometry Staining of Surface Marker Expression

For the analysis of surface protein expression, hiPSC-derived neurons on d30 of differentiation were dissociated and stained. All solutions were prepared immediately prior to the procedure. An FC washing buffer was prepared by combining 30 mL of HBSS (Thermo Fisher Scientific #14025092) with 150 µL of 1 M MgCl₂ (AppliChem #AP131396.1210), 6 µL of Thiazovivin, and 1.5 mL of reconstituted DNase I (Worthington Biochemical #LK003172; each vial diluted in 500 µL HBSS). The neuron dissociation solution was prepared by mixing 5 mL of Accutase (Thermo Fisher Scientific #A1110501) with 5 mL of HBSS containing two vials of reconstituted papain (Worthington Biochemical #LK003178), supplemented with 3 µL of Thiazovivin and 500 µL of reconstituted DNase I. An FC blocking buffer was prepared by diluting 530 µL of 7.5% BSA Fraction V (ThermoFisher Scientific #15260037) with 3.5 mL of the FACS washing buffer. In preparation, the centrifuge was pre-cooled to 4°C.

The HBSS and neuron dissociation solution were warmed to 37°C for 5-10 min. Spent culture medium was removed from neurons cultured in 6-well plates, and the cells were washed once with the warmed HBSS. 400 µL of the warmed dissociation solution was added to each well, and the plates were incubated for 20 minutes at 37°C. The reaction was stopped by adding 400 µL of FC wash buffer, causing the neurons to lift off as a single sheet. This sheet was transferred to a microcentrifuge tube and vigorously triturated with a P1000 pipette, then centrifuged at 2300 RPM for 7 min. After discarding the supernatant, the pellet was resuspended in 150 µL of FC wash buffer and further triturated with a 1 mL pipette tip to create a single-cell suspension. The entire suspension was transferred to a 96-well plate and centrifuged at 300 x g for 5 min at 4°C. The supernatant was discarded, and the cells were incubated in 100 µL of blocking buffer for 25 min at room temperature. The plate was again centrifuged at 300g for 5 min, and the supernatant was aspirated. The cell pellets were then resuspended in 50 µL of blocking buffer containing a cocktail of the following fluorescence-conjugated primary antibodies: anti-CDH2 (N-Cadherin) PerCP-Cy™5.5 (1:15 dilution, BioLegend #350813), anti-THY1 (CD90) PE (1:15 dilution, Ctag Lab #PE-FLA98128), anti-ITGB1 (CD29) Alexa Fluor® 647 (1:15 dilution, R&D Systems #FAB17781R) and anti-ITGA3 (CD49c) APC-Vio®770 (1:8 dilution, Miltenyi Biotec #130-105-409). Staining proceeded for 45 min on ice in the dark. Following incubation, cells were pelleted (300 x g, 5 min), washed once with FC wash buffer, and finally resuspended in wash buffer. The samples were transferred to fresh microcentrifuge tubes and kept on ice. Immediately before analysis, each sample was filtered through a 70 µm FC filter tube. Data were acquired on the BD FACSymphony™ A1 Cell Analyzer.

### Flow Cytometry Surfacer Marker Data Processing and Analysis

The initial gating strategy in FlowJo (v10) isolated for live and single cells which constituted the starting population for all subsequent computational analysis. Single-cell data for four surface markers (CDH2, THY1, ITGB1, and ITGA3) were exported from FlowJo (v10) as individual .csv files for each sample. Raw fluorescence intensity values from the compensated channels were imported into R. To manage the wide dynamic range of fluorescence data and stabilize variance, all intensity values were transformed using the inverse hyperbolic sine (arsinh) transformation with a cofactor of 150. As an initial quality control step, the total number of cells per sample was quantified, and the Pearson correlation between the four markers was calculated and visualized. To visualize the high-dimensional relationships between all cells, a Uniform Manifold Approximation and Projection (UMAP) was performed. For this, a representative dataset was created by randomly subsampling 2,000 cells from each sample. The UMAP algorithm from the uwot package^185^ was run on the combined, subsampled data, and the resulting two-dimensional embedding was used to visualize the single-cell landscape, with cells colored by either genotype or individual marker expression. For quantitative comparisons between cell lines, the Median Fluorescence Intensity (MFI) was calculated for each of the three biological replicates per line and were visualized using box plots. To quantify the proportion and intensity of marker-positive cells, a gating strategy was applied. Signal intensity thresholds for positivity were established for each marker on the arcsinh-transformed scale based on dedicated Fluorescence Minus One (FMO) controls. Using these gates, the percentage of positive cells were calculated and visualized. To determine the statistical significance of changes relative to the control, replicate-level percentage of positive cells and MFI values for each mutant cell line were compared to the WT line using a one-way ANOVA with a Welch’s t-test between each mutant line and the WT control.

## SUPPLEMENTAL INFORMATION

Supplemental Tables 1-25 are available via https://data.mendeley.com/preview/w5ktr5kj52?a=0e8bdfb9-15eb-44c1-9489-e3fa41ef6b37

---

**Supplementary Figure 1.**
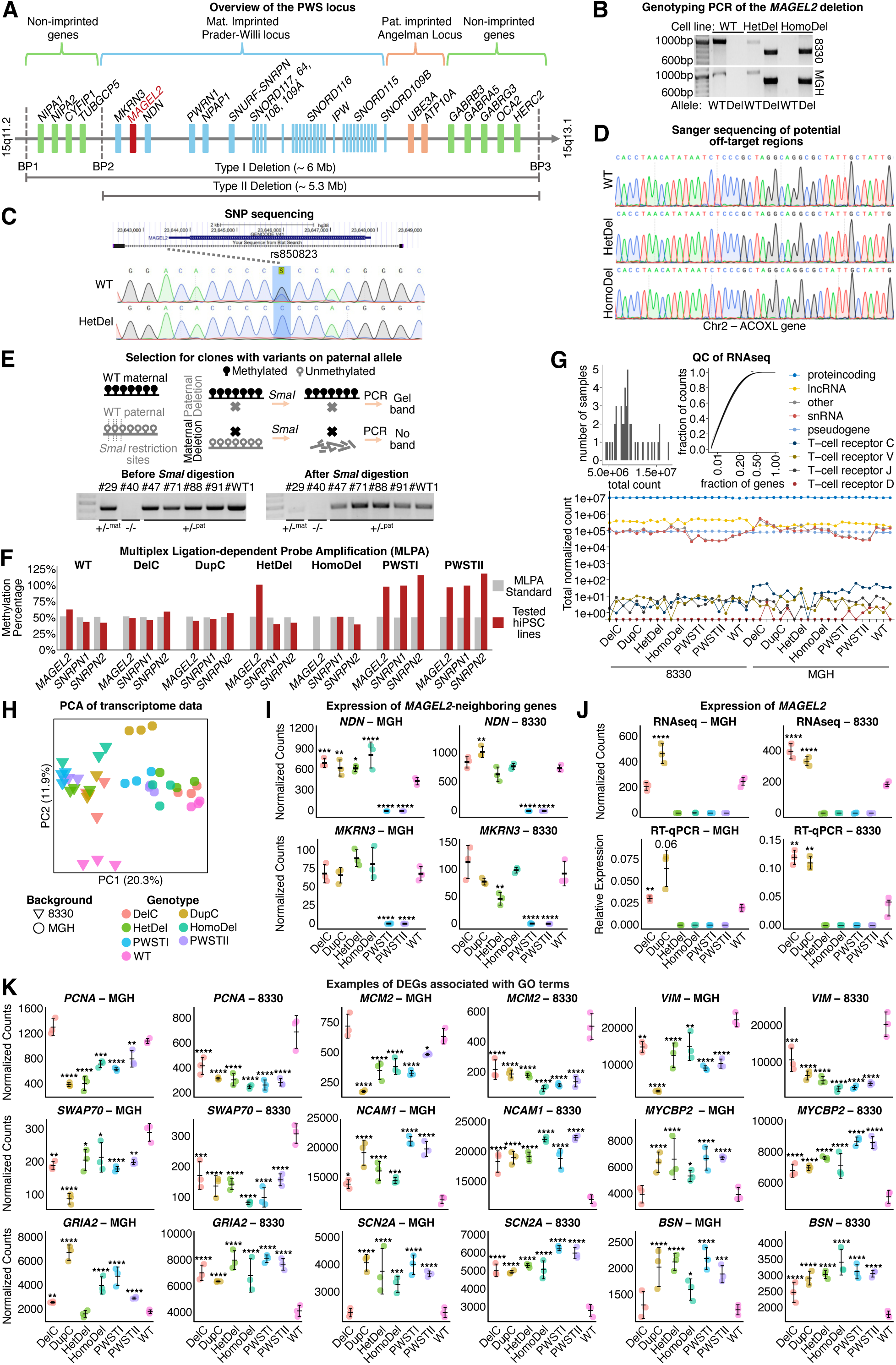
: Transcriptomic consequences of CRISPR/Cas9 engineered variants modeling SYS and PWS on cortical neurons. (A) Scheme of the PWS locus highlighting the *MAGEL2* gene (red). (B-D) Validation of the targeting. (B) Examples of genotyping PCRs using primers flanking the CRISPR/Cas9 deleted region in HetDel and HomoDel hiPSC lines. (C) UCSC genome browser alignment and Sanger sequencing of heterozygous SNPs rs850823 (G>A/C/T) located within the CRISPR/Cas9 deleted region confirm HetDel clonality. (D) Exemplified potential off-target region (Chr2 – ACOXL gene) with no detectable off-target edits. (E) SmaI digestion scheme to determine deleted allele in HetDel hiPSC clones with PCR results pre- and post-digestion. HomoDel clone #40 serves as a negative control. Clone #29 shows maternal deletion; clones #47, #71, #88, and #91 show paternal deletions. (F) MLPA confirms expected imprinted pattern consistent with the CRISPR/Cas9 targeting. (G) Comparable library size, complexity, and gene biotype distribution across all samples. (H) PCA of the transcriptomic data. Samples are colored by genotype and shaped by genetic background (n=3). (I) RNA-seq expression of *MAGEL2*-neigbouring genes *NDN* and *MKRN3* (schematically indicated in A). (J) *MAGEL2* expression from RNA-seq (Bonferroni-adjusted p-values) and qPCR results (n=3, ANOVA with pairwise Welch’s t-tests vs. WT). (K) Expression levels of selected DEGs enriched in GO terms across genotypes. In all quantifications, error bars = SD, (*p < 0.05, **p < 0.01, ***p < 0.001, and ****p < 0.0001).

**Supplementary Figure 2.**
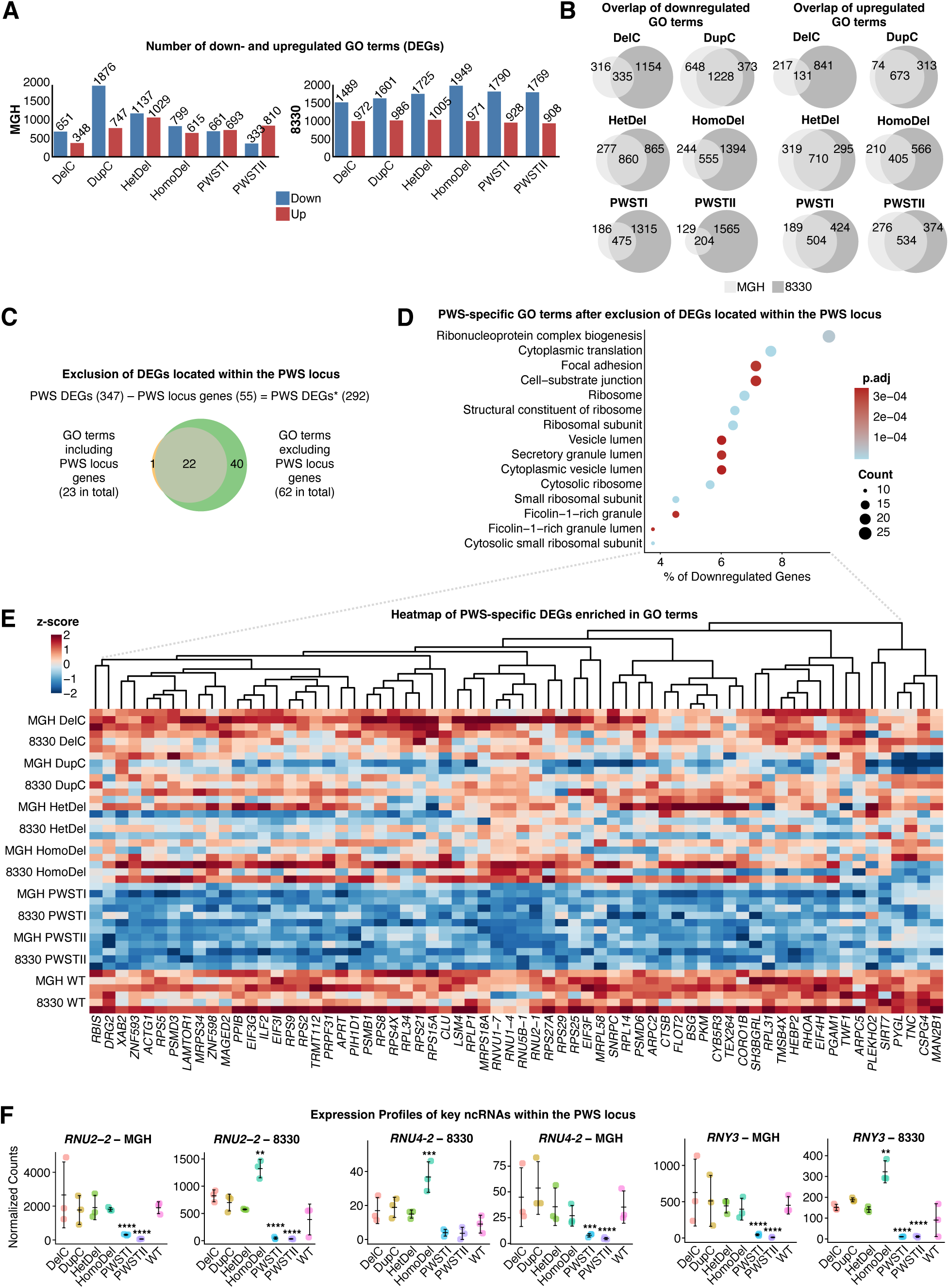
: PWS-specific GO terms are enriched in genes associated with ribosomal biology. (A) Number of enriched GO terms for down- and upregulated DEGs. (B) Venn diagram showing background overlapping DEG GO terms for down and upregulated DEGs with significance of overlap determined by a hypergeometric test. (C) Venn diagram of background-overlapping enriched GO terms for downregulated genes in neurons with PWS-deletion, including (left) and excluding (right) DEGs within the PWS locus. (D) Dot-plot of remaining PWS-specific GO terms after excluding DEGs of the PWS locus. (E) Heatmap of PWS-specific DEGs from enriched GO terms in (D). (F) Expression of key ncRNAs. Normalized counts with Bonferroni adjusted p-values. In all quantifications, error bars = SD, (*p < 0.05, **p < 0.01, ***p < 0.001, and ****p < 0.0001).

**Supplementary Figure 3.**
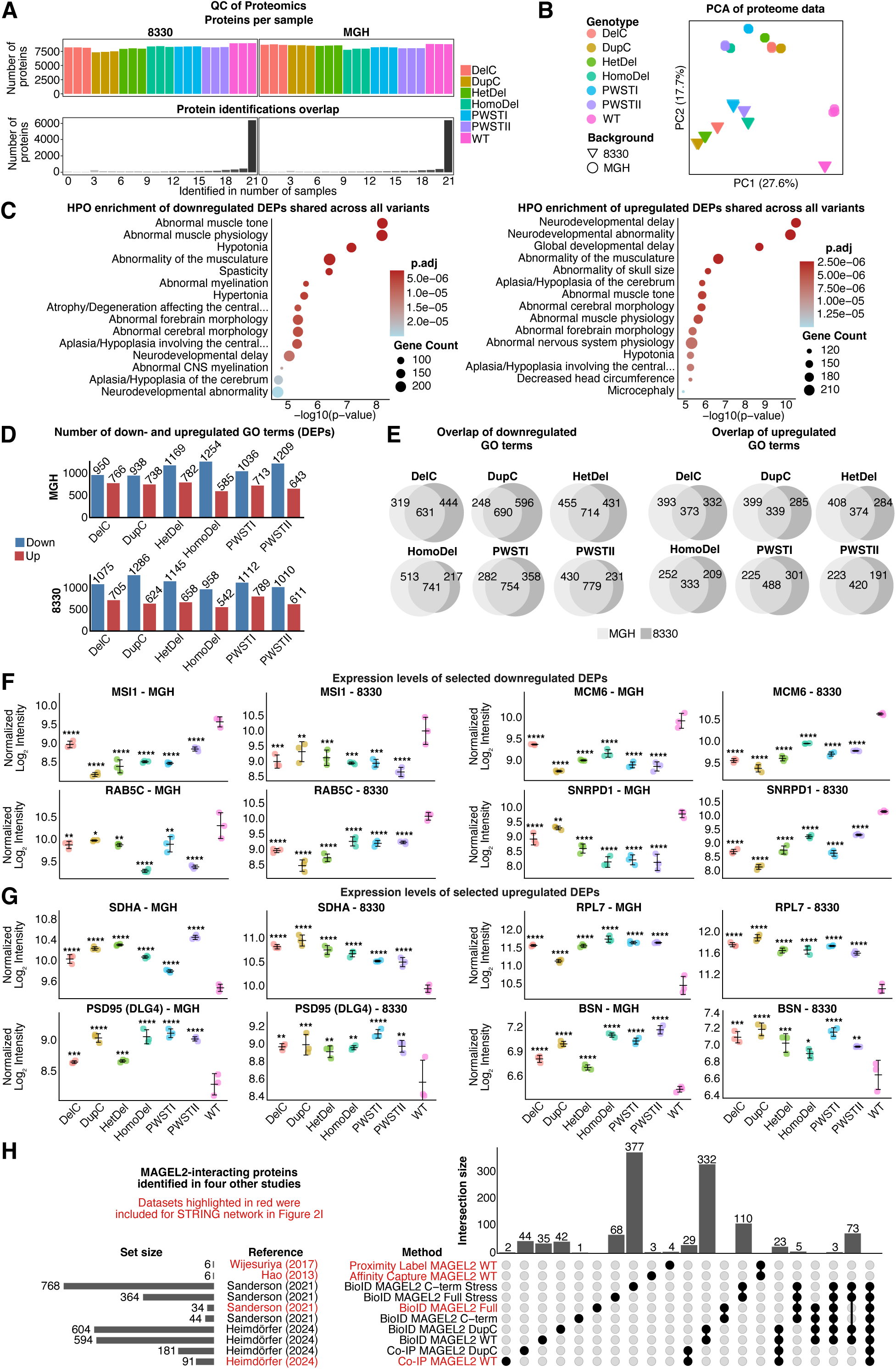
: Analysis of the proteomic data integrating GO, HPO and interactor network analysis. (A) Quality control plots for the proteome data, showing the number of identified proteins per sample (top) and number of overlapping identified proteins across samples (bottom). (B) PCA of the proteomic data. Samples are colored by genotype and shaped by genetic background (n=3). (C) HPO enrichment for the down- and upregulated proteins overlapping across all genotypes and backgrounds. (D) Number of up- and downregulated enriched GO terms for DEPs. (E) Venn diagram showing background overlapping DEP GO terms for down and upregulated DEPs with significance of overlap determined by a hypergeometric test. (F-G) Expression of selected downregulated (F) and upregulated (G) proteins with significance determined using Bonferroni adjusted p-values of the primary differential expression analysis. (H) UpSet plot integrating MAGEL2-interacting proteins identified from four publications. In all quantifications, error bars represent SD, and asterisks denote statistical significance (*p < 0.05, **p < 0.01, ***p < 0.001, and ****p < 0.0001).

**Supplementary Figure 4:**
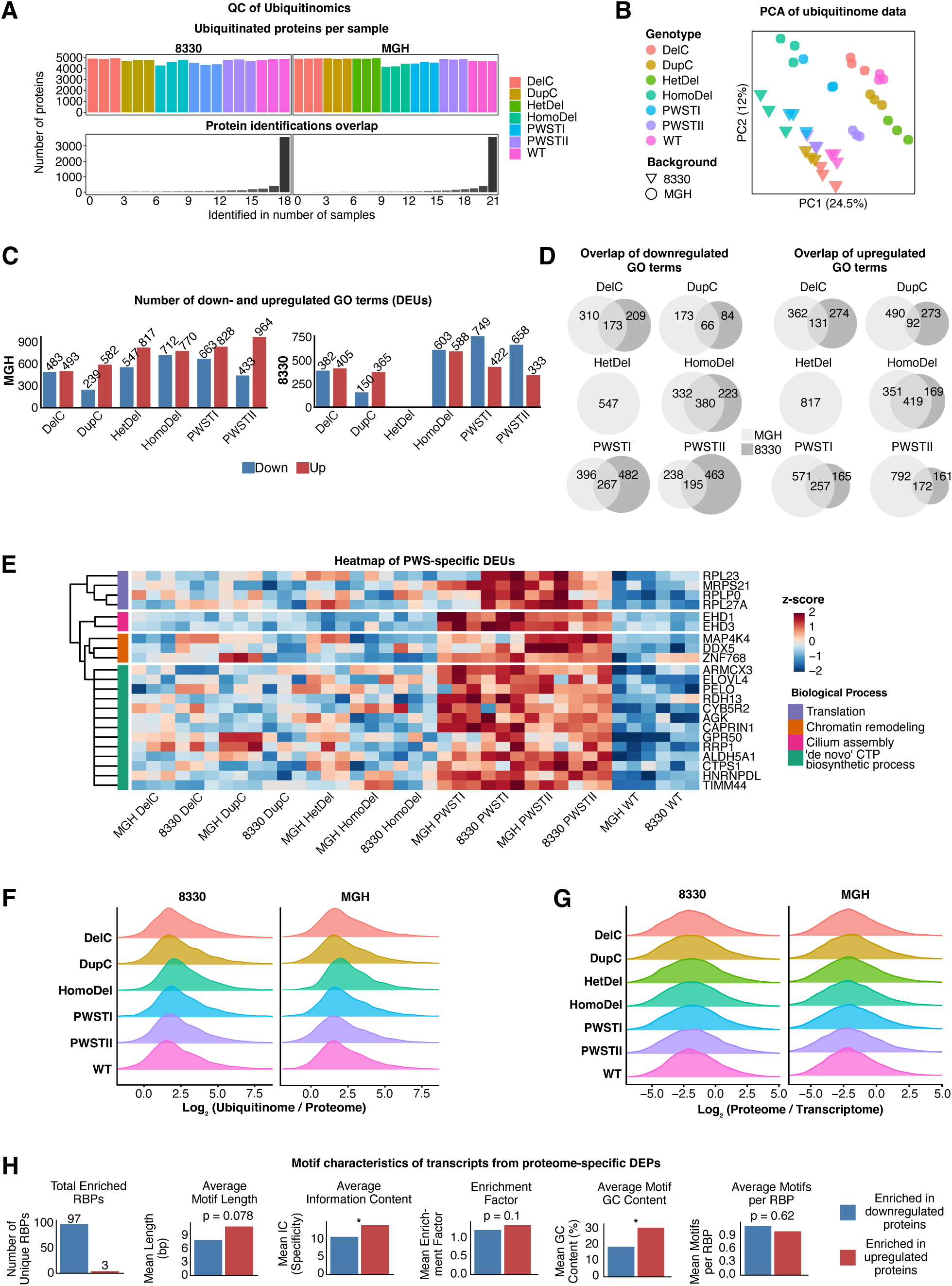
High quality of ubiquitinome data and integration of different omics layers. (A) Quality control plots for the ubiquitinome data, showing the number of identified proteins per sample (top) and number of overlapping identified proteins across samples (bottom). (B) PCA of the ubiquitinomic data. Samples are colored by genotype and shaped by genetic background (n=3). (C) Number of enriched GO terms for up- and downregulated DEUs. (D) Venn diagram showing background overlapping DEU GO terms for down and upregulated DEUs with significance of overlap determined by a hypergeometric test. (E) Heatmap of all PWS-specific upregulated DEUs. (F) Ridgeline plot of the log2-transformed ratio of ubiquitinated protein to total protein abundance (Ubiquitinome / Proteome). (G) Ridgeline plot of the log2-transformed ratio of protein to transcript abundance (Proteome / Transcriptome). (H) Bar plots comparing transcript motif characteristics between proteome-specific upregulated and downregulated proteins. Significance was determined using a Wilcoxon rank-sum test. In all quantifications, asterisks denote statistical significance (*p < 0.05, **p < 0.01, ***p < 0.001, and ****p < 0.0001).

**Supplementary Figure 5.**
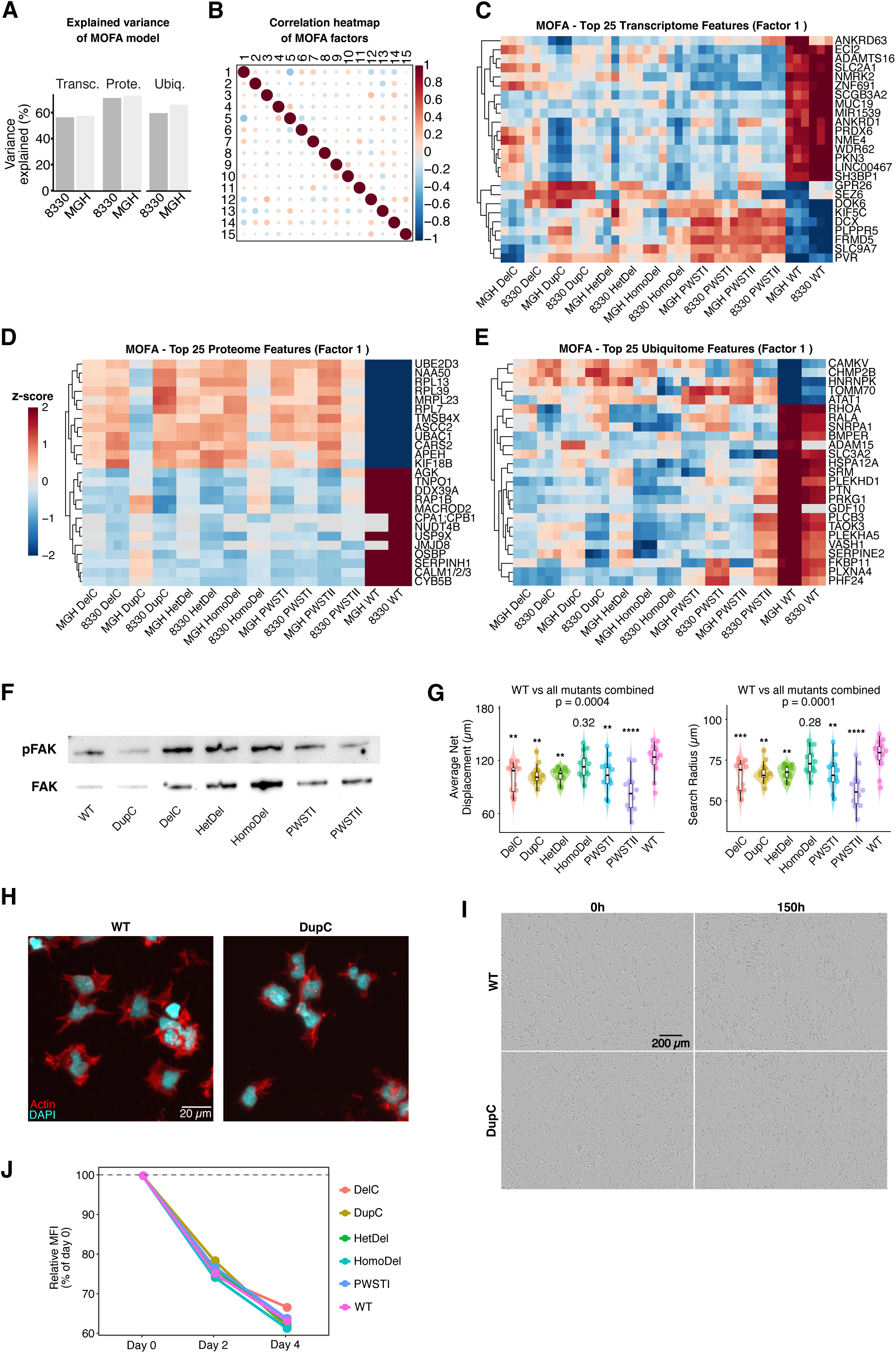
: MOFA-analysis and multi-omics validation (A) Explained variance of MOFA model in the three omics layers. (B) Correlation heatmap of the MOFA factors, demonstrating their general independence. (C-E) Heatmaps showing the z-scores of the top 25 features driving Factor 1 for the transcriptome (C), proteome (D), and ubiquitinome (E). (F) Representative Western blots of pFAK and FAK on d30 of differentiation. (G) Quantification of average net displacement and search radius from the live-cell imaging single-cell migration analysis (n=12), with significance determined by an ANOVA followed by pairwise Welch’s t-tests vs. WT. (H) Representative images from the cell spreading assay (n=3), stained for F-actin and nuclei. (I) Representative images of differentiating neurons during live-cell imaging. (J) Line graph showing the relative MFI of proliferation dye over four days in undifferentiated hiPSCs (n=1), demonstrating no difference in proliferation between genotypes. Error bars represent SEM, and asterisks denote statistical significance (*p < 0.05, **p < 0.01, ***p < 0.001, and ****p < 0.0001).

**Supplementary Figure 6.**
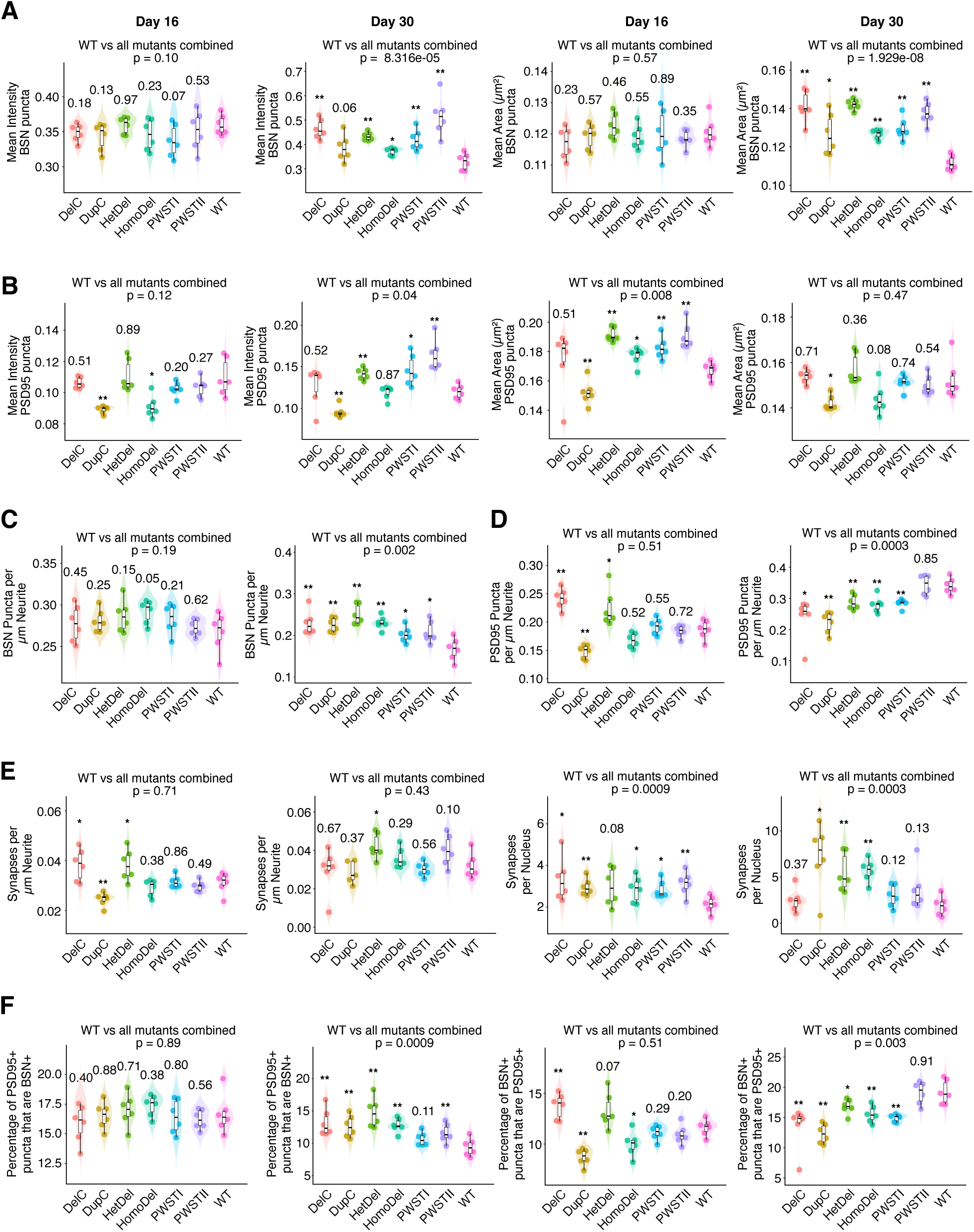
: Immunofluorescence characterizations of focal adhesions and synaptic architecture. (A-B) Quantification of mean intensity and area for BSN+ (A) and PSD-95+ puncta (B; n=6). (C-D) Number of BSN+ (C) and PSD-95+ (D) positive puncta per µm neurite (n=6). (E) Number of synapses (puncta that are both BSN+ and PSD-95+ positive) per µm neurite and nucleus (n=6). (F) Percentage of PSD-95+ puncta that are BSN+ and percentage of BSN+ puncta that are PSD-95+. For all quantifications, significance determined by an ANOVA, followed by pairwise Welch’s t-tests vs. WT. Error bars represent SEM, and asterisks denote statistical significance (*p < 0.05, **p < 0.01, ***p < 0.001, and ****p < 0.0001).

**Supplementary Figure 7.**
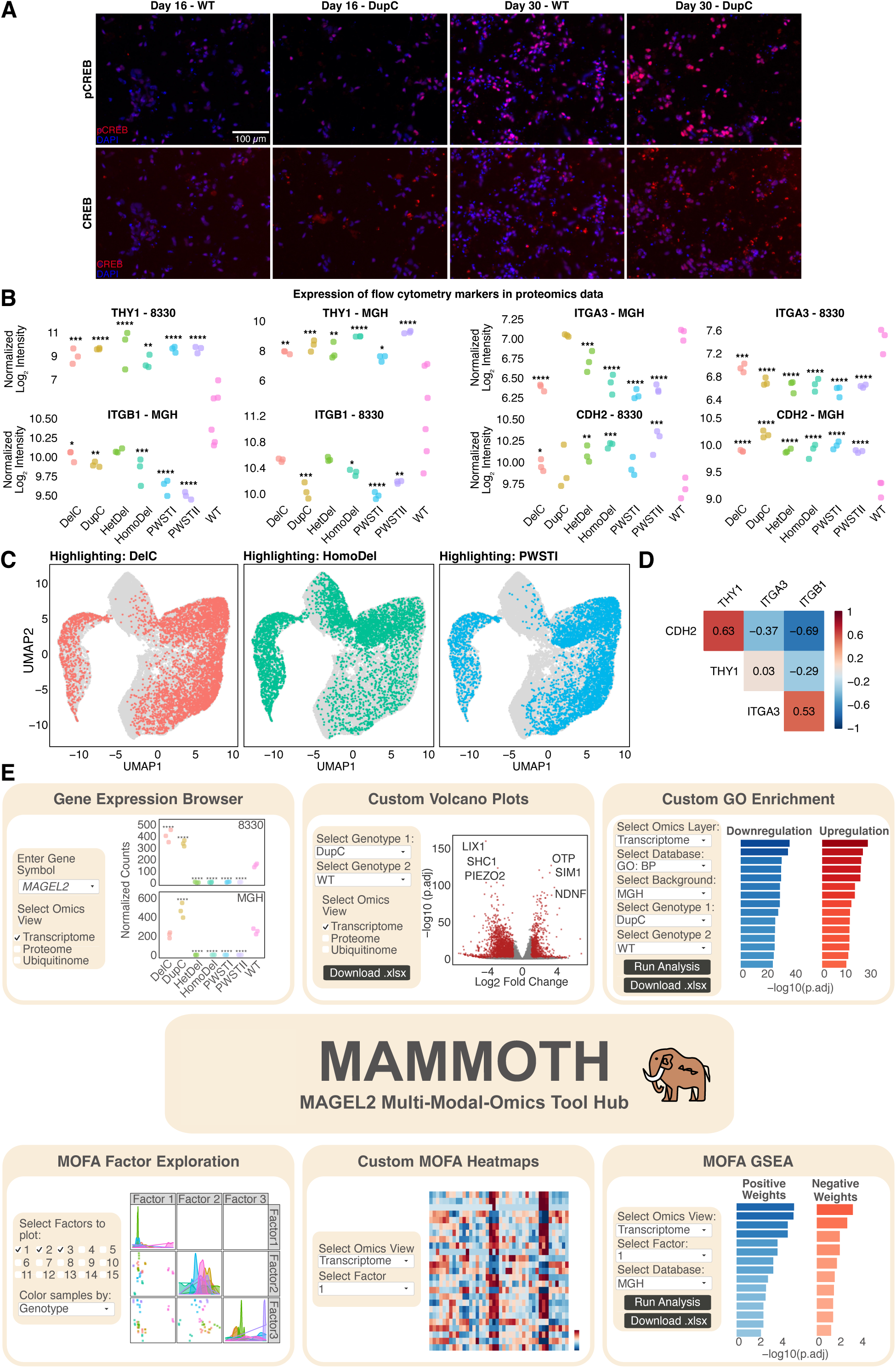
: Flow cytometry analyses and overview of the MAMMOTH data portal. (B) Representative images of IF staining for pCREB, CREB, and nuclei (DAPI) in WT and DupC neurons. (C) Expression of flow cytometry markers in proteomics data. (D) UMAP projection of single-cell data from differentiated d30 neurons, showing the distribution of cells based on genotype. (E) Correlation matrix of the flow cytometry surface markers. (F) Composite image highlighting different features of the interactive MAMMOTH web application.

## Notes

### Competing Interest Statement

The authors have declared no competing interest.

